# Cardiometabolic health and physical robustness map onto distinct patterns of brain structure and neurotransmitter systems

**DOI:** 10.1101/2024.06.14.599066

**Authors:** Eliana Nicolaisen-Sobesky, Somayeh Maleki Balajoo, Mostafa Mahdipour, Agoston Mihalik, Mahnaz Olfati, Felix Hoffstaedter, Janaina Mourao-Miranda, Masoud Tahmasian, Simon B. Eickhoff, Sarah Genon

**Affiliations:** Institute of Neuroscience and Medicine (INM-7: Brain and Behaviour), Research Centre Jülich, Jülich, Germany; Institute of Systems Neuroscience, Heinrich Heine University Düsseldorf, Düsseldorf, Germany; Department of Psychiatry, University of Cambridge, Cambridge, United Kingdom; Institute of Medical Science and Technology, Shahid Beheshti University, Tehran, Iran; UCL Hawkes Institute, Department of Computer Science, University College London, London, United Kingdom; Department of Nuclear Medicine, University Hospital and Medical Faculty, University of Cologne, Cologne, Germany

**Author notes:** Corresponding authors: **Eliana Nicolaisen-Sobesky** Institute of Neuroscience and Medicine (INM-7: Brain and Behaviour), Research Centre Jülich, Jülich, 52428, Germany, **Sarah Genon** Institute of Neuroscience and Medicine (INM-7: Brain and Behaviour), Research Centre Jülich, Jülich, 52428, Germany.

**Keywords:** Brain structure, physical robustness, cardiometabolic, risk factors, neurotransmitter systems

## Abstract

The link between brain health and risk/protective factors for non-communicable diseases (such as high blood pressure, high body mass index, diet, smoking, physical activity, etc.) is increasingly acknowledged. However, the specific effects that these factors have on brain health are still poorly understood, delaying their implementation in precision brain health. Here, we studied the multivariate relationships between risk factors for non-communicable diseases and brain structure, including cortical thickness (CT) and grey matter volume (GMV). Furthermore, we adopted a systems-level perspective to understand such relationships, by characterizing the cortical patterns (yielded in association to risk factors) with regards to brain morphological and functional features, as well as with neurotransmitter systems. Similarly, we related the pattern of risk/protective factors dimensions with a peripheral marker of inflammation.

First, we identified latent dimensions linking a broad set of risk factors for non-communicable diseases to parcel-wise CT and GMV across the whole cortex. Data was obtained from the UK Biobank (n=7370, age range=46-81 years). We used regularized canonical correlation analysis (RCCA) embedded in a machine learning framework. This approach allows us to capture inter-individual variability in a multivariate association and to assess the generalizability of the model. The brain patterns (captured in association with risk/protective factors) were characterized from a multi-level perspective, by performing correlations (spin tests) between them and different brain patterns of structure, function, and neurotransmitter systems. The association between the risk/protective factors pattern and C-reactive protein (CRP, a marker of inflammation) was examined using Spearman correlation.

We found two significant and partly replicable latent dimensions. One latent dimension linked cardiometabolic health to brain patterns of CT and GMV and was consistent across sexes. The other latent dimension linked physical robustness (including non-fat mass and strength) to patterns of CT and GMV, with the association to GMV being consistent across sexes and the association to CT appearing only in men. The CT and GMV patterns of both latent dimensions were associated to the binding potentials of several neurotransmitter systems. Finally, the cardiometabolic health dimension was correlated to CRP, while physical robustness was only very weakly associated to it.

We observed robust, multi-level and multivariate links between both cardiometabolic health and physical robustness with respect to CT, GMV, and neurotransmitter systems. Interestingly, we found that cardiometabolic health and physical robustness are associated with not only increases in CT or GMV, but also with decreases of CT or GMV in some brain regions. Our results also suggested a role for low-grade chronic inflammation in the association between cardiometabolic health and brain structural health. These findings support the relevance of adopting a holistic perspective in health, by integrating neurocognitive and physical health. Moreover, our findings contribute to the challenge to the classical conceptualization of neuropsychiatric and physical illnesses as categorical entities. In this perspective, future studies should further examine the effects of risk/protective factors on different brain regions in order to deepen our understanding of the clinical significance of such increased and decreased CT and GMV.

## Introduction

Non-communicable diseases, including cardiovascular, metabolic, mental, and neurological disorders, represent the predominant global public health challenge nowadays [1,2]. Most non-communicable diseases share a common set of risk/protective factors [3–5] (here referred simply as risk factors), like tobacco smoking, unhealthy diet, physical inactivity [1,2], excessive alcohol consumption, hypertension [2], sleep problems [6], obesity (increased body mass index (BMI)) and air pollution [2,6]. Since non-communicable diseases include mental and neurological disorders, the same set of risk factors also affects brain health [2,7].

Two of the biomarkers for brain health are cortical thickness (CT) and Grey Matter Volume (GMV). Several studies have analyzed the link between these structural markers and specific risk factors such as BMI [8–14], waist circumference [13,14], cigarette smoking [13,15,16], physical exercise [17], and diet [18]. Even though these studies have contributed to our understanding of the link between risk factors and brain structure they have some caveats. One limitation is that the reported findings have been inconsistent across studies. For instance, associations between BMI and both, global or frontal CT, have been reported as negative [8–10,13,14], positive [11–13], or not significant [9,10]. To address this point, studies using robust approaches that evaluate the generalizability of results are needed. In this regard, machine learning approaches use cross-validation to test for the generalizability of the implemented models.

A second limitation of studies linking brain structure to risk factors is that they mostly used univariate/bivariate approaches. In other words, so far studies have mainly linked specific risk factors with global measures of the brain, or with specific brain regions. [19]. However, these factors usually do not happen in isolation (e.g. BMI, diet and physical activity are strongly interrelated in population data) and hence represent a multivariate set. Accordingly, previous studies only provide a partial view of this association [15,16,19,20] impeding the discovery of distributed brain networks that are associated to a range of risk factors simultaneously [21]. Identifying distributed brain networks and their interactions, as well as their relationships with several risk factors requires “doubly” multivariate approaches. These approaches can take full advantage of the rich phenotyping included in large-scale datasets such as the UK Biobank (UKB) and embrace collinearity among risk factors variables, as well as among brain regions.

In this respect, canonical correlation analysis (CCA) is a multivariate, data-driven approach that can be used to discover large-scale distributed cortical patterns associated with several risk factors [22,23]. Of note, a regularized version of CCA (RCCA) mitigates the effect of collinearity and yields more stable results than CCA [22], improving the interpretability of results. CCA and RCCA have been previously used to search for robust and generalizable multivariate associations between different data modalities, such as brain structure, brain function, hippocampal structure, behavior, environmental variables, and psychiatric features [20,23–28]. Using these methods, numerous studies have shown that several healthy and illness-related phenotypes are linked to axes of brain structural organization [23,24,28].

Therefore, despite the link between risk factors for non-communicable diseases and brain health being acknowledged, there are still several important aspects of this association which are incompletely understood [21,29]. The lack of understanding of the relationship between risk factors and brain health prevents their use as biomarkers in the clinical practice and hence is a major research priority [21]. Since structural brain changes may be long-lasting and lead to various neuropsychiatric diseases, it is critical to pinpoint modifiable risk factors that can reduce the risk of those conditions. Understanding how risk factors for non-communicable diseases are related to brain health from a comprehensive and broad perspective requires understanding how multiple risk factors simultaneously relate to structural patterns across the whole brain. Thus, the main question of this study is how several risk factors for non-communicable diseases are associated with patterns of CT and GMV.

Another important point to consider is the multi-level nature of the interplay between risk factors and brain health. For instance, risk factors for non-communicable diseases have been associated with several brain features, including brain structure [30,31], function [32], genetics, and neurotransmitter systems [33]. This needed integration of different neurobiological features [34] can now be done quantitively with neuromaps [35], which provides access to a wide set of brain maps, including for instance genetic transcription and brain molecular features. Similarly, a role for immunoinflammatory processes in the association between risk factors and brain health has been frequently discussed [36–40]. Linking CCA-derived cortical pattern and risk factors pattern to known neurobiological patterns and inflammation will allow to gain a systems-level understanding of the association between risk factors and brain health.

Hence, to gain a comprehensive, generalizable, and multi-level understanding of the relationship between risk factors and brain health, it is needed to use robust machine learning approaches with generalizability testing, along with methods that integrate different data modalities [28,41]. With that aim, we first searched for latent dimensions linking a wide range of risk factors for non-communicable diseases to region-wise CT and GMV across the whole brain cortex using RCCA embedded in a machine learning framework (Figure 1). In other words, we searched for those combinations of risk factors which are most relevant for CT and GMV interindividual variability [20]. On a second step we aimed to characterize the captured brain patterns from a neurobiological perspective comparing them with existing brain maps spanning brain structure, function, genetic variability, and neurotransmitter systems. Finally, we analyzed the link between the risk factors dimension and a peripheral marker of inflammation: C-reactive protein (CRP).

**Figure 1.**
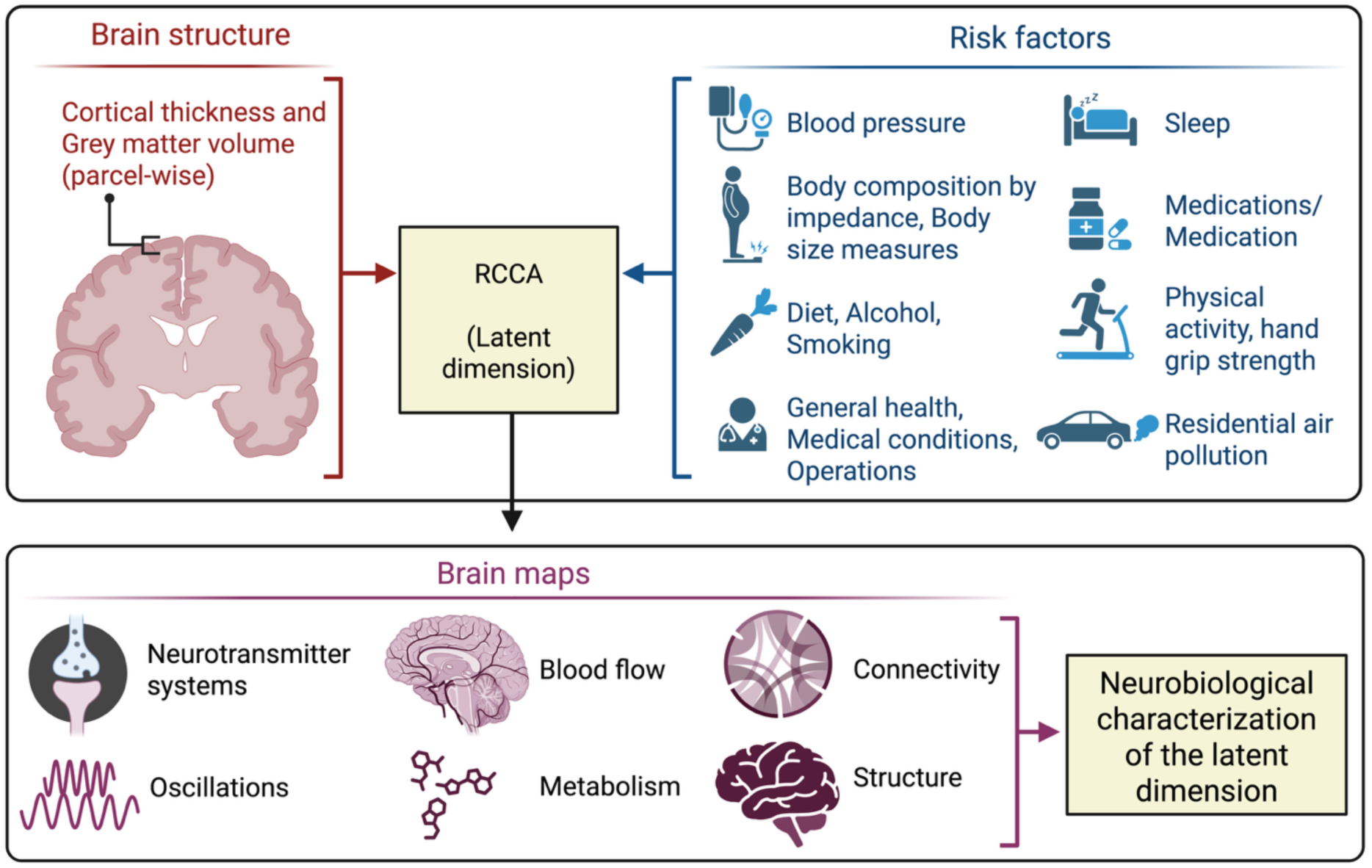
Overview of the analyses. Top panel: Latent dimensions were search using RCCA, linking cortical thickness and grey matter volume measures (parcel-wise across the whole brain cortex) with a wide set of risk factors for non-communicable diseases. Below panel: In order to interpret the latent dimension, the brain loadings were compared with several brain features. Created in BioRender. Nicolaisen, E. (2025) https://BioRender.com/22pyru2

## Methods

### Participants

We used data from UKB (application 41655) [42,43]. Inclusion criteria for this study were having no self-reported illnesses (Data-Field ID 20002-2.*) and having complete data in all variables utilized in this study (excluding responses like “Do not know” or “Prefer not to answer”). To ensure that our sample represent male and female population equally, we selected a sample (n=7370) with equal number of age-matched males and females. We hereafter refer to it as the main or mixed sample, since it includes individuals of both sexes/genders: 3685 women (age: range 46-81 years, mean 62.3, standard deviation 7.6) and 3685 men (age: range 46-81 years, mean 62.3, standard deviation 7.6). In order to check for a potential sex-bias in the results, we also analyzed each sex-specific subsample, separately (see Supplementary Methods 1.1).

### Risk factors data

We identified 68 risk factor variables in the UKB dataset (after removing variables which were duplicated (i.e. same variable measured several times), variables with missing values, and variables with skewed distribution). These 68 variables spanned categories of general health, body size measures, diet, physical activity, residential air pollution, sleep health, alcohol consumption, and smoking (Table S1). As done previously, we computed waist-to-hip ratio dividing waist circumference by hip circumference [13]. All variables corresponding to risk factors were acquired on the Imaging visit, with the exception of those in the Category of “Residential air pollution” (Table S1), which were acquired before the Imaging visit.

In order to first gain a better understanding of the interrelation between risk factors alone (without taking into account brain data), the intercorrelation among risk factors was analyzed with Pearson’s correlation and visualized in correlation matrices.

### Neuroimaging data

Neuroimaging data for the sample used in this study were collected by UKB in four sites using identical protocols and 3T Siemens Skyra scanners with standard Siemens 32-channel receive head coils [44]. T1-weighted structural imaging (3D-MPRAGE, sagittal) was acquired with the following parameters [42,44]: voxel resolution: 1×1×1 mm, FoV: 208×256×256 matrix, TI/TR = 880/2000 ms, in-plane acceleration iPAT=2. T1 images were processed by UKB using a custom pipeline based on FSL [44], including gradient distortion correction, cutting down the field of view, registration (linear and then non-linear) to the MNI152 standard-space T1 template, brain extraction, defacing, and brain segmentation [44]. CT and GMV were estimated using FreeSurfer a2009s.

We used mean raw CT and mean raw GMV of 148 cortical parcels (Destrieux atlas, a2009s) [45]. To check the effect of brain size correction on the results, we also used proportional estimations of CT and GMV (computed subject-wise, dividing the mean raw CT or GMV of each parcel by the mean raw CT or Total Intracranial Volume (TIV), respectively, across all parcels) and CT and GMV corrected for brain size (regressing out mean CT or TIV, respectively, in a cross-validation consistent manner to avoid data leakage) [46]. These three different brain structural measures (raw, proportional, or corrected) represent different biological properties and can show different patterns of associations with risk factors. For instance, raw CT represents the absolute CT of a brain region, while proportional CT can be understood as the ratio between a region’s CT and the whole brain’s CT. Along the same line, corrected CT can be seen as the regional value adjusted for the whole brain value, that is, the regional value which is not explained by the total brain value. Accordingly, proportional and corrected values are relative values, i.e. when taking into account the whole brain value. This means that a corrected CT value for a given region expresses to which extend it has high or low CT when taking into account the global brain structure. Accordingly, corrected values are more likely to reveal structural patterns that pertain to specific brain regions. However, the drawback of correction or adjustment for head/brain size is that it may discard relevant variance, i.e. variance that pertains to the effect we are interested in. For example, if variability in brain/head size is associated with variability in physical robustness for relevant biological reasons (e.g. physically more robust individuals tend to be taller and more massive), adjusting for brain/head size can remove relevant variance related to physical robustness. Therefore, row and uncorrected/unadjusted values have pros and cons and offer different insights in the current study.

### Regularized canonical correlation analysis

Canonical Correlation Analysis (CCA) is a multivariate technique that can discover latent dimensions alinking interindividual variability in ***X*** and ***Y*** [22,23,47]. Here, ***X*** included either CT or GMV for each parcel, while ***Y*** included risk factors. CCA searches linear combinations of variables in ***X*** (brain weights ***u***) and of variables in ***Y*** (risk factor weights ***Y***), which maximize the canonical correlation between the brain scores and risk factor scores [22,23]. The scores correspond to the projection of ***X*** and ***Y*** onto their respective weights (***Xu*** and ***YY***) (these correspond to subject-wise values). One limitation of CCA is that it can overfit the data or yield unstable results, especially in high-dimensional datasets [48]. Therefore, we here used a regularized version of CCA (RCCA) which reduces the overfitting of the model by adding L2-norm constraints to the weights [22,23,49,50].

To search for multivariate associations between risk factors and both, CT and GMV in the main sample, we ran two RCCA models (one per brain structural measure). To check for the effect of brain size correction on the results, we ran four more analyses for cortical models: using either proportional or brain-size-corrected CT and GMV. In addition, to investigate potential sex-bias in the results, we ran 6 additional RCCA models: three in the subsample of women and three in the age-matched subsample of men (see Supplementary Materials 1.1). Age and site effects were regressed out avoiding data leakage in the machine learning framework (regression parameters were estimated in the training set and applied to the training, test, and holdout sets) [46,51]. In the main sample, sex was also regressed out. Confounding variables were included in the ***Y*** matrix to check if their variance was properly removed.

To visualize and interpret the latent dimensions, loadings were computed [22] (these correspond to variable-wise values). Brain loadings correspond to the correlation of the original brain variables (***X***) with brain scores (***Xu***). Similarly, risk factor loadings correspond to the correlation between the risk factors original variables (***Y***) with the respective scores (***YY***). Loadings indicate which variables are more strongly associated with the latent dimension. To interpret the latent dimensions, only stable loadings were considered (loadings whose error bar did not cross zero).

### Machine learning framework

We utilized a machine learning framework that uses multiple holdouts of the data [23,52]. This framework implements two consecutive splits: the outer split divides the whole data into optimization and hold-out sets, and is used for statistical evaluation, and the inner split divides the optimization set into training and test sets and is used for model selection. In this study, we used 5 inner splits and 5 outer splits. Model selection was performed based on the highest test canonical correlation and the highest stability (similarity of weights estimated using Pearson’s correlation across the 5 inner splits).

### Statistical evaluation of the latent dimensions

Statistical significance of the latent dimensions was tested with permutation tests. In each of the 1000 iterations, the rows of the ***Y*** matrix were shuffled in the optimization and hold-out sets. The RCCA model (hyperparameters) that was previously selected with the original data was now fitted on the permuted optimization set, and weights were obtained. Then, the permuted hold-out set was projected onto these weights. A canonical correlation under the null model was hence obtained, and a p-value was computed. The permutation test was repeated for each one of the outer splits, hence yielding 5 p-values which were corrected by multiple comparisons (Bonferroni method over 5 comparisons). The statistical significance of the latent dimensions was evaluated using the omnibus hypothesis [52]. Here, the null hypothesis states that there is no effect in any of the splits. Hence, if at least one split yields a p-value below 0.05, the null hypothesis is rejected, and the latent dimension is considered significant. When a significant latent dimension was found, its variance was removed from the data using deflation [23], and an additional latent dimension was sought.

### Stability of the latent dimensions across sexes and brain structural measures

The latent dimensions were compared based on their average risk factor loadings and average brain loadings (average across the outer 5 splits). The risk factor loadings were compared with Pearson’s correlation across models. The brain loadings were compared using spin test with 10000 permutations to account for spatial dependencies of brain data [53] using neuromaps software [35]. The p-values were corrected by multiple comparisons using Bonferroni method.

### Neurobiological characterization of the brain structural patterns

The brain maps provided in neuromaps [35] (Table S3) (excluding the map ‘hill2010’ which is provided only in one hemisphere) were compared to the maps of CT or GMV loadings using spin test [53] with 10000 permutations. This was performed to assess if the latent dimensions captured a CT or GMV pattern significantly associated with other patterns of brain features. Multiple comparisons were corrected using the Bonferroni method.

### Association of the latent dimensions with demographics

The association of the risk factor scores (subject-wise values) and demographics was analyzed with Spearman correlation (Supplementary materials 1.2, Table S2).

### Association of the latent dimensions with peripheral inflammation

We explore the association between the composite variable of risk factors (i.e. the risk factor scores which are subject-wise values) and a peripheral marker of inflammation with Spearman correlation. We focused on C-reactive Protein (CRP, field-id 30710-1.0 in the UKB dataset) as the most relevant marker for chronic and systemic low-grade inflammation having a good signal-to-noise ratio for population studies, a relative stability in time, and a relatively low missingness in UK Biobank. Multiple comparisons were corrected with Bonferroni.

### Latent dimensions linking risk factors to subcortical and cerebellar volumes

Although the current study focuses on cortical structure, we also added supplementary analyses examining latent dimensions yielded with subcortical and cerebellar structures for readers’ information. For that, we ran one additional RCCA analysis linking the same set of risk factors as before with a set of subcortical and cerebellar volumes in the main/mixed sample (see Supplementary methods 1.3). Age, sex and site were regressed out.

### Ethics

Analyses on the data have been approved by the University Hospital Düsseldorf ethics committee votes 2018-317-RetroDEuA, 2018-317_1-RetroDEuA, and 2018-317_2.

## Results

### Risk factors collinearity

The distribution of risk factors is shown in Figures S1-S2. In the correlation matrices for risk factors, two groups of highly intercorrelated variables were evident (Figures S3-S4). One group shows intercorrelation among body composition measures, including BMI, body fat percentage, body fat mass, body fat-free mass, body water mass, basal metabolic rate (which is estimated from fat-free body mass), impedance of whole body, waist circumference, hip circumference, and waist-to-hip ratio. Another group of highly intercorrelated variables was characterized by air pollution.

### Latent dimension linking cardiometabolic health to brain structure

A first significant latent dimension was found when analyzing the link between risk factors to raw CT in the main/mixed sample (merging women and men) (r_range_ = 0.30-0.34, p_range_ = 0.005-0.005) (Figure 2, Figures S5-S6, Table S4). Conceptually, this latent dimension captures variability in cardiometabolic health, for instance with positive loadings for higher physical activity and a negative pole for factors of metabolic conditions, such as body fat, BMI and waist-to-hip ratio, as well as blood pressure. Of note, when analyzing the link between the same set of risk factors and GMV in the main/mixed sample a latent dimension with a highly similar profile of risk factors loadings was yielded (Figure 2, Figures S5 and S7, Table S7; Figure S8 for comparison across latent dimensions). Note that this latent dimension corresponded to the first yielded latent dimension for all models except for the models for raw GMV, in which it was yielded as second latent dimension.

**Figure 2.**
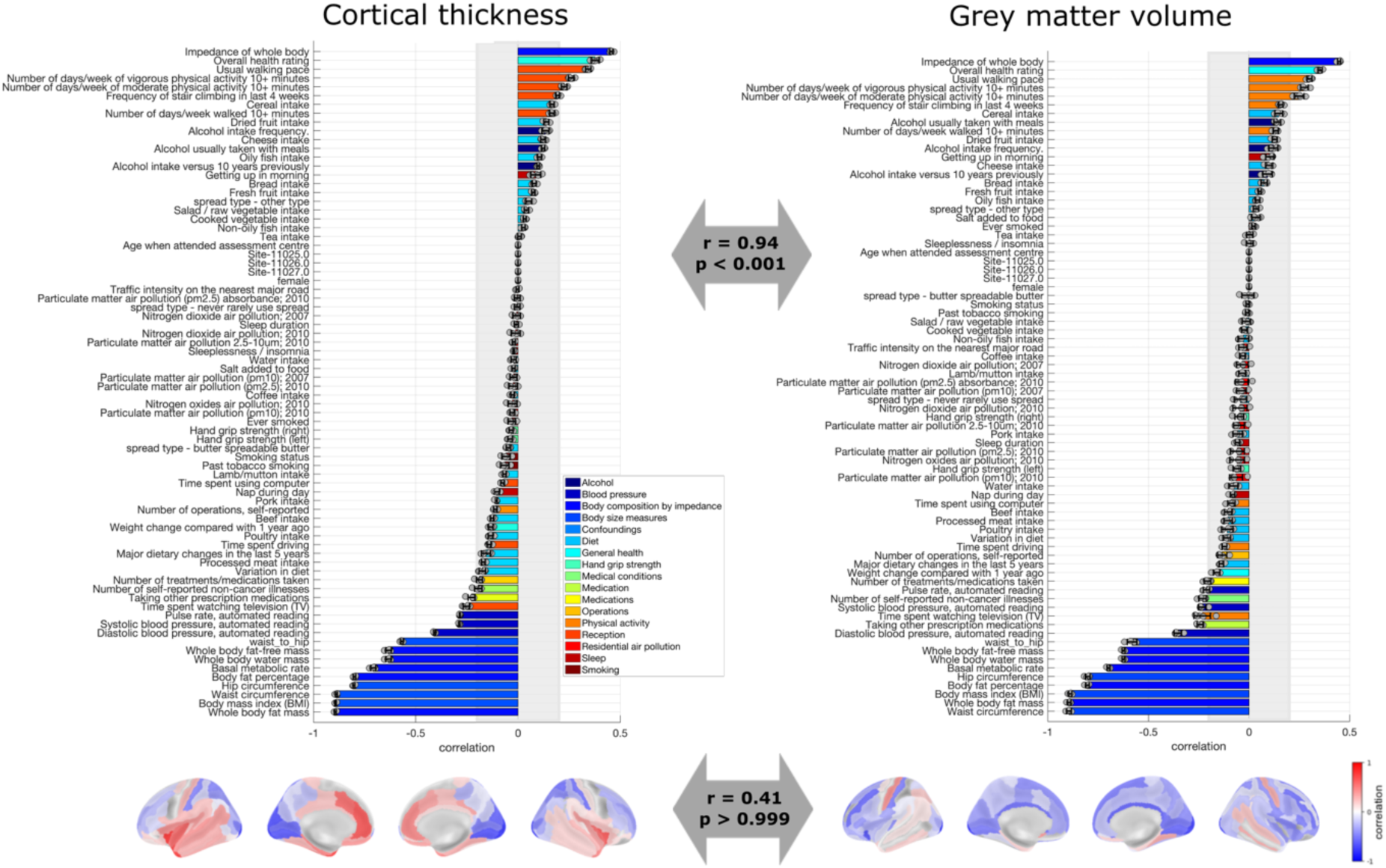
Loadings of the latent dimension of cardiometabolic health. Risk factor loadings and brain loadings in the main/mixed sample for cortical thickness (left) and grey matter volume (right). The arrows depict the Spearman correlation of the loadings across models. The loadings represent the average over the five outer splits. Error bars depict one standard deviation. The shadowed zone marks loadings between −0.2 and 0.2.

The pattern of cardiometabolic health was linked to a specific brain pattern of CT and a specific brain pattern of GMV (Figure 2). The CT pattern shows that better cardiometabolic health is associated with higher CT in the insula, cingulate cortex, temporal lobe, inferior parietal, orbitofrontal, and primary motor cortex, and to lower CT in the primary somatosensory cortex, superior frontal areas, superior parietal areas, and occipital areas. In addition, better cardiometabolic health is associated to higher GMV in primary motor cortex and temporal cortex, and lower GMV in frontal, parietal, and occipital areas.

Notably, this latent dimension of cardiometabolic health was stable across sexes and across brain size corrections (Figure S8, Supplementary results 2.1-2.3). Associations between the Thus, overall, this latent dimension appears to capture a composite aspect of cardiometabolic health that robustly relates to brain structure.

### Latent dimension linking physical robustness to brain structure

Our results showed another significant latent dimension linking risk factors to raw CT (r_range_ = 0.09-0.12, p_range_ = 0.005-0.005) (Figure 3, Figures S25-S26, Table S4). Importantly, when studying raw CT, this latent dimension was significant and stable in the sample of men but not in the sample of women (see Supplementary results 2.5). For this reason, in the next steps we focus on the sample of men when considering this latent dimension with respect to CT. Also, note that this latent dimension corresponded to the second yielded latent dimension for all models except for the models for raw GMV, in which it was yielded as first latent dimension. This latent dimension captured variability in physical robustness (as opposed to physical frailty), with a positive pole associated to for instance whole body fat-free mass, grip strength and overall health rating, and a negative pole associated with smoking and air pollution.

**Figure 3.**
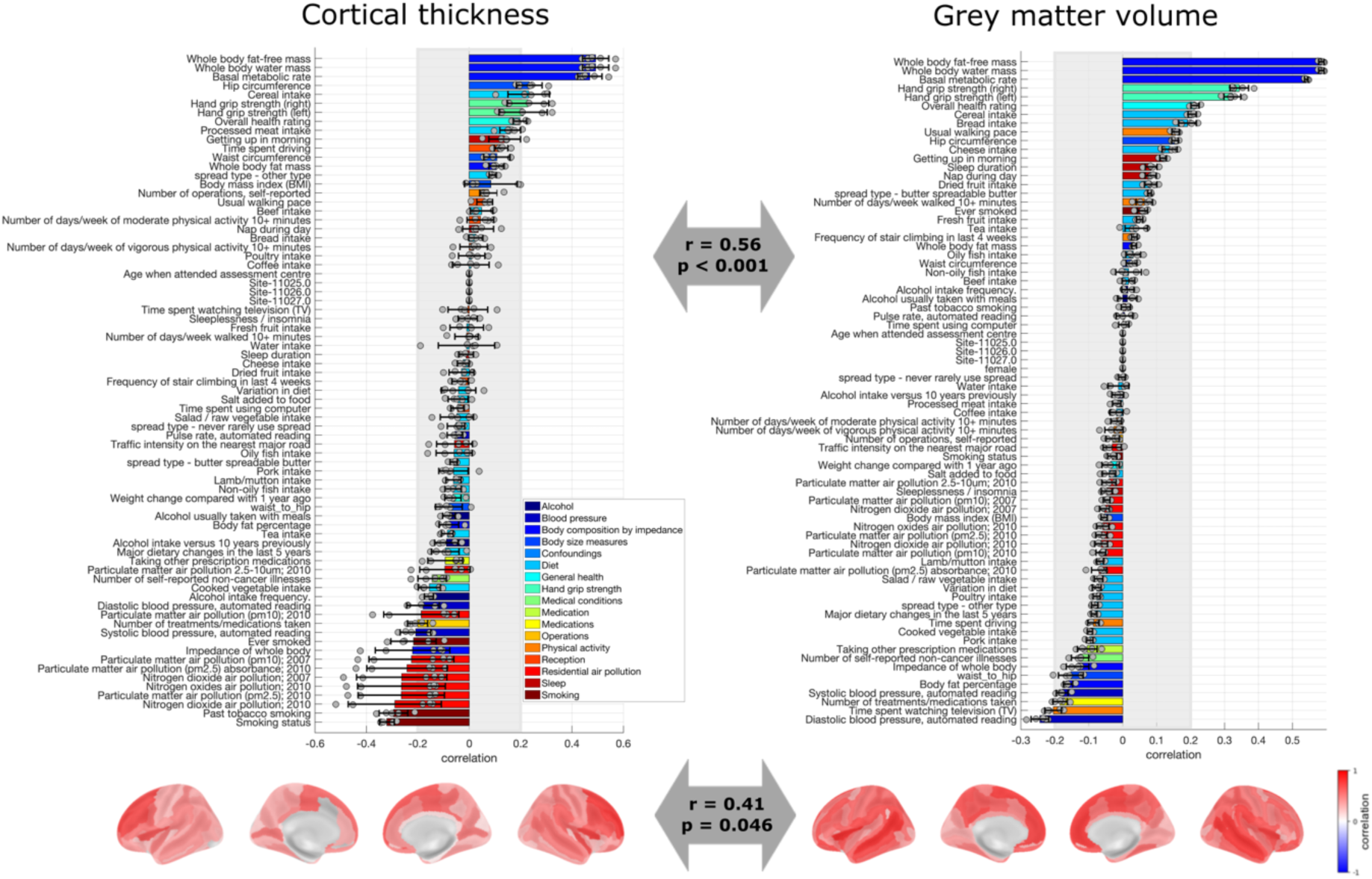
Loadings of the latent dimension of physical robustness. Risk factor loadings and brain loadings in the men sample for cortical thickness (left) and the main/mixed sample for and grey matter volume (right). The arrows depict the Spearman correlation of the loadings across models. The loadings represent the average over the five outer splits. Error bars depict one standard deviation. The shadowed zone marks loadings between −0.2 and 0.2.

This latent dimension of physical robustness was associated with a specific brain pattern of CT and a specific brain pattern of GMV (Figure 3), though the statistical comparison showed a marginally significant Spearman correlation indicating that they do share variance (r = 0.41, p = 0.046). However, this significant comparison was yielded only for the case of raw CT and raw GMV, but not for corrected brain structural measures (Figure S31-f). Conceptually, our results indicated that higher physical robustness was associated with higher CT, especially in superior frontal areas and insula. Moreover, the physical robustness pattern was associated to higher GMV, especially in the anterior cingulate cortex, medial superior frontal areas, orbitofrontal cortex, and temporal lobe.

This latent dimension was stable across sexes when yielded with GMV, but it was not significant/stable in the sample of women when yielded with CT (Supplementary results 2.5-2.7). When yielded with GMV, and when yielded in the sample of men with CT, this latent dimension was stable across brain size correction procedures (Figure S31, Supplementary results 2.5-2.7). Associations between the latent dimensions and demographics are shown Supplementary Results 2.8.

### Neurobiological characterization of the pattern of brain structural loadings associated with cardiometabolic health

To characterize the latent dimensions from a neurobiological perspective, we compared the CT and GMV loadings with brain maps spanning brain function, structure, and neurotransmitter systems.

For the latent dimension associated to cardiometabolic health, several associations between the CT and GMV patterns and different brain maps were significant (Table S10, Figure 4, Figure S41). Namely, the map of CT loadings was positively associated with the cortical distribution of serotonin receptor 5-HT1a [54,55], dopamine transporter DAT [56], receptor GABAa [57] and acetylcholine transporter VAChT [34,58]. In addition, the CT pattern was positively associated with cortical thickness [59], and fifth gradient of resting-state functional connectivity (RSFC) [60], as well as negatively associated with glucose metabolism [61], histone deacetylase [62] and with the first principal component of genes in the Allen Human Brain Atlas [63,64]. In turn, the GMV pattern of the cardiometabolic health latent dimension was positively associated with the cortical distribution of dopamine receptor D2 [65] and dopamine transporter DAT [66], and negatively associated with serotonin receptor 5-HT1b [34,67], and histamine receptor H3 [68]. Moreover, the GMV pattern was positively associated with the 8^th^ gradient of RSFC [60], and negatively associated with histone deacetylase [62] and 10th gradient of RSFC [60].

**Figure 4.**
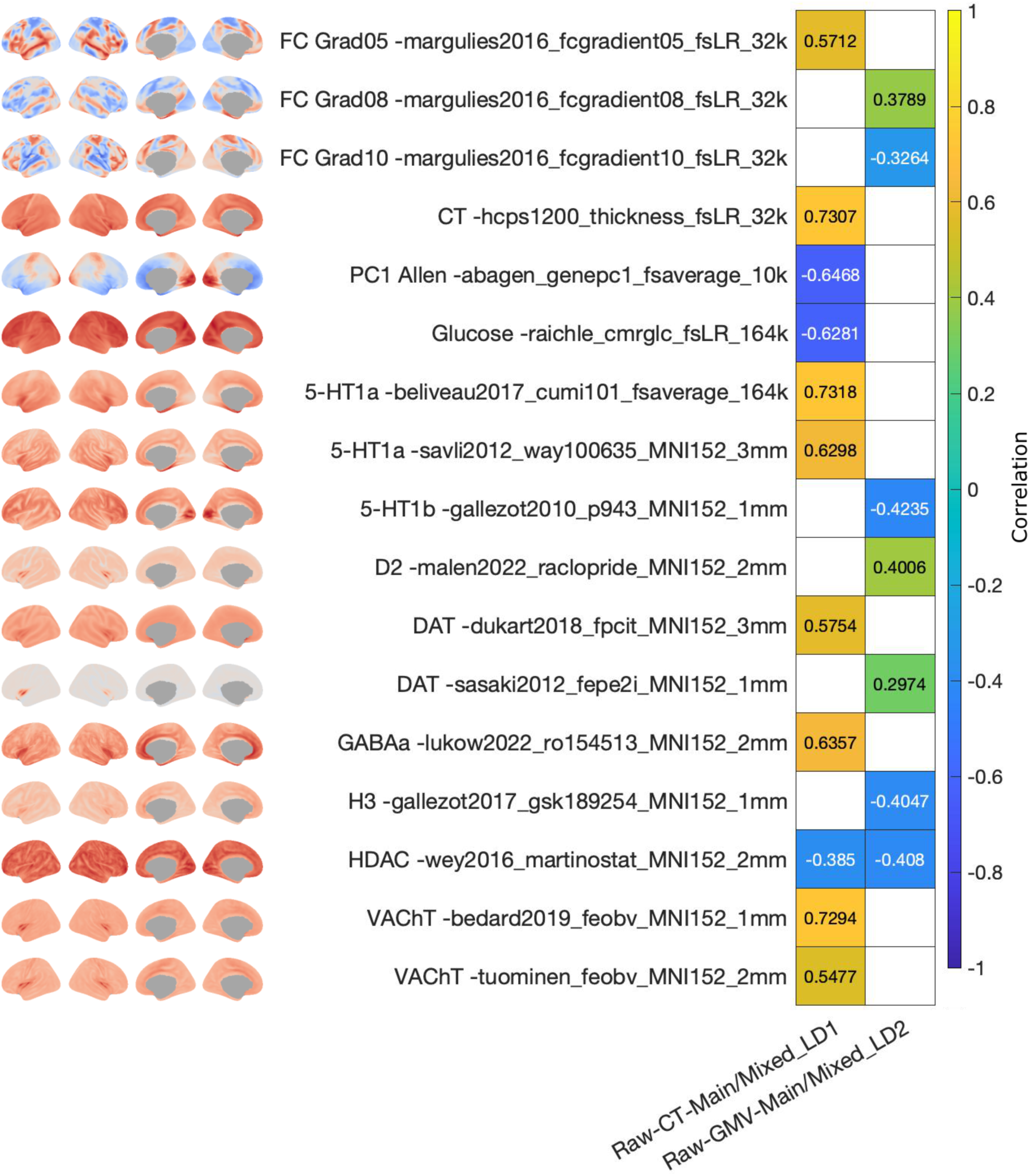
Association of brain structural loadings with neuromaps for the latent dimension of cardiometabolic health. Only data for neuromaps that yielded a significant association with at least one of the shown loadings maps are depicted. Colorbar represents the Spearman correlation. White tiles represent non-significant associations.

Conceptually, this indicates that the latent dimension captured a brain axis in which cortical regions whose CT covaried positively the most (increased the most, red regions in Figure 2) with the pattern of cardiometabolic health were regions that showed the highest binding potentials for 5-HT1a, DAT, GABAa and VAChT. Similarly, the brain regions whose GMV increased the most (red regions) with cardiometabolic health were regions with the highest binding potential for D2 and DAT. In turn, the brain regions whose GMV covaried negatively the most (decreased the most, blue regions) with cardiometabolic health were regions with the highest binding potentials for 5-HT1b and H3.

### Neurobiological characterization of the pattern of brain structural loadings associated with physical robustness

The brain patterns yielded for the latent dimension of physical robustness also showed several significant associations with brain maps (Table S11, Figure 5, Figure S42). The CT pattern yielded in the sample of men was positively associated with glutamate receptor mGluR5 [34,69], RSFC intersubject variability [70], and developmental area scaling [71]. The GMV pattern was positively associated with the serotonin receptors 5-HT1a [54], with glutamate receptors mGluR5 [34,69,72] and NMDA [34,73–75], with histamine receptor H3 [68], with opioid receptors MOR [76,77] and KOR [78], with cannabinoid receptor CB1 [79,80], and with receptor GABAa [57]. In addition, the GMV pattern was positively associated with the 1^st^ gradient of RSFC [60], intersubject variability of RSFC [70], cortical thickness [59], developmental area scaling [71], evolutionary expansion [81] and sensory-association axis [82]. Moreover, the GMV pattern was negatively associated with T1w/T2w [59] and with functional homology [81].

**Figure 5.**
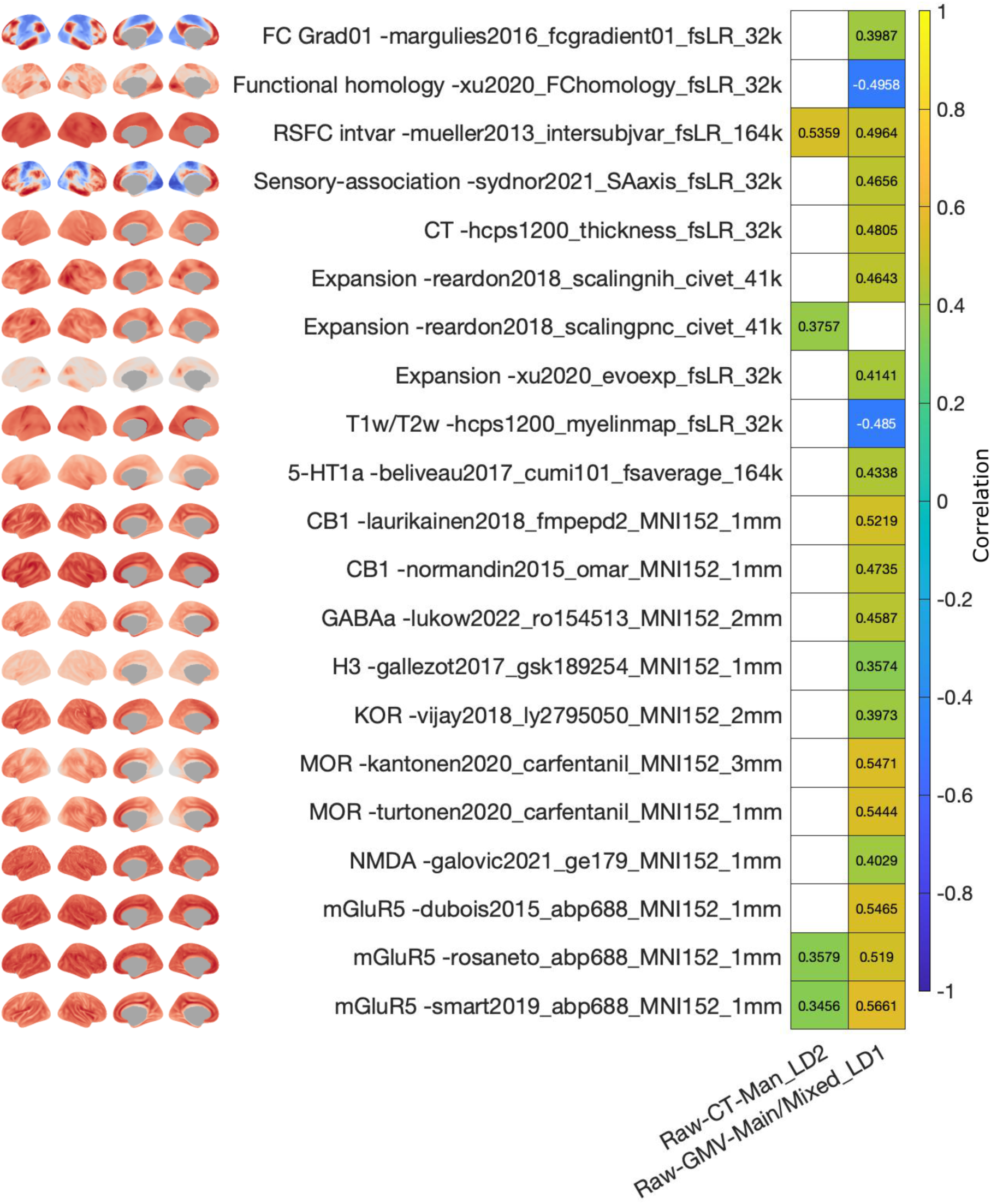
Association of brain structural loadings with neuromaps for the latent dimension of physical robustness. Only data for neuromaps that yielded a significant association with at least one of the shown loadings maps are depicted. Colorbar represents the Spearman correlation. White tiles represent non-significant associations.

Conceptually, this indicates that the latent dimension captured a brain axis in which cortical regions whose CT increased the most with the pattern of physical robustness (red regions in Figure 3) were those with highest binding potential for mGluR5. In addition, regions whose GMV increased the most with physical robustness (red regions) were those with highest binding potential for 5-HT1a, mGluR5, NMDA, H3, MOR, KOR, CB1 and GABAa.

### Correlation of the latent dimensions with peripheral marker of inflammation

CRP was significantly associated to the risk factors scores of cardiometabolic health when yielded in association with raw CT (r = -0.39, p < 0.001) and with raw GMV (r = -0.39, p < 0.001) in the main/mixed sample. It was also significantly associated to the risk factors pattern of physical robustness yielded in association with raw GMV in the main/mixed sample (r = - 0.08, p = 0.04) but with a very low effect size. CRP was not significantly associated with the risk factors yielded in association with raw CT in the sample of men (r = 0.038, p = 0.999). Thus, overall, the composite variable of cardiometabolic health appears moderately correlated to a marker of low-grade systemic inflammation, while physical robustness was only very weakly associated (r < 0.1) to such a marker.

### Latent dimensions linking risk factors to subcortical and cerebellar volumes

Results yielded by the analysis of subcortical and cerebellar data are reported in Supplementary results section 2.9. Overall, more significant latent dimensions appeared and only one was partly related to the two main dimensions or risk factors found in association to cortical structure. This suggests that subcortical regions have different patterns of interindividual variability than cortical regions (likely higher interindividual variability and more vulnerability to exposome factors). Accordingly, their associations with risk factors should be the focus of specific studies. In the following discussion, we focus on cortical structure.

## Discussion

We report two latent dimensions characterizing the interplay between a wide range of risk factors for non-communicable diseases with region-wise CT and GMV across the whole cortex. One of these latent dimensions highlighted the relevance of cardiometabolic health for inter-individual variability of brain structure in a healthy sample. Importantly, this latent dimension was stable across sexes, across brain structural measures (CT and GMV) and across brain size corrections (raw, proportional, and brain-size corrected CT). Accordingly, this latent dimension cannot be explained by a confounding effect of head size/morphology that could be conveyed in brain structural estimates. The other latent dimension highlights the relationship between physical robustness and brain structure. Even though this latent dimension was stable within analyses, the stability across sexes, brain structural measures, and brain size corrections was limited. In addition, our results underline the multi-level nature of the association between risk factors and brain structure, linking the brain patterns of both latent dimensions with the spatial distribution of several neurotransmitter systems.

We showed that the variability in cardiometabolic health is related to an axis of variability in CT extending from the insula and cingulate cortex to the occipital lobes and superior parietal regions. Overall, this latent dimension indicates that regions engaged in processing internal information such as emotional and motivational systems (including for instance insula, orbitofrontal cortex, and anterior cingulate cortex) show positive loadings. Accordingly, this CT pattern was significantly related to dopamine and serotonin systems. In contrast, more dorsal regions typically engaged in dorsal attention and executive systems (such as the lateral superior prefrontal and parietal cortex) show negative CT loadings on the latent dimension. Similarly, cardiometabolic health was negatively associated with GMV in the frontal regions. However, it was positively associated with GMV primarily in sensorimotor regions (peri-central regions), but also to a lesser extent to the ventral attention network (lateral parietal and middle temporal regions). This suggests that cardiometabolic health may be differently related to variability in CT and variability in GMV in the population. More concretely, our results suggest that better cardiometabolic health is related to increased CT in emotional and motivational systems along with increased GMV in sensorimotor cortex. Future studies should further elucidate the mechanism behind these associations to identify whether improving cardiometabolic health can promote brain structural health in motivation, emotion and sensorimotor networks and hence improve mental health and motor function in aging. The potential role of neurotransmitters system in these mechanisms is further discussed below.

In the current stage of knowledge, our results are in line with recent reports pointing to the important role of cardiometabolic health for brain health. For instance, several studies have linked cardiometabolic factors to brain structure, such as reduced total brain volumes [38,83,84], reduced grey matter volumes [31,38,84], or to structural markers of brain aging [85]. Interestingly, our findings are in contrast with works that have reported only reductions, but not increases, in brain structural measures in association with risk factors or cardiometabolic health. Differences in the health status of the samples might explain these differential effects. Apart from associations with brain structure, cardiometabolic health has also been associated with brain function, such as cognition [38], dementia [83], and other neuropsychiatric disorders and symptoms [21,29,38,86,87]. In our study, a moderate correlation between the composite variable of cardiometabolic health and a marker of peripheral inflammation further suggests a role for low-grade chronic inflammation in the association between cardiometabolic health and brain health.

Actually, several studies have pointed out that a causal factor or mediator in the association between risk factors for non-communicable diseases and brain health might be inflammation [36–40], and in particular low-grade systemic inflammation [21,36,38,39,88,89]. Risk factors for non-communicable diseases are also risk factors for low-grade systemic chronic inflammation [6,39,89]. In turn, low-grade systemic chronic inflammation has been associated with non-communicable diseases [6,88], including neuropsychiatric illnesses [21,38,39,88–92]. For instance, chronic inflammation in the adipose tissue has been associated with the development of neurodegenerative disorders such as Alzheimer’s disease [39,93]. Moreover, inflammatory factors have been reported to mediate the association between BMI and other risk factors with cortical structure and behavior [36,37] (however, see [94]). In fact, inflammation has been proposed as the cause of comorbidities between non-communicable diseases [86,90], such as depression and cardiovascular or neurodegenerative illnesses [90]. Thus, we can speculate here that cardiometabolic health, partly via inflammation pathways, influence brain health with potential impacts on emotion and motivation behavioral systems, as well as to motor systems.

Our results also show that interindividual variability in physical robustness (fat-free mass and strength) is associated with interindividual variability in brain structure, in both men and women for GMV, but only robustly in men for CT. At the brain level, this CT pattern in men shows highest loadings mainly across the whole lateral frontal cortex and the superior medial frontal gyrus. The pattern of GMV associated to physical robustness also shows high loadings on medial frontal regions, along with lateral temporal regions. Accordingly, this GMV pattern is also significantly related to cortical patterns of developmental area scaling, along with evolutionary expansion and sensory-association axis. This may suggest common mechanisms underlying physical health (likely musculoskeletal health) and brain health in key regions for human high-level cognition (such as the frontal cortex). Interestingly, time spent watching television appears on the negative pole of these dimensions suggesting that sedentary lifestyles may threaten this aspect of human brain health.

It should be noted that the variability of physical robustness in women appears to be linked to GMV but not to CT, while in men it is associated to both structural markers. This sex-variability could be explained by the risk factors profile in association with CT in men capturing a negative pole of air pollution and smoking. Previous studies have pointed towards sex-variability in the effect of air pollution on health and particularly on brain and behavior [95–99]. This could be due to sex-specific exposures to pollutants, with more interindividual variability in exposures in men allowing us to observe associations with health outcomes. In line with this hypothesis, the distribution of variables related to air pollution in our sample of men include a few more extreme values than in women (Figure S2). Alternatively, a potential sex-specific effect of air pollutants could also be due to different biological aspects being differentially affected in men in comparison to women, or to sex-specific pathways underpinning the association between air pollution and brain structure. The mechanisms underpinning this difference should be explored in future studies. Moreover, studies analyzing the association between health variables and neuroimaging should consider this and perform sex-specific analyses. In the current stage of knowledge, we can only speculate from our results that air pollution and smoking may represent a specific threat for men brain health in high-level associations regions.

Interestingly, the brain patterns associated to both dimensions of risk factors were generally significantly correlated to the spatial distribution of most neurotransmitter systems. It should also be noted that the neurotransmitters associated to the latent dimension have in turn been linked to phenotypes related to the risk factors captured in the latent dimension. For instance, the serotoninergic system is associated with energy balance and feeding behavior [100,101], obesity [100] and physical activity [102]. Specifically, the 5-HT1a receptor has been associated with food intake [101], anorexia nervosa and bulimia nervosa [103]. The dopaminergic system has been associated with physical activity [102], and specifically, the receptor D2 has been associated with disorders related to eating behavior, such as anorexia nervosa, bulimia nervosa, and obesity [103]. Overall, the evidence indicates that these neurotransmitter systems are associated with imbalances in energy homeostasis and feeding behavior, which are phenotypes related to the pattern of risk factors captured in our latent dimension. This suggests that the clinically relevant interaction between brain and body health that has been called to attention recently [21,29] might be mediated by processes associated with neurotransmitter systems and/or importantly alter neurotransmitter systems. For instance, cardiometabolic factors may lead to perturbations in the serotoninergic system via brain structural alterations ultimately leading to immune-metabolic depression [104].

Evidence of the mechanistic cause linking neurotransmitter systems with both, risk factors for non-communicable diseases and brain structure, also points to inflammation. Several studies have shown a crosstalk between the immune system and several neurotransmitter systems, such as serotoninergic, noradrenergic and dopaminergic systems [92,105]. For instance, certain neurotransmitter receptors, including 5-HT1a, D1 and D2, have immunologic functions [88,105], and can lead to the disruption of homeostasis [88] and to inflammation. Accordingly, immune cells express neurotransmitter receptors [105]. In turn, immunological factors can regulate normal cellular functions, including neurotransmission and synaptic plasticity [88]. In sum, the crosstalk between inflammatory factors and neurotransmitter systems is bidirectional [88,105] and is relevant for several non-communicable diseases [105]. Moreover, since these neurotransmitter systems are implicated in mental disorders and in somatic non-communicable diseases [88,90], the mechanisms underlying the comorbidity between mental and somatic disorders might be partly related with alterations in neurotransmitter systems [86]. Accordingly, it is likely that a vicious cycle is engaged when cardiometabolic and musculoskeletal systems, immunoinflammatory systems, brain structure, and neurotransmitters systems are altered complexifying treatment of neuropsychiatric disorders.

In that context, our findings contribute to the recognized urgency to characterize brain-body interactions for its implementation in clinical practice [21,29,86]. For instance, it is not common to monitor physical illnesses in patients with neuropsychiatric disorders. However, recently it has been pointed out that neuropsychiatric disorders are associated with symptoms of physical illnesses and that poor physical health is a more pronounced effect than brain phenotypes in these patients [29]. Given that the interplay between brain and body health is not well understood, the clinical practice nowadays has limited tools to exploit this interaction not only for a comprehensive monitoring of health, but also for its use as biomarkers. Hence, research characterizing brain and body interactions is a major priority for global health because it will guide the discovery of new integrated disease manifestations and pave the way for the development of new therapies and clinical interventions [21,29].

## Conclusion

Our study shows two latent dimensions linking cardiometabolic health and physical robustness to patterns of CT and GMV variability across the whole cortex. In turn, the captured CT and GMV brain patterns were associated with the cortical distribution of several neurotransmitter systems. Furthermore, cardiometabolic health was associated to a peripheral marker of low-grade systemic inflammation. Hence, our study shows that body health, brain structure, and neurotransmitter systems are interrelated, highlighting the multi-level nature of health. Also, our work contributes to questioning the classic consideration of neuropsychiatric and somatic illnesses as separate categories [88] and supports the view of a needed integration of brain and physical health in clinical practice [21,29].

## Declarations

### Authors’ contributions

ENS conceptualized the study, developed software, prepared data, performed analyses, and contributed to discussion and interpretation of results. SMB contributed to study conceptualization, data preparation, software development, and to interpretation and discussion of results. MM contributed to data preparation and software development. AM contributed to software and methodology development. FH contributed to data preparation and preprocessing. JMM contributed to software and methodology development. MT contributed to discussion and interpretation of results. SBE acquired funding and contributed to discussion and interpretation of results. SG acquired funding, conceptualized the study, and contributed to discussion and interpretation of results. The contribution of ENS has been done in partial fulfillment of the requirements for a Ph.D. thesis. All authors contributed to the revision of the manuscript. All authors contributed to writing and reviewing of the manuscript and approved the final version. ENS and SMB have accessed and verified the data.

## Acknowledgements

This research has been conducted using the UK Biobank Resource under Application Number 41655. This work was supported by the Deutsche Forschungsgemeinschaft (DFG, GE 2835/2– 1, GE 2835/9-1, SFB 1451 – Project-ID 431549029) (https://www.dfg.de/), granted to SG. SG received salary from GE 2835/2–1. MM received salary from GE 2835/9-1. SMB is supported by the MODS project funded from the programme “Profilbildung 2020” (grant no. PROFILNRW-2020-107-A), an initiative of the Ministry of Culture and Science of the State of Northrhine Westphalia (https://www.mkw.nrw/). SMB is supported by the EBRAINS 2.0 Project funded from the European Union’s Horizon Europe Programme under the Specific Grant Agreement No. 101147319. EBRAINS is funded by the Horizon Europe Framework Programme (©2023 https://ebrains.eu, https://research-and-innovation.ec.europa.eu/funding/funding-opportunities/funding-programmes-and-open-calls/horizon-europe_en). The funders did not play any role in the study design, data collection and analysis, decision to publish, or preparation of the manuscript.

## Data and code sharing statement

Access to UKB data is explained at https://www.ukbiobank.ac.uk/enable-your-research. The code used for the machine learning framework (https://doi.org/10.5281/zenodo.7153571) has been made publicly available at https://github.com/mlnl/cca_pls_toolkit

## Captions

### 1. Supplementary methods

1.1 Sex-specific analyses.

1.2 Comparison of latent dimensions with demographics.

1.3 Latent dimensions linking risk factors to subcortical and cerebellar volumes.

### 2. Supplementary results

2.1 Latent dimensions linking cardiometabolic health to raw cortical thickness and raw grey matter volume.

2.2 Latent dimensions linking cardiometabolic health to proportional cortical thickness and proportional grey matter volume.

2.3 Latent dimensions linking cardiometabolic health to corrected cortical thickness and corrected grey matter volume.

2.4 Latent dimension of cardiometabolic health and demographics.

2.5 Latent dimensions linking physical robustness to raw cortical thickness and raw grey matter volume.

2.6 Latent dimensions linking physical robustness to proportional cortical thickness and proportional grey matter volume.

2.7 Latent dimensions linking physical robustness to corrected cortical thickness and corrected grey matter volume.

2.8 Latent dimension of physical robustness and demographics.

2.9 Latent dimensions linking risk factors to subcortical and cerebellar volumes

### 3. Supplementary tables

Table S1. Variables of risk factors used in this study.

Table S2. Demographic variables used in this study.

Table S3. Brain maps included in neuromaps that were compared with the latent dimension.

Table S4. Statistics of the latent dimensions for raw cortical thickness.

Table S5. Statistics of the latent dimensions for raw grey matter volume.

Table S6. Statistics of the latent dimensions for proportional cortical thickness.

Table S7. Statistics of the latent dimensions for proportional grey matter volume.

Table S8. Statistics of the latent dimensions for corrected cortical thickness.

Table S9. Statistics of the latent dimensions for corrected grey matter volume.

Table S10. Association between the brain loadings of the latent dimension of cardiometabolic health and neuromaps.

Table S11. Association between the brain loadings of the latent dimension of physical robustness and neuromaps.

Table S12. Statistics of the latent dimensions for subcortical and cerebellar volumes.

### 4. Supplementary figures

Figure S1. Distribution of risk factors. Part 1.

Figure S2. Distribution of risk factors. Part 2.

Figure S3. Correlation matrix of risk factors in women.

Figure S4. Correlation matrix of risk factors in men.

Figure S5. Latent dimension of cardiometabolic health.

Figure S6. Model optimization for the latent dimension of cardiometabolic health for raw cortical thickness in the main sample.

Figure S7. Model optimization for the latent dimension of cardiometabolic health for raw grey matter volume in the main sample.

Figure S8. Comparison of risk factor loadings and brain loadings for the cardiometabolic health latent dimension across samples and brain structural measures.

Figure S9. Loadings of the latent dimension of cardiometabolic health for raw cortical thickness in women.

Figure S10. Loadings of the latent dimension of cardiometabolic health for raw cortical thickness in men.

Figure S11. Loadings of the latent dimension of cardiometabolic health for raw grey matter volume in women.

Figure S12. Loadings of the latent dimension of cardiometabolic health for raw grey matter volume in men.

Figure S13. Loadings of the latent dimension of cardiometabolic health for proportional cortical thickness in the main sample.

Figure S14. Loadings of the latent dimension of cardiometabolic health for proportional cortical thickness in women.

Figure S15. Loadings of the latent dimension of cardiometabolic health for proportional cortical thickness in men.

Figure S16. Loadings of the latent dimension of cardiometabolic health for proportional grey matter volume in the main sample.

Figure S17. Loadings of the latent dimension of cardiometabolic health for proportional grey matter volume in women.

Figure S18. Loadings of the latent dimension of cardiometabolic health for proportional grey matter volume in men.

Figure S19. Loadings of the latent dimension of cardiometabolic health for corrected cortical thickness in the main sample.

Figure S20. Loadings of the latent dimension of cardiometabolic health for corrected cortical thickness in women.

Figure S21. Loadings of the latent dimension of cardiometabolic health for corrected cortical thickness in men.

Figure S22. Loadings of the latent dimension of cardiometabolic health for corrected grey matter volume in the main sample.

Figure S23. Loadings of the latent dimension of cardiometabolic health for corrected grey matter volume in women.

Figure S24. Loadings of the latent dimension of cardiometabolic health for corrected grey matter volume in men.

Figure S25. Latent dimension of physical robustness.

Figure S26. Model optimization for the latent dimension of physical robustness for raw cortical thickness in the sample of men.

Figure S27. Model optimization for the latent dimension of physical robustness for raw grey matter volume in the main sample.

Figure S28. Loadings of the latent dimension of physical robustness for raw cortical thickness in women.

Figure S29. Loadings of the latent dimension of physical robustness for raw grey matter volume in women.

Figure S30. Loadings of the latent dimension of physical robustness for raw grey matter volume in men.

Figure S31. Comparison of risk factor loadings and brain loadings for the physical robustness latent dimension across samples and brain structural measures.

Figure S32. Loadings of the latent dimension of physical robustness for proportional cortical thickness in men.

Figure S33. Loadings of the latent dimension of physical robustness for proportional grey matter volume in the main sample.

Figure S34. Loadings of the latent dimension of physical robustness for proportional grey matter volume in women.

Figure S35. Loadings of the latent dimension of physical robustness for proportional grey matter volume in men.

Figure S36. Loadings of the latent dimension of physical robustness for corrected cortical thickness in men.

Figure S37. Loadings of the latent dimension of physical robustness for corrected cortical thickness in women.

Figure S38. Loadings of the latent dimension of physical robustness for corrected grey matter volume in the main sample.

Figure S39. Loadings of the latent dimension of physical robustness for corrected grey matter volume in women.

Figure S40. Loadings of the latent dimension of physical robustness for corrected grey matter volume in men.

Figure S41. Association of brain structural loadings with brain maps for the latent dimension of cardiometabolic health.

Figure S42. Association of brain structural loadings with brain maps for the latent dimension of physical robustness.

Figure S43. Loadings of the first latent dimension linking risk factors to subcortical and cerebellar volumes.

Figure S44. Loadings of the second latent dimension linking risk factors to subcortical and cerebellar volumes.

Figure S45. Loadings of the third latent dimension linking risk factors to subcortical and cerebellar volumes.

Figure S46. Loadings of the fourth latent dimension linking risk factors to subcortical and cerebellar volumes.

Figure S47. Loadings of the fifth latent dimension linking risk factors to subcortical and cerebellar volumes.

Figure S48. Comparison of risk factor loadings between latent dimensions yielded with subcortex-cerebellum and with CT and GMV in cortex.

## Bibliography

[1] World Health Organization. Noncommunicable diseases: progress monitor 2022. 2022.

[2] World Health Organization. Noncommunicable diseases country profiles 2018. 2018.

[3] Lloyd-Jones DM, Allen NB, Anderson CAM, Black T, Brewer LC, Foraker RE, et al. Life’s Essential 8: Updating and Enhancing the American Heart Association’s Construct of Cardiovascular Health: A Presidential Advisory from the American Heart Association. Circulation 2022;146:E18–43. 10.1161/CIR.0000000000001078.

[4] Livingston G, Huntley J, Liu KY, Costafreda SG, Selbæk G, Alladi S, et al. Dementia prevention, intervention, and care: 2024 report of the Lancet standing Commission. The Lancet 2024;404:572–628. 10.1016/S0140-6736(24)01296-0.

[5] Farina FR, Bridgeman K, Gregory S, Crivelli L, Foote IF, Jutila OEI, et al. Next generation brain health: transforming global research and public health to promote prevention of dementia and reduce its risk in young adult populations. The Lancet Healthy Longevity 2024;5. 10.1016/j.lanhl.2024.100665.

[6] Furman D, Campisi J, Verdin E, Carrera-Bastos P, Targ S, Franceschi C, et al. Chronic inflammation in the etiology of disease across the life span. Nature Medicine 2019;25:1822–32. 10.1038/s41591-019-0675-0.

[7] Beydoun MA, Beydoun HA, Gamaldo AA, Teel A, Zonderman AB, Wang Y. Epidemiologic studies of modifiable factors associated with cognition and dementia: systematic review and meta-analysis. BMC Public Health 2014;14:1–33. 10.1186/1471-2458-14-643.

[8] Opel N, Thalamuthu A, Milaneschi Y, Grotegerd D, Flint C, Leenings R, et al. Brain structural abnormalities in obesity: relation to age, genetic risk, and common psychiatric disorders: Evidence through univariate and multivariate mega-analysis including 6420 participants from the ENIGMA MDD working group. Molecular Psychiatry 2021;26:4839–52. 10.1038/s41380-020-0774-9.

[9] Medic N, Ziauddeen H, Ersche KD, Farooqi IS, Bullmore ET, Nathan PJ, et al. Increased body mass index is associated with specific regional alterations in brain structure. International Journal of Obesity 2016;40:1177–82. 10.1038/ijo.2016.42.

[10] Tüngler A, Auwera SV der, Wittfeld K, Frenzel S, Terock J, Röder N, et al. Body mass index but not genetic risk is longitudinally associated with altered structural brain parameters. Scientific Reports 2021;11:1–11. 10.1038/s41598-021-03343-3.

[11] Ronan L, Alexander-bloch AF, Wagstyl K, Farooqi S, Brayne C, Tyler LK, et al. Obesity associated with increased brain age from midlife. Neurobiology of Aging 2016;47:63–70. 10.1016/j.neurobiolaging.2016.07.010.

[12] Kim H, Kim C, Seo SW, Na DL, Kim HJ, Kang M, et al. Association between body mass index and cortical thickness: Among elderly cognitively normal men and women. International Psychogeriatrics 2015;27:121–30. 10.1017/S1041610214001744.

[13] Gurholt TP, Kaufmann T, Frei O, Alnæs D, Haukvik UK, Meer D van der, et al. Population-based body–brain mapping links brain morphology with anthropometrics and body composition. Translational Psychiatry 2021;11. 10.1038/s41398-021-01414-7.

[14] Caunca MR, Gardener H, Simonetto M, Cheung YK, Alperin N, Yoshita M, et al. Measures of obesity are associated with MRI markers of brain aging: The Northern Manhattan Study. Neurology 2019;93:e791–803. 10.1212/WNL.0000000000007966.

[15] Kharabian-Masouleh S, Beyer F, Lampe L, Loeffler M, Luck T, Riedel-Heller SG, et al. Gray matter structural networks are associated with cardiovascular risk factors in healthy older adults. Journal of Cerebral Blood Flow and Metabolism 2018;38:360–72. 10.1177/0271678X17729111.

[16] Chen X, Wen W, Anstey KJ, Sachdev PS. Effects of cerebrovascular risk factors on gray matter volume in adults aged 60-64 years: A voxel-based morphometric study. Psychiatry Research - Neuroimaging 2006;147:105–14. 10.1016/j.pscychresns.2006.01.009.

[17] Vakhrusheva J, Marino B, Stroup TS, Kimhy D. Aerobic Exercise in People with Schizophrenia: Neural and Neurocognitive Benefits. Current Behavioral Neuroscience Reports 2016;3:165–75. 10.1007/s40473-016-0077-2.

[18] Gu Y, Brickman AM, Stern Y, Habeck CG, Razlighi QR, Luchsinger JA, et al. Mediterranean diet and brain structure in a multiethnic elderly cohort. Neurology 2015;85:1744–51. 10.1212/WNL.0000000000002121.

[19] Jacków-Nowicka J, Podgórski P, Bladowska J, Szcześniak D, Rymaszewska J, Zatońska K, et al. The Impact of Common Epidemiological Factors on Gray and White Matter Volumes in Magnetic Resonance Imaging–Is Prevention of Brain Degeneration Possible? Frontiers in Neurology 2021;12:1–11. 10.3389/fneur.2021.633619.

[20] Xu J, Liu N, Polemiti E, Garcia-Mondragon L, Tang J, Liu X, et al. Effects of urban living environments on mental health in adults. Nature Medicine 2023;29:1456–67. 10.1038/s41591-023-02365-w.

[21] Firth J, Siddiqi N, Koyanagi A, Siskind D, Rosenbaum S, Galletly C, et al. The Lancet Psychiatry Commission: a blueprint for protecting physical health in people with mental illness. The Lancet Psychiatry 2019;6:675–712. 10.1016/S2215-0366(19)30132-4.

[22] Mihalik A, Chapman J, Adams RA, Winter NR, Ferreira FS, Shawe-Taylor J, et al. Canonical Correlation Analysis and Partial Least Squares for Identifying Brain– Behavior Associations: A Tutorial and a Comparative Study. Biological Psychiatry: Cognitive Neuroscience and Neuroimaging 2022;7:1055–67. 10.1016/j.bpsc.2022.07.012.

[23] Mihalik A, Ferreira FS, Moutoussis M, Ziegler G, Adams RA, Rosa MJ, et al. Multiple Holdouts With Stability: Improving the Generalizability of Machine Learning Analyses of Brain–Behavior Relationships. Biological Psychiatry 2020;87:368–76. 10.1016/j.biopsych.2019.12.001.

[24] Han F, Gu Y, Brown GL, Zhang X, Liu X. Neuroimaging contrast across the cortical hierarchy is the feature maximally linked to behavior and demographics. NeuroImage 2020;215:116853. 10.1016/j.neuroimage.2020.116853.

[25] Küppers V, Bi H, Nicolaisen-Sobesky E, Hoffstaedter F, Yeo BTT, Drzezga A, et al. Lower motor performance is linked with poor sleep quality, depressive symptoms, and grey matter volume alterations. bioRxiv 2024:2024.06.07.597666. 10.1101/2024.06.07.597666.

[26] Maleki Balajoo S, Plachti A, Nicolaisen-Sobesky E, Dong D, Hoffstaedter F, Meuth S, et al. Data-driven identification of a left anterior hippocampus’ morphological network associated with self-regulation. Research Square 2024. 10.21203/rs.3.rs-4170788/v1.

[27] Mihalik A, Ferreira FS, Rosa MJ, Moutoussis M, Ziegler G, Monteiro JM, et al. Brain-behaviour modes of covariation in healthy and clinically depressed young people. Scientific Reports 2019;9:1–11. 10.1038/s41598-019-47277-3.

[28] Nicolaisen-Sobesky E, Mihalik A, Kharabian-Masouleh S, Ferreira FS, Hoffstaedter F, Schwender H, et al. A cross-cohort replicable and heritable latent dimension linking behaviour to multi-featured brain structure. Communications Biology 2022;5:1–36. 10.1038/s42003-022-04244-5.

[29] Tian YE, Biase MAD, Mosley PE, Lupton MK, Xia Y, Fripp J, et al. Evaluation of Brain-Body Health in Individuals with Common Neuropsychiatric Disorders. JAMA Psychiatry 2023;80:567–76. 10.1001/jamapsychiatry.2023.0791.

[30] Beyer F, Masouleh SK, Kratzsch J, Schroeter ML, Röhr S, Riedel-Heller SG, et al. A metabolic obesity profile is associated with decreased gray matter volume in cognitively healthy older adults. Frontiers in Aging Neuroscience 2019;10:1–14. 10.3389/fnagi.2019.00202.

[31] Schaare HL, Masouleh SK, Beyer F, Kumral D, Uhlig M, Reinelt JD, et al. Association of peripheral blood pressure with gray matter volume in 19-to 40-year-old adults. Neurology 2019;92:E758–73. 10.1212/WNL.0000000000006947.

[32] Beyer F, Masouleh SK, Huntenburg JM, Lampe L, Luck T, Riedel-Heller SG, et al. Higher body mass index is associated with reduced posterior default mode connectivity in older adults. Human Brain Mapping 2017;38:3502–15. 10.1002/hbm.23605.

[33] Murray S, Tulloch A, Gold MS, Avena NM. Hormonal and neural mechanisms of food reward, eating behaviour and obesity. Nature Reviews Endocrinology 2014;10:540–52. 10.1038/nrendo.2014.91.

[34] Hansen JY, Shafiei G, Markello RD, Smart K, Cox SML, Nørgaard M, et al. Mapping neurotransmitter systems to the structural and functional organization of the human neocortex. Nature Neuroscience 2022;25:1569–81. 10.1038/s41593-022-01186-3.

[35] Markello RD, Hansen JY, Liu Z, Bazinet V, Shafiei G, Suárez LE, et al. neuromaps : structural and functional interpretation of brain maps. Nature Methods 2022. 10.1038/s41592-022-01625-w.

[36] Corlier F, Hafzalla G, Faskowitz J, Kuller LH, Becker JT, Lopez OL, et al. Systemic inflammation as a predictor of brain aging: Contributions of physical activity, metabolic risk, and genetic risk. NeuroImage 2018;172:118–29. 10.1016/j.neuroimage.2017.12.027.

[37] Marsland AL, Gianaros PJ, Kuan DCH, Sheu LK, Krajina K, Manuck SB. Brain morphology links systemic inflammation to cognitive function in midlife adults. Brain, Behavior, and Immunity 2015;48:195–204. 10.1016/j.bbi.2015.03.015.

[38] Miller AA, Spencer SJ. Obesity and neuroinflammation: A pathway to cognitive impairment. Brain, Behavior, and Immunity 2014;42:10–21. 10.1016/j.bbi.2014.04.001.

[39] Patel V, Edison P. Cardiometabolic risk factors and neurodegeneration: a review of the mechanisms underlying diabetes, obesity and hypertension in Alzheimer’s disease. Journal of Neurology, Neurosurgery and Psychiatry 2024;95:581–9. 10.1136/jnnp-2023-332661.

[40] Patel Y, Woo A, Shi S, Ayoub R, Shin J, Botta A, et al. Obesity and the cerebral cortex: Underlying neurobiology in mice and humans. Brain, Behavior, and Immunity 2024;119:637–47. 10.1016/j.bbi.2024.04.033.

[41] Genon S, Eickhoff SB, Kharabian-Masouleh S. Linking interindividual variability in brain structure to behaviour. Nature Reviews Neuroscience 2022;23:307–18. 10.1038/s41583-022-00584-7.

[42] Miller KL, Alfaro-Almagro F, Bangerter NK, Thomas DL, Yacoub E, Xu J, et al. Multimodal population brain imaging in the UK Biobank prospective epidemiological study. Nature Neuroscience 2016;19:1523–36. 10.1038/nn.4393.

[43] Sudlow C, Gallacher J, Allen N, Beral V, Burton P, Danesh J, et al. UK Biobank: An Open Access Resource for Identifying the Causes of a Wide Range of Complex Diseases of Middle and Old Age. PLoS Med 2015;12:e1001779. 10.1371/journal.pmed.1001779.

[44] Alfaro-Almagro F, Jenkinson M, Bangerter NK, Andersson JLR, Sotiropoulos SN, Jbabdi S, et al. Image processing and Quality Control for the first 10,000 brain imaging datasets from UK Biobank. NeuroImage 2018;166:400–24. 10.1016/j.neuroimage.2017.10.034.

[45] Destrieux C, Fischl B, Dale A, Halgren E. Automatic parcellation of human cortical gyri and sulci using standard anatomical nomenclature. NeuroImage 2010;53:1–15. 10.1016/j.neuroimage.2010.06.010.

[46] Sasse L, Nicolaisen-Sobesky E, Dukart J, Eickhoff SB, Götz M, Hamdan S, et al. Overview of leakage scenarios in supervised machine learning. J Big Data 2025;12:135. 10.1186/s40537-025-01193-8.

[47] Hotelling H. Relations between two sets of variates. Biometrika 1936;28:3/4.

[48] Helmer M, Warrington S, Mohammadi-Nejad A-R, Ji JL, Howell A, Rosand B, et al. On stability of Canonical Correlation Analysis and Partial Least Squares with application to brain-behavior associations. BioRxiv 2023. 10.1101/2020.08.25.265546.

[49] Vinod HD. Canonical ridge and econometrics of joint production. Journal of Econometrics 1976;4:147–66. 10.1016/0304-4076(76)90010-5.

[50] Hardoon DR, Szedmak S, Shawe-Taylor J. Canonical correlation analysis: An overview with application to learning methods. Neural Computation 2004;16:2639–64. 10.1162/0899766042321814.

[51] Kapoor S, Narayanan A. Leakage and the reproducibility crisis in machine-learning-based science. Patterns 2023;4. 10.1016/j.patter.2023.100804.

[52] Monteiro JM, Rao A, Shawe-Taylor J, Mourão-Miranda J. A multiple hold-out framework for Sparse Partial Least Squares. Journal of Neuroscience Methods 2016;271:182–94. 10.1016/j.jneumeth.2016.06.011.

[53] Alexander-Bloch AF, Shou H, Liu S, Satterthwaite TD, Glahn DC, Shinohara RT, et al. On testing for spatial correspondence between maps of human brain structure and function. NeuroImage 2018;178:540–51. 10.1016/j.neuroimage.2018.05.070.

[54] Beliveau V, Ganz M, Feng L, Ozenne B, Højgaard L, Fisher PM, et al. A High-Resolution In Vivo Atlas of the Human Brain’s Serotonin System 2017;37:120–8. 10.1523/JNEUROSCI.2830-16.2016.

[55] Savli M, Bauer A, Mitterhauser M, Ding YS, Hahn A, Kroll T, et al. Normative database of the serotonergic system in healthy subjects using multi-tracer PET. NeuroImage 2012;63:447–59. 10.1016/j.neuroimage.2012.07.001.

[56] Dukart J, Holiga Š, Chatham C, Hawkins P, Forsyth A, Mcmillan R, et al. Cerebral blood flow predicts differential neurotransmitter activity 2018:1–11. 10.1038/s41598-018-22444-0.

[57] Lukow PB, Martins D, Veronese M, Vernon AC, McGuire P, Turkheimer FE, et al. Cellular and molecular signatures of in vivo imaging measures of GABAergic neurotransmission in the human brain. Communications Biology 2022;5:1–11. 10.1038/s42003-022-03268-1.

[58] Bedard M, Aghourian M, Legault-denis C, Postuma RB, Soucy J, Gagnon J, et al. Brain Cholinergic Alterations in Idiopathic REM Sleep Behaviour Disorder: A PET Imaging Study with 18F-FEOBV. Sleep Medicine 2018. 10.1016/j.sleep.2018.12.020.

[59] Glasser MF, Coalson TS, Robinson EC, Hacker CD, Harwell J, Yacoub E. A multi-modal parcellation of human cerebral cortex. Nature Publishing Group 2016;536:171–8. 10.1038/nature18933.

[60] Margulies DS, Ghosh SS, Goulas A, Falkiewicz M, Huntenburg JM, Langs G, et al. Situating the default-mode network along a principal gradient of macroscale cortical organization. Proceedings of the National Academy of Sciences of the United States of America 2016;113:12574–9. 10.1073/pnas.1608282113.

[61] Vaishnavi SN, Vlassenko AG, Rundle MM, Snyder AZ, Mintun MA, Raichle ME. Regional aerobic glycolysis in the human brain. Proceedings of the National Academy of Sciences of the United States of America 2010;107:17757–62. 10.1073/pnas.1010459107.

[62] Wey HY, Gilbert TM, Zürcher NR, She A, Bhanot A, Taillon BD, et al. Insights into neuroepigenetics through human histone deacetylase PET imaging. Science Translational Medicine 2016;8. 10.1126/scitranslmed.aaf7551.

[63] Hawrylycz MJ, Lein ES, Guillozet-Bongaarts AL, Shen EH, Ng L, Miller JA, et al. An anatomically comprehensive atlas of the adult human brain transcriptome. Nature 2012;489:391–9. 10.1038/nature11405.

[64] Markello RD, Arnatkeviciute A, Poline B, Fulcher BD. Standardizing workflows in imaging transcriptomics with the abagen toolbox 2021:1–27.

[65] Malén T, Karjalainen T, Isojärvi J, Vehtari A, Bürkner P-C, Putkinen V, et al. Atlas of type 2 dopamine receptors in the human brain: Age and sex dependent variability in a large PET cohort. NeuroImage 2022;255:119149. 10.1016/j.neuroimage.2022.119149.

[66] Sasaki T, Ito H, Kimura Y, Arakawa R, Takano H, Seki C, et al. Quantification of Dopamine Transporter in Human Brain Using PET with^18^ F-FE-PE2I. J Nucl Med 2012;53:1065–73. 10.2967/jnumed.111.101626.

[67] Gallezot J-D, Nabulsi N, Neumeister A, Planeta-Wilson B, Williams WA, Singhal T, et al. Kinetic Modeling of the Serotonin 5-HT_1B_ Receptor Radioligand [^11^ C]P943 in Humans. J Cereb Blood Flow Metab 2010;30:196–210. 10.1038/jcbfm.2009.195.

[68] Gallezot J-D, Planeta B, Nabulsi N, Palumbo D, Li X, Liu J, et al. Determination of receptor occupancy in the presence of mass dose: [^11^ C]GSK189254 PET imaging of histamine H_3_ receptor occupancy by PF-03654746. J Cereb Blood Flow Metab 2017;37:1095–107. 10.1177/0271678X16650697.

[69] Smart K, Cox SML, Scala SG, Tippler M, Jaworska N, Boivin M, et al. Sex differences in [11C]ABP688 binding: a positron emission tomography study of mGlu5 receptors. Eur J Nucl Med Mol Imaging 2019;46:1179–83. 10.1007/s00259-018-4252-4.

[70] Mueller S, Wang D, Fox MD, Yeo BTT, Sepulcre J, Sabuncu MR, et al. Individual Variability in Functional Connectivity Architecture of the Human Brain. Neuron 2013;77:586–95. 10.1016/j.neuron.2012.12.028.

[71] Reardon PK, Seidlitz J, Vandekar S, Liu S, Patel R, Lalonde FM, et al. Normative brain size variation and brain shape diversity in humans 2018;1227:4–8.

[72] DuBois JM, Rousset OG, Rowley J, Porras-Betancourt M, Reader AJ, Labbe A, et al. Characterization of age/sex and the regional distribution of mGluR5 availability in the healthy human brain measured by high-resolution [11C]ABP688 PET. Eur J Nucl Med Mol Imaging 2016;43:152–62. 10.1007/s00259-015-3167-6.

[73] Galovic M, Al-Diwani A, Vivekananda U, Torrealdea F, Erlandsson K, Fryer TD, et al. *In vivo* NMDA receptor function in people with NMDA receptor antibody encephalitis 2021. 10.1101/2021.12.04.21267226.

[74] Galovic M, Erlandsson K, Fryer TD, Hong YT, Manavaki R, Sari H, et al. Validation of a combined image derived input function and venous sampling approach for the quantification of [18F]GE-179 PET binding in the brain. NeuroImage 2021;237:118194. 10.1016/j.neuroimage.2021.118194.

[75] McGinnity CJ, Hammers A, Riaño Barros DA, Luthra SK, Jones PA, Trigg W, et al. Initial Evaluation of^18^ F-GE-179, a Putative PET Tracer for Activated *N* -Methyl d-Aspartate Receptors. J Nucl Med 2014;55:423–30. 10.2967/jnumed.113.130641.

[76] Kantonen T, Karjalainen T, Isojärvi J, Nuutila P, Tuisku J, Rinne J, et al. Interindividual variability and lateralization of μ-opioid receptors in the human brain: Individual differences in the μ-opioid receptor system. NeuroImage 2020;217. 10.1016/j.neuroimage.2020.116922.

[77] Turtonen O, Saarinen A, Nummenmaa L, Tuominen L, Tikka M, Armio RL, et al. Adult Attachment System Links With Brain Mu Opioid Receptor Availability In Vivo. Biological Psychiatry: Cognitive Neuroscience and Neuroimaging 2021;6:360–9. 10.1016/j.bpsc.2020.10.013.

[78] Vijay A, Cavallo D, Goldberg A, De Laat B, Nabulsi N, Huang Y, et al. PET imaging reveals lower kappa opioid receptor availability in alcoholics but no effect of age. Neuropsychopharmacol 2018;43:2539–47. 10.1038/s41386-018-0199-1.

[79] Laurikainen H, Tuominen L, Tikka M, Merisaari H, Armio R-L, Sormunen E, et al. Sex difference in brain CB1 receptor availability in man. NeuroImage 2019;184:834–42. 10.1016/j.neuroimage.2018.10.013.

[80] Normandin MD, Zheng M-Q, Lin K-S, Mason NS, Lin S-F, Ropchan J, et al. Imaging the Cannabinoid CB1 Receptor in Humans with [^11^ C] OMAR: Assessment of Kinetic Analysis Methods, Test–Retest Reproducibility, and Gender Differences. J Cereb Blood Flow Metab 2015;35:1313–22. 10.1038/jcbfm.2015.46.

[81] Xu T, Nenning K-H, Schwartz E, Hong S-J, Vogelstein JT, Goulas A, et al. Cross-species functional alignment reveals evolutionary hierarchy within the connectome. NeuroImage 2020;223:117346. 10.1016/j.neuroimage.2020.117346.

[82] Sydnor VJ, Larsen B, Bassett DS, Alexander-Bloch A, Fair DA, Liston C, et al. Neurodevelopment of the association cortices: Patterns, mechanisms, and implications for psychopathology. Neuron 2021;109:2820–46. 10.1016/j.neuron.2021.06.016.

[83] Tai XY, Veldsman M, Lyall DM, Littlejohns TJ, Langa KM, Husain M, et al. Cardiometabolic multimorbidity, genetic risk, and dementia: a prospective cohort study. The Lancet Healthy Longevity 2022;3:e428–36. 10.1016/S2666-7568(22)00117-9.

[84] Vergoossen LWM, Jansen JFA, Backes WH, Schram MT. Cardiometabolic determinants of early and advanced brain alterations: Insights from conventional and novel MRI techniques. Neuroscience and Biobehavioral Reviews 2020;115:308–20. 10.1016/j.neubiorev.2020.04.001.

[85] Beck D, Lange AMG de, Pedersen ML, Alnæs D, Maximov II, Voldsbekk I, et al. Cardiometabolic risk factors associated with brain age and accelerate brain ageing. Human Brain Mapping 2021;43:700–20. 10.1002/hbm.25680.

[86] Pillinger T, D’Ambrosio E, McCutcheon R, Howes OD. Is psychosis a multisystem disorder? A meta-review of central nervous system, immune, cardiometabolic, and endocrine alterations in first-episode psychosis and perspective on potential models. Molecular Psychiatry 2019;24:776–94. 10.1038/s41380-018-0058-9.

[87] Schaare HL, Kumral D, Uhlig M, Lemcke L, Villringer A. Associations between mental health, blood pressure and the development of hypertension. Nature Communications 2023;14:1–17. 10.1038/s41467-023-37579-6.

[88] Gusev E, Sarapultsev A. Interplay of G-Proteins and Serotonin in the Neuroimmunoinflammatory Model of Chronic Stress and Depression: A Narrative Review. Current Pharmaceutical Design 2024;30:180–214. 10.2174/0113816128285578231218102020.

[89] O’Brien PD, Hinder LM, Callaghan BC, Feldman EL. Neurological consequences of obesity. The Lancet Neurology 2017;16:465–77. 10.1016/S1474-4422(17)30084-4.

[90] Anisman H, Merali Z, Hayley S. Neurotransmitter, peptide and cytokine processes in relation to depressive disorder: Comorbidity between depression and neurodegenerative disorders. Progress in Neurobiology 2008;85:1–74. 10.1016/j.pneurobio.2008.01.004.

[91] Iakunchykova O, Leonardsen EH, Wang Y. Genetic evidence for causal effects of immune dysfunction in psychiatric disorders: where are we? Transl Psychiatry 2024;14:63. 10.1038/s41398-024-02778-2.

[92] Leonard BE. Impact of inflammation on neurotransmitter changes in major depression: An insight into the action of antidepressants. Progress in Neuro-Psychopharmacology and Biological Psychiatry 2014;48:261–7. 10.1016/j.pnpbp.2013.10.018.

[93] Nguyen TT, Hulme J, Vo TK, Vo GV. The Potential Crosstalk Between the Brain and Visceral Adipose Tissue in Alzheimer’s Development. Neurochemical Research 2022;47:1503–12. 10.1007/s11064-022-03569-1.

[94] Iakunchykova O, Schirmer H, Roe JM, Sørensen Ø, Wilsgaard T, Hopstock LA, et al. Longitudinal and concurrent C-reactive protein and diet associations with cognitive function in the population-based Tromsø study. Journal of Alzheimer’s Disease 2025;104:403–13. 10.1177/13872877251317624.

[95] Cotter DL, Ahmadi H, Cardenas-Iniguez C, Bottenhorn KL, Gauderman WJ, McConnell R, et al. Exposure to multiple ambient air pollutants changes white matter microstructure during early adolescence with sex-specific differences. Commun Med 2024;4:155. 10.1038/s43856-024-00576-x.

[96] Kim H, Noh J, Noh Y, Oh SS, Koh S-B, Kim C. Gender Difference in the Effects of Outdoor Air Pollution on Cognitive Function Among Elderly in Korea. Front Public Health 2019;7:375. 10.3389/fpubh.2019.00375.

[97] Kuźma Ł, Struniawski K, Pogorzelski S, Bachórzewska-Gajewska H, Dobrzycki S. Gender Differences in Association between Air Pollution and Daily Mortality in the Capital of the Green Lungs of Poland–Population-Based Study with 2,953,000 Person-Years of Follow-Up. JCM 2020;9:2351. 10.3390/jcm9082351.

[98] Mo S, Wang Y, Peng M, Wang Q, Zheng H, Zhan Y, et al. Sex disparity in cognitive aging related to later-life exposure to ambient air pollution. Science of The Total Environment 2023;886:163980. 10.1016/j.scitotenv.2023.163980.

[99] Oiamo T, Luginaah I. Extricating Sex and Gender in Air Pollution Research: A Community-Based Study on Cardinal Symptoms of Exposure. IJERPH 2013;10:3801–17. 10.3390/ijerph10093801.

[100] van Galen KA, Horst KW ter, Serlie MJ. Serotonin, food intake, and obesity. Obesity Reviews 2021;22:1–13. 10.1111/obr.13210.

[101] Voigt JP, Fink H. Serotonin controlling feeding and satiety. Behavioural Brain Research 2015;277:14–31. 10.1016/j.bbr.2014.08.065.

[102] Cordeiro LMS, Rabelo PCR, Moraes MM, Teixeira-Coelho F, Coimbra CC, Wanner SP, et al. Physical exercise-induced fatigue: The role of serotonergic and dopaminergic systems. Brazilian Journal of Medical and Biological Research 2017;50:1–13. 10.1590/1414-431X20176432.

[103] Frank GKW. Advances from neuroimaging studies in eating disorders. CNS Spectrums 2015;20:391–400. 10.1017/S1092852915000012.

[104] Penninx BWJH, Lamers F, Jansen R, Berk M, Khandaker GM, De Picker L, et al. Immuno-metabolic depression: from concept to implementation. The Lancet Regional Health - Europe 2025;48:101166. 10.1016/j.lanepe.2024.101166.

[105] Matt SM, Gaskill PJ. Where Is Dopamine and how do Immune Cells See it?: Dopamine-Mediated Immune Cell Function in Health and Disease. Journal of Neuroimmune Pharmacology 2020;15:114–64. 10.1007/s11481-019-09851-4.

