## Supplementary materials for "Cardiometabolic health and physical robustness map onto distinct patterns of brain structure and neurotransmitter systems"

**Captions**

1. Supplementary methods

1.1 Sex-specific analyses.
1.2 Comparison of latent dimensions with demographics.
1.3 Latent dimensions linking risk factors to subcortical and cerebellar volumes.

2. Supplementary results

2.1 Latent dimensions linking cardiometabolic health to raw cortical thickness and raw grey matter volume.
2.2 Latent dimensions linking cardiometabolic health to proportional cortical thickness and proportional grey matter volume.
2.3 Latent dimensions linking cardiometabolic health to corrected cortical thickness and corrected grey matter volume.
2.4 Latent dimension of cardiometabolic health and demographics.
2.5 Latent dimensions linking physical robustness to raw cortical thickness and raw grey matter volume.
2.6 Latent dimensions linking physical robustness to proportional cortical thickness and proportional grey matter volume.
2.7 Latent dimensions linking physical robustness to corrected cortical thickness and corrected grey matter volume.
2.8 Latent dimension of physical robustness and demographics.

2.9 Latent dimensions linking risk factors to subcortical and cerebellar volumes

3. Supplementary tables

Table S1. Variables of risk factors used in this study.
Table S2. Demographic variables used in this study.
Table S3. Brain maps included in neuromaps that were compared with the latent dimension.
Table S4. Statistics of the latent dimensions for raw cortical thickness.
Table S5. Statistics of the latent dimensions for raw grey matter volume.
Table S6. Statistics of the latent dimensions for proportional cortical thickness.
Table S7. Statistics of the latent dimensions for proportional grey matter volume.
Table S8. Statistics of the latent dimensions for corrected cortical thickness.
Table S9. Statistics of the latent dimensions for corrected grey matter volume.
Table S10. Association between the brain loadings of the latent dimension of cardiometabolic health and neuromaps.
Table S11. Association between the brain loadings of the latent dimension of physical robustness and neuromaps.
Table S12. Statistics of the latent dimensions for subcortical and cerebellar volumes.

4. Supplementary figures

Figure S1. Distribution of risk factors. Part 1.
Figure S2. Distribution of risk factors. Part 2.
Figure S3. Correlation matrix of risk factors in women.
Figure S4. Correlation matrix of risk factors in men.
Figure S5. Latent dimension of cardiometabolic health.

Figure S6. Model optimization for the latent dimension of cardiometabolic health for raw cortical thickness in the main sample.
Figure S7. Model optimization for the latent dimension of cardiometabolic health for raw grey matter volume in the main sample.
Figure S8. Comparison of risk factor loadings and brain loadings for the cardiometabolic health latent dimension across samples and brain structural measures.
Figure S9. Loadings of the latent dimension of cardiometabolic health for raw cortical thickness in women.
Figure S10. Loadings of the latent dimension of cardiometabolic health for raw cortical thickness in men.
Figure S11. Loadings of the latent dimension of cardiometabolic health for raw grey matter volume in women.
Figure S12. Loadings of the latent dimension of cardiometabolic health for raw grey matter volume in men.
Figure S13. Loadings of the latent dimension of cardiometabolic health for proportional cortical thickness in the main sample.
Figure S14. Loadings of the latent dimension of cardiometabolic health for proportional cortical thickness in women.
Figure S15. Loadings of the latent dimension of cardiometabolic health for proportional cortical thickness in men.
Figure S16. Loadings of the latent dimension of cardiometabolic health for proportional grey matter volume in the main sample.
Figure S17. Loadings of the latent dimension of cardiometabolic health for proportional grey matter volume in women.
Figure S18. Loadings of the latent dimension of cardiometabolic health for proportional grey matter volume in men.
Figure S19. Loadings of the latent dimension of cardiometabolic health for corrected cortical thickness in the main sample.
Figure S20. Loadings of the latent dimension of cardiometabolic health for corrected cortical thickness in women.
Figure S21. Loadings of the latent dimension of cardiometabolic health for corrected cortical thickness in men.
Figure S22. Loadings of the latent dimension of cardiometabolic health for corrected grey matter volume in the main sample.
Figure S23. Loadings of the latent dimension of cardiometabolic health for corrected grey matter volume in women.
Figure S24. Loadings of the latent dimension of cardiometabolic health for corrected grey matter volume in men.
Figure S25. Latent dimension of physical robustness.
Figure S26. Model optimization for the latent dimension of physical robustness for raw cortical thickness in the sample of men.
Figure S27. Model optimization for the latent dimension of physical robustness for raw grey matter volume in the main sample.
Figure S28. Loadings of the latent dimension of physical robustness for raw cortical thickness in women.
Figure S29. Loadings of the latent dimension of physical robustness for raw grey matter volume in women.
Figure S30. Loadings of the latent dimension of physical robustness for raw grey matter volume in men.
Figure S31. Comparison of risk factor loadings and brain loadings for the physical robustness latent dimension across samples and brain structural measures.
Figure S32. Loadings of the latent dimension of physical robustness for proportional cortical thickness in men.
Figure S33. Loadings of the latent dimension of physical robustness for proportional grey matter volume in the main sample.
Figure S34. Loadings of the latent dimension of physical robustness for proportional grey matter volume in women.
Figure S35. Loadings of the latent dimension of physical robustness for proportional grey matter volume in men.
Figure S36. Loadings of the latent dimension of physical robustness for corrected cortical thickness in men.
Figure S37. Loadings of the latent dimension of physical robustness for corrected cortical thickness in women.
Figure S38. Loadings of the latent dimension of physical robustness for corrected grey matter volume in the main sample.
Figure S39. Loadings of the latent dimension of physical robustness for corrected grey matter volume in women.
Figure S40. Loadings of the latent dimension of physical robustness for corrected grey matter volume in men.
Figure S41. Association of brain structural loadings with brain maps for the latent dimension of cardiometabolic health.

Figure S42. Association of brain structural loadings with brain maps for the latent dimension of physical robustness.
Figure S43. Loadings of the first latent dimension linking risk factors to subcortical and cerebellar volumes.
Figure S44. Loadings of the second latent dimension linking risk factors to subcortical and cerebellar volumes.
Figure S45. Loadings of the third latent dimension linking risk factors to subcortical and cerebellar volumes.
Figure S46. Loadings of the fourth latent dimension linking risk factors to subcortical and cerebellar volumes.
Figure S47. Loadings of the fifth latent dimension linking risk factors to subcortical and cerebellar volumes.
Figure S48. Comparison of risk factor loadings between latent dimensions yielded with subcortex-cerebellum and with CT and GMV in cortex.

### 1. S1 appendix: Supplementary methods

#### *1.1 Sex-specific analyses*

In the main paper, we present analyses in a sample including both, men and women (the main/mixed sample). In addition, we performed sex-specific analyses because previous works have shown sex differences in the association between brain structure and risk factors (1,2) and also in order to avoid sex-bias in the results (3,4). The existence of a sex bias in human neuroscience is recognized, with the consequence that much of our knowledge in the field is based on men, and do not or might not generalize to women (5). To perform the sex-specific analyses, we run our pipeline independently in the subsample of women and in the subsample of men.

#### *1.2 Comparison of latent dimensions with demographics*

To assess the role of demographic variables in the latent dimensions, we performed Spearman correlation between the risk factors scores and household income, employment status, and education (Table S2). Multiple comparisons were corrected using the Bonferroni method.

#### *1.3 Latent dimensions linking risk factors to subcortical and cerebellar volumes*

Using the same framework described in the main paper for the analyses with cortical data (CT and GMV), we also analyzed latent dimensions linking the same set of risk factors with subcortical and cerebellar volumes in the main sample. Subcortical and cerebellar volumes were estimated with the FMRIB pipeline and parceled with the Harvard-Oxford and Diedrichsen atlases (6)**.** Age, site and sex were regressed out avoiding data leakage (7).

### 2. S2. Appendix: Supplementary results

#### *2.1 Latent dimensions linking cardiometabolic health to raw cortical thickness and raw grey matter volume*

When analyzed in association with raw CT, the latent dimension of cardiometabolic health was significant in the sample of women (Figure S9, Table S4) and in the sample of men (Figure S10, Table S4).

When analyzed in association with raw GMV, the latent dimension of cardiometabolic health was significant in the sample of women (Figure S11, Table S5) and in the sample of men (Figure S12, Table S5).

Interestingly, the risk factors loadings associated with cardiometabolic health were similar when yielded with raw CT or with raw GMV, while the CT and GMV loadings were not significantly associated (Figure S8).

#### *2.2 Latent dimensions linking cardiometabolic health to proportional cortical thickness and proportional grey matter volume*

When analyzed in association with proportional CT, the latent dimension of cardiometabolic health was significant in the main/mixed sample (Figure S13, Table S6), as well as in the sample of women (Figure S14, Table S6) and in the sample of men (Figure S15, Table S6).

When analyzed in association with proportional GMV, the latent dimension of cardiometabolic health was significant in the main/mixed sample (Figure S16, Table S7), as well as in the sample of women (Figure S17, Table S7) and in the sample of men (Figure S18, Table S7).

Interestingly, the risk factors loadings associated with cardiometabolic health were similar when yielded with proportional CT or with proportional GMV, while the CT and GMV loadings were not significantly associated (Figure S8).

#### *2.3 Latent dimensions linking cardiometabolic health to corrected cortical thickness and corrected grey matter volume*

When analyzed in association with corrected CT, the latent dimension of cardiometabolic health was significant in the main/mixed sample (Figure S19, Table S8), as well as in the sample of women (Figure S20, Table S8) and in the sample of men (Figure S21, Table S8).

When analyzed in association with corrected GMV, the latent dimension of cardiometabolic health was significant in the main/mixed sample (Figure S22, Table S9), as well as in the sample of women (Figure S23, Table S9) and in the sample of men (Figure S24, Table S9).

Interestingly, the risk factors loadings associated with cardiometabolic health were similar when yielded with corrected CT or with corrected GMV, while the CT and GMV loadings were not significantly associated (Figure S8).

#### *2.4 Latent dimension of cardiometabolic health and demographics*

Our results showed that qualifications (education) and household income were significantly associated with the risk factor scores of the latent dimension of cardiometabolic health when yielded with raw CT (qualifications: $r = 0.15$, $p < 0.001$; income: $r = 0.07$, $p < 0.001$) and raw GMV (qualifications: $r = 0.13$, $p < 0.001$; income: $r = 0.06$, $p < 0.001$) in the main/mixed subsample. This indicates that individuals who score high on the positive pole of risk factors (that are characterized for example by higher physical activity) and hence who score very low on the negative pole of risk factors (that are characterized for example with lower BMI and body size) are associated with higher qualifications and higher household income.

Analyses could not be performed for ethnic background and some employment dummy variables (those corresponding to values 3-7) because of imbalanced distribution.

#### *2.5 Latent dimensions linking physical robustness to raw cortical thickness and raw grey matter volume*

When analyzed in association with raw CT, the latent dimension of physical robustness was significant in the sample of men (Figure 4 in main paper, Table S4). However, in the sample of women it was significant in only one split ($r = 0.103$, $p = 0.01$) out of five ($r_{range} = 0.013\text{-}0.027$, $p_{range} = 0.384\text{-}0.999$). Considering that, and that the loadings were unstable for most of the risk factors loadings and brain regions in the sample of women (Figure S28), for the latent dimension of physical robustness and cortical thickness we focus on the sample of men.

When analyzed in association with raw GMV, the latent dimension of physical robustness was significant in the sample of women (Figure S29, Table S5) and in the sample of men (Figure S30, Table S5).

Interestingly, the risk factors loadings associated with physical robustness were similar when yielded with raw CT or with raw GMV, while the CT and GMV loadings were not significantly associated (Figure S31).

*2.6 Latent dimensions linking physical robustness to proportional cortical thickness and proportional grey matter volume*

When analyzed in association with proportional CT, the latent dimension of physical robustness was significant in the sample of men (Figure S32, Table S6). However, in the sample of women it was not significant ($r_{range}$ = 0.003-0.060, $p_{range}$ = 0.220-0.999). For this reason, for the latent dimension of physical robustness and cortical thickness we focus on the sample of men.

When analyzed in association with proportional GMV, the latent dimension of physical robustness was significant in the main/mixed sample (Figure S33, Table S7), as well as in the sample of women (Figure S34, Table S7) and in the sample of men (Figure S35, Table S7).

Interestingly, the risk factors loadings associated with physical robustness were similar when yielded with proportional CT or with proportional GMV, while the CT and GMV loadings were not significantly associated (Figure S31).

*2.7 Latent dimensions linking physical robustness to corrected cortical thickness and corrected grey matter volume*

When analyzed in association with corrected CT, the latent dimension of physical robustness was significant in the sample of men (Figure S36, Table S8). However, in the sample of women it was significant in only one split ($r$ = 0.108, $p$ = 0.01) out of five ($r_{range}$ = 0.003-0.047, $p_{range}$ = 0.469-0.999). Considering that, and that the loadings were quite unstable (Figure S37), for the latent dimension of physical robustness and cortical thickness we focus on the sample of men.

When analyzed in association with corrected GMV, the latent dimension of physical robustness was significant in the main/mixed sample (Figure S38, Table S9), as well as in the sample of women (Figure S39, Table S9) and in the sample of men (Figure S40, Table S9).

Interestingly, the risk factors loadings associated with physical robustness were similar when yielded with corrected CT or with corrected GMV, while the CT and GMV loadings were not significantly associated (Figure S31).

*2.8 Latent dimension of physical robustness and demographics*

Our results showed household income was significantly associated with the risk factor scores of the latent dimension of physical robustness when yielded with raw CT in the sample of men ($r = 0.05$, $p = 0.039$) and with raw GMV in the main/mixed sample ($r = 0.08$, $p < 0.001$). Also, qualifications (education) was associated with the risk factor scores when yielded with raw GMV in the main/mixed sample ($r = 0.10$, $p < 0.001$).

Analyses could not be performed for ethnic background and some employment dummy variables (those corresponding to values 3-7) because of imbalanced distribution.

*2.9 Latent dimensions linking risk factors to subcortical and cerebellar volumes*

The analysis of subcortical and cerebellar data yielded five significant latent dimensions (Figures S43-S47, Table S12). Of those, the risk factor loadings of the first subcortex-cerebellum latent dimension showed significant associations with the loadings of both, the latent dimension of cardiometabolic health and the latent dimension of physical robustness yielded with the cortical data (CT and GMV) (Figure S48). The second latent dimension yielded with subcortex-cerebellum also showed significant associations with the latent dimension of cardiometabolic health. In other words, this first latent dimension that explains interindividual variability in non-cortical regions seems to combine both cardiometabolic health and physical robustness factors. It relates to variability across a wide range of non-cortical regions, such as the amygdala, the hippocampus, the thalamus and several cerebellar regions, but to a lesser extend in basal ganglia regions. This suggests that the set of factors that explain variability in subcortical regions in an aging population may be importantly more complex than the set of factors that relate cortical interindividual variability. Accordingly, we would encourage future studies to further specifically investigate factors that may affect subcortical structures across the lifespan, instead of considering cortical and subcortical regions globally.

### 3. S3 appendix: Supplementary tables

**Table S1. Variables of risk factors used in this study.**

| UDI | Label | Category | Modified or reversed |
|---|---|---|---|
| 46-2.0 | Hand grip strength (left) | Hand grip strength | |
| 47-2.0 | Hand grip strength (right) | Hand grip strength | |
| 48-2.0 | Waist circumference | Body size measures | |
| 49-2.0 | Hip circumference | Body size measures | |
| 102-0.0 | Pulse rate, automated reading | Blood pressure | |
| 135-0.0 | Number of self-reported non-cancer illnesses | Medical conditions | |
| 136-0.0 | Number of operations, self-reported | Operations | |
| 137-0.0 | Number of treatments/medications taken | Medications | |
| 864-2.0 | Number of days/weeks walked 10+ minutes | Physical activity | |
| 884-2.0 | Number of days/weeks of moderate physical activity 10+ minutes | Physical activity | |
| 904-2.0 | Number of days/weeks of vigorous physical activity 10+ minutes | Physical activity | |

| | | | |
|---|---|---|---|
| 924-2.0 | Usual walking pace | Physical activity | |
| 943-2.0 | Frequency of stair climbing in last 4 weeks | Physical activity | |
| 1070-2.0 | Time spent watching television (TV) | Physical activity | Modified (-10 replaced by 0) |
| 1080-2.0 | Time spent using computer | Physical activity | Modified (-10 replaced by 0) |
| 1090-2.0 | Time spent driving | Physical activity | Modified (-10 replaced by 0) |
| 1160-2.0 | Sleep duration | Sleep | |
| 1170-2.0 | Getting up in morning | Sleep | |
| 1190-2.0 | Nap during day | Sleep | |
| 1200-2.0 | Sleeplessness / insomnia | Sleep | |
| 1249-2.0 | Past tobacco smoking | Smoking | Reversed ([1,2,3,4] replaced by [14,13,12,11]) |
| 1289-2.0 | Cooked vegetable intake | Diet | Modified (-10 replaced by 0) |
| 1299-2.0 | Salad / raw vegetable intake | Diet | Modified (-10 replaced by 0) |
| 1309-2.0 | Fresh fruit intake | Diet | Modified (-10 replaced by 0) |

| | | | |
|---|---|---|---|
| 1319-2.0 | Dried fruit intake | Diet | Modified (-10 replaced by 0) |
| 1329-2.0 | Oily fish intake | Diet | |
| 1339-2.0 | Non-oily fish intake | Diet | |
| 1349-2.0 | Processed meat intake | Diet | |
| 1359-2.0 | Poultry intake | Diet | |
| 1369-2.0 | Beef intake | Diet | |
| 1379-2.0 | Lamb/mutton intake | Diet | |
| 1389-2.0 | Pork intake | Diet | |
| 1408-2.0 | Cheese intake | Diet | |
| 1438-2.0 | Bread intake | Diet | Modified (-10 replaced by 0) |
| 1458-2.0 | Cereal intake | Diet | Modified (-10 replaced by 0) |
| 1478-2.0 | Salt added to food | Diet | |
| 1488-2.0 | Tea intake | Diet | Modified (-10 replaced by 0) |
| 1498-2.0 | Coffee intake | Diet | Modified (-10 replaced by 0) |
| 1528-2.0 | Water intake | Diet | Modified (-10 replaced by 0) |

| | | | |
|---|---|---|---|
| 1538-2.0 | Major dietary changes in the last 5 years | Diet | Modified (2 replaced by 1) |
| 1548-2.0 | Variation in diet | Diet | |
| 1558-2.0 | Alcohol intake frequency. | Alcohol | Reversed ([1,2,3,4,5,6] replaced by [16,15,14,13,12,11]) |
| 1618-2.0 | Alcohol usually taken with meals | Alcohol | Modified (1, 0, -6 replaced by 12,10,22 respectively) |
| 1628-2.0 | Alcohol intake versus 10 years previously | Alcohol | Reversed ([1,2,3] replaced by [12,11,10]) |
| 2178-2.0 | Overall health rating | General health | Reversed ([1,2,3,4] replaced by [14,13,12,11]) |
| 2306-2.0 | Weight change compared with 1 year ago | General health | Modified ([0,2,3] replaced by [11, 12, 10]) |
| 2492-2.0 | Taking other prescription medications | Medication | |
| 4079-0.0 | Diastolic blood pressure, automated reading | Blood pressure | |
| 4080-0.0 | Systolic blood pressure, automated reading | Blood pressure | |
| 20116-2.0 | Smoking status | Smoking | |
| 20160-2.0 | Ever smoked | Smoking | |

| | | |
|---|---|---|
| 21001-2.0 | Body mass index (BMI) | Body size measures |
| 21003-2.0 | Age when attended assessment centre | Reception |
| 23099-2.0 | Body fat percentage | Body composition by impedance |
| 23100-2.0 | Whole body fat mass | Body composition by impedance |
| 23101-2.0 | Whole body fat-free mass | Body composition by impedance |
| 23102-2.0 | Whole body water mass | Body composition by impedance |
| 23105-2.0 | Basal metabolic rate | Body composition by impedance |
| 23106-2.0 | Impedance of whole body | Body composition by impedance |
| 24003-0.0 | Nitrogen dioxide air pollution; 2010 | Residential air pollution |
| 24004-0.0 | Nitrogen oxides air pollution; 2010 | Residential air pollution |
| 24005-0.0 | Particulate matter air pollution (pm10); 2010 | Residential air pollution |

| | | | |
|---|---|---|---|
| 24006-0.0 | Particulate matter air pollution (pm2.5); 2010 | Residential air pollution | |
| 24007-0.0 | Particulate matter air pollution (pm2.5) absorbance; 2010 | Residential air pollution | |
| 24008-0.0 | Particulate matter air pollution 2.5-10um; 2010 | Residential air pollution | |
| 24011-0.0 | Traffic intensity on the nearest major road | Residential air pollution | |
| 24018-0.0 | Nitrogen dioxide air pollution; 2007 | Residential air pollution | |
| 24019-0.0 | Particulate matter air pollution (pm10); 2007 | Residential air pollution | |
| 46-2.0 | Hand grip strength (left) | Hand grip strength | |
| 54-2.0 | UK Biobank assessment centre (site) | Reception | Dummy |
| 1428-2.0 | Spread type | Diet | Dummy |

Some scores were reversed to facilitate interpretability, represented as dummy variables, or modified to create ordinal variables.

**Table S2. Demographic variables used in this study.**

| **UDI** | **Label** | **Category** | **Modified or reversed** |
|---|---|---|---|
| 738-2.0 | Average total household income before tax | Household | |

| | | | |
|---|---|---|---|
| 21000-0.0 | Ethnic background | Ethnicity | |
| 6138-2.* | Qualifications | Education | Modified and reversed: [1,2,3,4,5,6] were replaced by [16,15,14,14,None,15]. Original value 5 was replaced by None in order to create ordinal variable. |
| 6142-2.* | Current employment status | Employment | Dummy |

**Table S3. Brain maps included in neuromaps that were compared with the latent dimension.**

| | |
|---|---|
| ('abagen', 'genepc1', 'fsaverage', '10k') | PC1 of genes in the Allen Human Brain Atlas |
| ('aghourian2017', 'feobv', 'MNI152', '1mm') | PET tracer binding (SUVR) to VAChT (acetylcholine transporter) |
| ('alarkurtti2015', 'raclopride', 'MNI152', '3mm') | PET tracer binding (BPnd) to D2 (dopamine receptor) |
| ('bedard2019', 'feobv', 'MNI152', '1mm') | PET tracer binding (SUVR) to VAChT (acetylcholine transporter) |
| ('beliveau2017', 'az10419369', 'fsaverage', '164k') | PET tracer binding (Bmax) to 5-HT1b (serotonin receptor) |
| ('beliveau2017', 'cimbi36', 'fsaverage', '164k') | PET tracer binding (Bmax) to 5-HT2a (serotonin receptor) |
| ('beliveau2017', 'cumi101', 'fsaverage', '164k') | PET tracer binding (Bmax) to 5-HT1a (serotonin receptor) |
| ('beliveau2017', 'dasb', 'fsaverage', '164k') | PET tracer binding (Bmax) to 5-HTT (serotonin transporter) |
| ('beliveau2017', 'sb207145', 'fsaverage', '164k') | PET tracer binding (Bmax) to 5-HT4 (serotonin receptor) |

| | |
|---|---|
| ('ding2010', 'mrb', 'MNI152', '1mm') | PET tracer binding (BPnd) to NET (norepinephrine transporter) |
| ('dubois2015', 'abp688', 'MNI152', '1mm') | PET tracer binding (BPnd) to mGluR5 (glutamate receptor) |
| ('dukart2018', 'flumazenil', 'MNI152', '3mm') | PET tracer binding (BPnd) to GABAa (gaba receptor) |
| ('dukart2018', 'fpcit', 'MNI152', '3mm') | SPECT tracer binding (SUVR) to DAT (dopamine transporter) |
| ('fazio2016', 'madam', 'MNI152', '3mm') | PET tracer binding (BPnd) to 5-HTT (serotonin transporter) |
| ('finnema2016', 'ucbj', 'MNI152', '1mm') | PET tracer binding (BPnd) to SV2A (synaptic vesicle glycoprotein 2A, a synapse marker) |
| ('gallezot2010', 'p943', 'MNI152', '1mm') | PET tracer binding (BPnd) to 5-HT1b (serotonin receptor) |
| ('gallezot2017', 'gsk189254', 'MNI152', '1mm') | PET tracer binding (Vt) to H3 (histamine receptor) |
| ('galovic2021', 'ge179', 'MNI152', '1mm') | PET tracer binding (Vt) to NMDA (glutamate receptor) |
| ('hcps1200', 'megalpha', 'fsLR', '4k') | MEG alpha (8-12 Hz) power distribution from the Human Connectome Project S1200 release |
| ('hcps1200', 'megbeta', 'fsLR', '4k') | MEG beta (15-29 Hz) power distribution from the Human Connectome Project S1200 release |
| ('hcps1200', 'megdelta', 'fsLR', '4k') | MEG delta (2-4 Hz) power distribution from the Human Connectome Project S1200 release |
| ('hcps1200', 'meggamma1', 'fsLR', '4k') | MEG low gamma (30-59 Hz) power distribution from the Human Connectome Project S1200 release |
| ('hcps1200', 'meggamma2', 'fsLR', '4k') | MEG high gamma (60-90 Hz) power distribution from the Human Connectome Project S1200 release |

| | |
|---|---|
| ('hcps1200', 'megtheta', 'fsLR', '4k') | MEG theta (5-7 Hz) power distribution from the Human Connectome Project S1200 release |
| ('hcps1200', 'megtimescale', 'fsLR', '4k') | MEG intrinsic timescale from the Human Connectome Project S1200 release |
| ('hcps1200', 'myelinmap', 'fsLR', '32k') | MRI T1w/T2w ratio from the Human Connectome Project S1200 release |
| ('hcps1200', 'thickness', 'fsLR', '32k') | MRI cortical thickness from the Human Connectome Project S1200 release |
| ('hesse2017', 'methylreboxetine', 'MNI152', '3mm') | PET tracer binding (BPnd) to NET (norepinephrine transporter) |
| ('hillmer2016', 'flubatine', 'MNI152', '1mm') | PET tracer binding (Vt) to a4b2 (acetylcholine receptor) |
| ('jaworska2020', 'fallypride', 'MNI152', '1mm') | PET tracer binding (BPnd) to D2 (dopamine receptor) |
| ('kaller2017', 'sch23390', 'MNI152', '3mm') | PET tracer binding (BPnd) to D1 (dopamine receptor) |
| ('kantonen2020', 'carfentanil', 'MNI152', '3mm') | PET tracer binding (BPnd) to MOR (mu-opioid receptor) |
| ('kim2020', 'ps13', 'MNI152', '2mm') | PET tracer binding (Vt) to COX-1 (cyclooxygenase-1) |
| ('laurikainen2018', 'fmpepd2', 'MNI152', '1mm') | PET tracer binding (Vt) to CB1 (cannabinoid receptor) |
| ('lois2018', 'pbr28', 'MNI152', '2mm') | PET tracer binding (SUVR) to TSPO (translocator protein) |
| ('lukow2022', 'ro154513', 'MNI152', '2mm') | PET tracer binding (BPnd) to GABAa receptor, alpha5 subunit |
| ('malen2022', 'raclopride', 'MNI152', '2mm') | PET tracer binding (BPnd) to D2 (dopamine receptor) |
| ('margulies2016', 'fcgradient01', 'fsLR', '32k') | Diffusion map embedding gradient 1 of group-averaged functional connectivity |

| | |
|---|---|
| ('margulies2016', 'fcgradient02', 'fsLR', '32k') | Diffusion map embedding gradient 2 of group-averaged functional connectivity |
| ('margulies2016', 'fcgradient03', 'fsLR', '32k') | Diffusion map embedding gradient 3 of group-averaged functional connectivity |
| ('margulies2016', 'fcgradient04', 'fsLR', '32k') | Diffusion map embedding gradient 4 of group-averaged functional connectivity |
| ('margulies2016', 'fcgradient05', 'fsLR', '32k') | Diffusion map embedding gradient 5 of group-averaged functional connectivity |
| ('margulies2016', 'fcgradient06', 'fsLR', '32k') | Diffusion map embedding gradient 6 of group-averaged functional connectivity |
| ('margulies2016', 'fcgradient07', 'fsLR', '32k') | Diffusion map embedding gradient 7 of group-averaged functional connectivity |
| ('margulies2016', 'fcgradient08', 'fsLR', '32k') | Diffusion map embedding gradient 8 of group-averaged functional connectivity |
| ('margulies2016', 'fcgradient09', 'fsLR', '32k') | Diffusion map embedding gradient 9 of group-averaged functional connectivity |
| ('margulies2016', 'fcgradient10', 'fsLR', '32k') | Diffusion map embedding gradient 10 of group-averaged functional connectivity |
| ('mueller2013', 'intersubjvar', 'fsLR', '164k') | Intersubject variability of resting-state functional connectivity. |
| ('naganawa2020', 'lsn3172176', 'MNI152', '1mm') | PET tracer binding (BPnd) to M1 (acetylcholine receptor) |
| ('neurosynth', 'cogpc1', 'MNI152', '2mm') | PC1 of Neurosynth terms in the Cognitive Atlas (123 terms total) |
| ('norgaard2021', 'flumazenil', 'fsaverage', '164k') | PET and autoradiography informed GABAa benzodiazepine binding-site density (Bmax |

| | |
|---|---|
| ('normandin2015', 'omar', 'MNI152', '1mm') | PET tracer binding (Vt) to CB1 (cannabinoid receptor) |
| ('radnakrishnan2018', 'gsk215083', 'MNI152', '1mm') | PET tracer binding (BPnd) to 5-HT6 (serotonin receptor) |
| ('raichle', 'cbf', 'fsLR', '164k') | Cerebral blood flow |
| ('raichle', 'cbv', 'fsLR', '164k') | Cerebral blood volume |
| ('raichle', 'cmr02', 'fsLR', '164k') | Oxygen metabolism |
| ('raichle', 'cmrglc', 'fsLR', '164k') | Glucose metabolism |
| ('reardon2018', 'scalinghcp', 'civet', '41k') | Cortical areal scaling during development from the HCP dataset (S1200 release) |
| ('reardon2018', 'scalingnih', 'civet', '41k') | Cortical areal scaling during development from the NIH dataset |
| ('reardon2018', 'scalingpnc', 'civet', '41k') | Cortical areal scaling during development from the PNC dataset |
| ('rosaneto', 'abp688', 'MNI152', '1mm') | PET tracer binding (BPnd) to mGluR5 (glutamate receptor) |
| ('sandiego2015', 'flb457', 'MNI152', '1mm') | PET tracer binding (BPnd) to D2 (dopamine receptor) |
| ('sasaki2012', 'fepe2i', 'MNI152', '1mm') | PET tracer binding (BPnd) to DAT (dopamine transporter) |
| ('satterthwaite2014', 'meancbf', 'MNI152', '1mm') | Cerebral blood flow |
| ('savli2012', 'altanserin', 'MNI152', '3mm') | PET tracer binding (BPnd) to 5-HT2a (serotonin receptor) |
| ('savli2012', 'dasb', 'MNI152', '3mm') | PET tracer binding (BPnd) to 5-HTT (serotonin transporter) |
| ('savli2012', 'p943', 'MNI152', '3mm') | PET tracer binding (BPnd) to 5-HT1b (serotonin receptor) |
| ('savli2012', 'way100635', 'MNI152', '3mm') | PET tracer binding (BPnd) to 5-HT1a (serotonin receptor) |

| | |
|---|---|
| ('smart2019', 'abp688', 'MNI152', '1mm') | PET tracer binding (BPnd) to mGluR5 (glutamate receptor) |
| ('smith2017', 'flb457', 'MNI152', '1mm') | PET tracer binding (BPnd) to D2 (dopamine receptor) |
| ('sydnor2021', 'SAaxis', 'fsLR', '32k') | Sensory-association mean rank axis |
| ('tuominen', 'feobv', 'MNI152', '2mm') | PET tracer binding (SUVR) to VAChT (acetylcholine transporter) |
| ('turtonen2020', 'carfentanil', 'MNI152', '1mm') | PET tracer binding (BPnd) to MOR (mu-opioid receptor) |
| ('vijay2018', 'ly2795050', 'MNI152', '2mm') | PET tracer binding (Vt) to KOR (kappa-opioid receptor) |
| ('wey2016', 'martinostat', 'MNI152', '2mm') | PET tracer binding (SUVR) to class 1 HDAC isoforms 1, 2, and 3 (histone deacetylase) |
| ('xu2020', 'FChomology', 'fsLR', '32k') | Cross-species functional homology |
| ('xu2020', 'evoexp', 'fsLR', '32k') | Evolutionary cortical expansion |

244

**Table S4. Statistics of the latent dimensions for raw cortical thickness.**

| | Main/Mixed sample | | | | | | Women | | | | Men | | | | | |
|---|---|---|---|---|---|---|---|---|---|---|---|---|---|---|---|---|
| | 1st LD | | 2nd LD | | 3rd LD | | 1st LD | | 2nd LD | | 1st LD | | 2nd LD | | 3rd LD | |
| | Cardiomeatbolic health | | Physical robustness | | | | Cardiomeatbolic health | | Physical robustness | | Cardiomeatbolic health | | Physical robustness | | | |
| Split | r | p-value | r | p-value | r | p-value | r | p-value | r | p-value | r | p-value | r | p-value | r | p-value |
| 1 | 0.34 | **0.005** | 0.10 | **0.005** | 0.10 | **0.005** | 0.29 | **0.005** | 0.10 | **0.010** | 0.31 | **0.005** | 0.17 | **0.005** | 0.15 | **0.005** |
| 2 | 0.32 | **0.005** | 0.09 | **0.005** | 0.02 | >0.999 | 0.31 | **0.005** | 0.03 | >0.999 | 0.34 | **0.005** | 0.12 | **0.01** | 0.05 | 0.384 |
| 3 | 0.30 | **0.005** | 0.11 | **0.005** | 0.10 | **0.005** | 0.29 | **0.005** | 0.03 | >0.999 | 0.33 | **0.005** | 0.13 | **0.005** | -0.001 | >0.999 |
| 4 | 0.31 | **0.005** | 0.12 | **0.005** | 0.09 | **0.010** | 0.30 | **0.005** | 0.01 | >0.999 | 0.32 | **0.005** | 0.19 | **0.005** | 0.09 | **0.035** |
| 5 | 0.30 | **0.005** | 0.10 | **0.005** | 0.08 | **0.005** | 0.25 | **0.005** | 0.05 | 0.384 | 0.33 | **0.005** | 0.18 | **0.005** | 0.11 | **0.015** |

p-values are corrected using the Bonferroni method over 5 comparisons; significant p-values are in bold; r: Pearson's correlation coefficient

**Table S5. Statistics of the latent dimensions for raw grey matter volume.**

| | Main/Mixed sample | | | | Women | | | | Men | | | |
|---|---|---|---|---|---|---|---|---|---|---|---|---|
| | 1st LD | | 2nd LD | | 1st LD | | 2nd LD | | 1st LD | | 2nd LD | |
| | Physical robustness | | Cardiomeatbolic health | | Physical robustness | | Cardiomeatbolic health | | Physical robustness | | Cardiomeatbolic health | |
| Split | r | p-value | r | p-value | r | p-value | r | p-value | r | p-value | r | p-value |
| 1 | 0.27 | **0.005** | 0.20 | **0.005** | 0.21 | **0.005** | 0.17 | **0.005** | 0.29 | **0.005** | 0.11 | **0.015** |
| 2 | 0.27 | **0.005** | 0.16 | **0.005** | 0.23 | **0.005** | 0.20 | **0.005** | 0.27 | **0.005** | 0.15 | **0.005** |
| 3 | 0.27 | **0.005** | 0.20 | **0.005** | 0.28 | **0.005** | 0.12 | **0.010** | 0.28 | **0.005** | 0.17 | **0.005** |
| 4 | 0.27 | **0.005** | 0.17 | **0.005** | 0.24 | **0.005** | 0.15 | **0.005** | 0.26 | **0.005** | 0.12 | **0.010** |
| 5 | 0.25 | **0.005** | 0.20 | **0.005** | 0.25 | **0.005** | 0.16 | **0.005** | 0.25 | **0.005** | 0.14 | **0.005** |

p-values are corrected using the Bonferroni method over 5 comparisons; significant p-values are in bold; r: Pearson's correlation coefficient

**Table S6. Statistics of the latent dimensions for proportional cortical thickness.**

| | Main/Mixed sample | | | | | | Women | | Men | | | |
|---|---|---|---|---|---|---|---|---|---|---|---|---|
| | 1st LD | | 2nd LD | | 3rd LD | | 1st LD | | 1st LD | | 2nd LD | |
| | Cardiomeatbolic health | | Physical robustness | | | | Cardiomeatbolic health | | Cardiomeatbolic health | | Physical robustness | |
| Split | r | p-value | r | p-value | r | p-value | r | p-value | r | p-value | r | p-value |
| 1 | 0.34 | **0.005** | 0.11 | **0.005** | 0.09 | **0.005** | 0.29 | **0.005** | 0.31 | **0.005** | 0.17 | **0.005** |
| 2 | 0.33 | **0.005** | 0.06 | **0.05** | 0.02 | >0.99 | 0.32 | **0.005** | 0.34 | **0.005** | 0.11 | **0.005** |
| 3 | 0.30 | **0.005** | 0.11 | **0.005** | 0.08 | **0.005** | 0.29 | **0.005** | 0.32 | **0.005** | 0.13 | **0.005** |
| 4 | 0.32 | **0.005** | 0.12 | **0.005** | 0.11 | **0.005** | 0.29 | **0.005** | 0.31 | **0.005** | 0.17 | **0.005** |
| 5 | 0.30 | **0.005** | 0.09 | **0.005** | 0.06 | 0.070 | 0.26 | **0.005** | 0.32 | **0.005** | 0.16 | **0.005** |

p-values are corrected using the Bonferroni method over 5 comparisons; significant p-values are in bold; r: Pearson's correlation coefficient

**Table S7. Statistics of the latent dimensions for proportional grey matter volume.**

| | Main/Mixed sample | | | | | | Women | | | | Men | | | | | |
|---|---|---|---|---|---|---|---|---|---|---|---|---|---|---|---|---|
| | $1^{st}$ LD | | $2^{nd}$ LD | | $3^{rd}$ LD | | $1^{st}$ LD | | $2^{nd}$ LD | | $1^{st}$ LD | | $2^{nd}$ LD | | $3^{rd}$ LD | |
| | Cardiomeatbolic health | | Physical robustness | | | | Cardiomeatbolic health | | Physical robustness | | Cardiomeatbolic health | | Physical robustness | | | |
| Split | r | p-value | r | p-value | r | p-value | r | p-value | r | p-value | r | p-value | r | p-value | r | p-value |
| 1 | 0.18 | **0.005** | 0.17 | **0.005** | 0.07 | **0.035** | 0.17 | **0.005** | 0.12 | **0.010** | 0.12 | **0.010** | 0.21 | **0.005** | 0.07 | 0.13 |
| 2 | 0.16 | **0.005** | 0.16 | **0.005** | 0.10 | **0.005** | 0.21 | **0.005** | 0.12 | **0.005** | 0.13 | **0.010** | 0.16 | **0.005** | 0.08 | 0.12 |
| 3 | 0.22 | **0.005** | 0.19 | **0.005** | 0.08 | **0.020** | 0.16 | **0.005** | 0.13 | **0.005** | 0.18 | **0.005** | 0.16 | **0.005** | 0.09 | 0.055 |
| 4 | 0.19 | **0.005** | 0.19 | **0.005** | 0.14 | **0.005** | 0.15 | **0.005** | 0.09 | 0.055 | 0.12 | **0.005** | 0.20 | **0.005** | 0.09 | **0.045** |
| 5 | 0.15 | **0.005** | 0.15 | **0.005** | 0.08 | **0.015** | 0.17 | **0.005** | 0.12 | **0.005** | 0.15 | **0.005** | 0.15 | **0.005** | 0.12 | **0.010** |

p-values are corrected using the Bonferroni method over 5 comparisons; significant p-values are in bold; r: Pearson's correlation coefficient

**Table S8. Statistics of the latent dimensions for corrected cortical thickness.**

| | Main/Mixed sample | | | | | | Women | | | | Men | | | |
|---|---|---|---|---|---|---|---|---|---|---|---|---|---|---|
| | 1st LD | | 2nd LD | | 3rd LD | | 1st LD | | 2nd LD | | 1st LD | | 2nd LD | |
| | Cardiomeatbolic health | | Physical robustness | | | | Cardiomeatbolic health | | Physical robustness | | Cardiomeatbolic health | | Physical robustness | |
| Split | r | p-value | r | p-value | r | p-value | r | p-value | r | p-value | r | p-value | r | p-value |
| 1 | 0.34 | **0.005** | 0.11 | **0.005** | 0.09 | **0.005** | 0.29 | **0.005** | 0.01 | >0.99 | 0.31 | **0.005** | 0.17 | **0.005** |
| 2 | 0.33 | **0.005** | 0.06 | **0.024** | 0.01 | >0.99 | 0.32 | **0.005** | 0.003 | >0.99 | 0.34 | **0.005** | 0.11 | **0.005** |
| 3 | 0.30 | **0.005** | 0.11 | **0.005** | 0.08 | **0.005** | 0.30 | **0.005** | 0.05 | 0.479 | 0.32 | **0.005** | 0.20 | **0.010** |
| 4 | 0.32 | **0.005** | 0.12 | **0.005** | 0.11 | **0.005** | 0.30 | **0.005** | 0.12 | **0.010** | 0.31 | **0.005** | 0.18 | **0.005** |
| 5 | 0.30 | **0.005** | 0.09 | **0.005** | 0.08 | **0.005** | 0.26 | **0.005** | 0.03 | 0.374 | 0.32 | **0.005** | 0.17 | **0.015** |

p-values are corrected using the Bonferroni method over 5 comparisons; significant p-values are in bold; r: Pearson’s correlation coefficient

**Table S9. Statistics of the latent dimensions for corrected grey matter volume.**

| | Main/Mixed sample | | | | | | Women | | | | | | Men | | | |
|---|---|---|---|---|---|---|---|---|---|---|---|---|---|---|---|---|
| | 1st LD | | 2nd LD | | 3rd LD | | 1st LD | | 2nd LD | | 3rd LD | | 1st LD | | 2nd LD | |
| | Cardiomeatbolic health | | Physical robustness | | | | Cardiomeatbolic health | | Physical robustness | | | | Cardiomeatbolic health | | Physical robustness | |
| Split | r | p-value | r | p-value | r | p-value | r | p-value | r | p-value | r | p-value | r | p-value | r | p-value |
| 1 | 0.22 | **0.005** | 0.14 | **0.005** | 0.07 | **0.020** | 0.17 | **0.005** | 0.08 | 0.080 | 0.09 | **0.030** | 0.12 | **0.010** | 0.16 | **0.005** |
| 2 | 0.19 | **0.005** | 0.13 | **0.005** | 0.04 | 0.329 | 0.21 | **0.005** | 0.11 | **0.005** | 0.05 | 0.414 | 0.16 | **0.005** | 0.13 | **0.005** |
| 3 | 0.23 | **0.005** | 0.09 | **0.005** | 0.06 | **0.035** | 0.14 | **0.005** | 0.13 | **0.010** | 0.12 | **0.010** | 0.18 | **0.005** | 0.17 | **0.005** |
| 4 | 0.19 | **0.005** | 0.12 | **0.005** | 0.10 | **0.005** | 0.15 | **0.005** | 0.09 | **0.030** | 0.02 | <0.99 | 0.12 | **0.010** | 0.17 | **0.005** |
| 5 | 0.17 | **0.005** | 0.11 | **0.005** | 0.06 | **0.025** | 0.16 | **0.005** | 0.10 | **0.030** | 0.06 | 0.264 | 0.15 | **0.005** | 0.13 | **0.005** |

p-values are corrected using the Bonferroni method over 5 comparisons; significant p-values are in bold; r: Pearson's correlation coefficient

**Table S10. Association between the brain loadings of the latent dimension of cardiometabolic health and neuromaps.**

| | | | CT Main/MIxed | | | | | | GMV Main/MIxed | | | | | |
|---|---|---|---|---|---|---|---|---|---|---|---|---|---|---|
| | | | Raw | | Proportional | | Corrected | | Raw | | Proportional | | Corrected | |
| id | Category | Subcategory | r | p-value | r | p-value | r | p-value | r | p-value | r | p-value | r | p-value |
| abagen_genepc1_fsaverage_10k | Genetics | PC1 Allen | -0.647 | 0.026 | -0.599 | 0.103 | -0.625 | 0.026 | -0.024 | > 0.999 | -0.147 | > 0.999 | -0.164 | > 0.999 |
| aghourian2017_feobv_MNI152_1mm | Molecular | VAChT | 0.524 | > 0.999 | 0.525 | > 0.999 | 0.525 | > 0.999 | 0.012 | > 0.999 | -0.033 | > 0.999 | 0.115 | > 0.999 |
| alarkurtti2015_raclopride_MNI152_3mm | Molecular | D2 | -0.179 | > 0.999 | -0.197 | > 0.999 | -0.198 | > 0.999 | 0.309 | > 0.999 | 0.290 | > 0.999 | 0.289 | > 0.999 |
| bedard2019_feobv_MNI152_1mm | Molecular | VAChT | 0.729 | 0.026 | 0.711 | 0.026 | 0.729 | 0.026 | 0.267 | > 0.999 | 0.148 | > 0.999 | 0.277 | > 0.999 |
| beliveau2017_az10419369_fsaverage_164k | Molecular | 5-HT1b | -0.413 | > 0.999 | -0.425 | > 0.999 | -0.424 | > 0.999 | -0.411 | 0.052 | -0.481 | 0.026 | -0.306 | > 0.999 |
| beliveau2017_cimbi36_fsaverage_164k | Molecular | 5-HT2a | 0.083 | > 0.999 | 0.053 | > 0.999 | 0.069 | > 0.999 | 0.010 | > 0.999 | 0.312 | > 0.999 | 0.226 | > 0.999 |
| beliveau2017_cumi101_fsaverage_164k | Molecular | 5-HT1a | 0.732 | 0.026 | 0.685 | 0.026 | 0.709 | 0.026 | 0.275 | > 0.999 | 0.454 | > 0.999 | 0.380 | > 0.999 |
| beliveau2017_dasb_fsaverage_164k | Molecular | 5-HTT | 0.516 | > 0.999 | 0.538 | 0.284 | 0.541 | > 0.999 | -0.052 | > 0.999 | -0.359 | > 0.999 | -0.224 | > 0.999 |

| | | | | | | | | | | | | | | |
|---|---|---|---|---|---|---|---|---|---|---|---|---|---|---|
| beliveau2017_sb207145_fsaverage_164k | Molecular | 5-HT4 | 0.420 | > 0.999 | 0.381 | > 0.999 | 0.405 | > 0.999 | 0.314 | > 0.999 | 0.598 | 0.542 | 0.477 | > 0.999 |
| castrillon2023_cmrglc_MNI152_3mm | Metabolism | Glucose | -0.281 | > 0.999 | -0.324 | > 0.999 | -0.316 | > 0.999 | -0.241 | > 0.999 | -0.062 | > 0.999 | -0.013 | > 0.999 |
| ding2010_mrb_MNI152_1mm | Molecular | NET | -0.205 | > 0.999 | -0.199 | > 0.999 | -0.223 | > 0.999 | -0.041 | > 0.999 | 0.033 | > 0.999 | 0.090 | > 0.999 |
| dubois2015_abp688_MNI152_1mm | Molecular | mGluR5 | 0.365 | > 0.999 | 0.330 | > 0.999 | 0.350 | > 0.999 | -0.094 | > 0.999 | 0.117 | > 0.999 | 0.170 | > 0.999 |
| dukart2018_flumazenil_MNI152_3mm | Molecular | GABAa | -0.049 | > 0.999 | -0.041 | > 0.999 | -0.048 | > 0.999 | -0.212 | > 0.999 | -0.132 | > 0.999 | -0.033 | > 0.999 |
| dukart2018_fpcit_MNI152_3mm | Molecular | DAT | 0.575 | 0.026 | 0.553 | 0.026 | 0.571 | 0.026 | 0.238 | > 0.999 | 0.034 | > 0.999 | 0.043 | > 0.999 |
| fazio2016_madam_MNI152_3mm | Molecular | 5-HTT | 0.282 | > 0.999 | 0.262 | > 0.999 | 0.275 | > 0.999 | -0.169 | > 0.999 | -0.401 | 0.671 | -0.349 | > 0.999 |
| finnema2016_ucbj_MNI152_1mm | Molecular | SV2a | -0.054 | > 0.999 | -0.061 | > 0.999 | -0.067 | > 0.999 | -0.209 | > 0.999 | 0.043 | > 0.999 | 0.011 | > 0.999 |
| gallezot2010_p943_MNI152_1mm | Molecular | 5-HT1b | -0.428 | > 0.999 | -0.427 | > 0.999 | -0.427 | > 0.999 | -0.424 | 0.026 | -0.343 | > 0.999 | -0.239 | > 0.999 |
| gallezot2017_gsk189254_MNI152_1mm | Molecular | H3 | 0.194 | > 0.999 | 0.164 | > 0.999 | 0.170 | > 0.999 | -0.405 | 0.026 | -0.220 | > 0.999 | -0.167 | > 0.999 |
| galovic2021_ge179_MNI152_1mm | Molecular | NMDA | 0.360 | > 0.999 | 0.327 | > 0.999 | 0.342 | > 0.999 | 0.044 | > 0.999 | 0.257 | > 0.999 | 0.167 | > 0.999 |
| hcps1200_megalpha_fsLR_4k | Brain function | MEG 1alpha | -0.434 | > 0.999 | -0.343 | > 0.999 | -0.373 | > 0.999 | 0.195 | > 0.999 | 0.088 | > 0.999 | 0.017 | > 0.999 |

| | | | | | | | | | | | | | | |
|---|---|---|---|---|---|---|---|---|---|---|---|---|---|---|
| hcps1200_megbeta_fsLR_4k | Brain function | MEG 2beta | -0.242 | > 0.999 | -0.264 | > 0.999 | -0.273 | > 0.999 | 0.042 | > 0.999 | 0.073 | > 0.999 | 0.106 | > 0.999 |
| hcps1200_megdelta_fsLR_4k | Brain function | MEG 3delta | 0.498 | > 0.999 | 0.437 | > 0.999 | 0.476 | > 0.999 | -0.203 | > 0.999 | -0.152 | > 0.999 | -0.061 | > 0.999 |
| hcps1200_meggamma1_fsLR_4k | Brain function | MEG 4low gamma | 0.264 | > 0.999 | 0.198 | > 0.999 | 0.221 | > 0.999 | -0.284 | > 0.999 | -0.206 | > 0.999 | -0.110 | > 0.999 |
| hcps1200_meggamma2_fsLR_4k | Brain function | MEG 5high gamma | 0.462 | > 0.999 | 0.438 | > 0.999 | 0.467 | > 0.999 | -0.258 | > 0.999 | -0.076 | > 0.999 | -0.032 | > 0.999 |
| hcps1200_megtheta_fsLR_4k | Brain function | MEG 6theta | 0.292 | > 0.999 | 0.222 | > 0.999 | 0.238 | > 0.999 | -0.182 | > 0.999 | -0.096 | > 0.999 | -0.014 | > 0.999 |
| hcps1200_megtimescale_fsLR_4k | Brain function | MEG intts | 0.488 | > 0.999 | 0.428 | > 0.999 | 0.474 | > 0.999 | -0.226 | > 0.999 | -0.119 | > 0.999 | -0.082 | > 0.999 |
| hcps1200_myelinmap_fsLR_32k | Brain structure | T1w/T2w | -0.426 | > 0.999 | -0.361 | > 0.999 | -0.388 | > 0.999 | 0.054 | > 0.999 | -0.238 | > 0.999 | -0.180 | > 0.999 |
| hcps1200_thickness_fsLR_32k | Brain structure | CT | 0.731 | 0.026 | 0.672 | 0.026 | 0.706 | 0.026 | 0.062 | > 0.999 | 0.325 | > 0.999 | 0.264 | > 0.999 |
| hesse2017_methylreboxetine_MNI152_3mm | Molecular | NET | 0.286 | > 0.999 | 0.293 | > 0.999 | 0.273 | > 0.999 | 0.065 | > 0.999 | 0.100 | > 0.999 | 0.175 | > 0.999 |
| hillmer2016_flubatine_MNI152_1mm | Molecular | a4b2 | -0.174 | > 0.999 | -0.205 | > 0.999 | -0.208 | > 0.999 | -0.112 | > 0.999 | 0.044 | > 0.999 | 0.090 | > 0.999 |
| jaworska2020_fallypride_MNI152_1mm | Molecular | D2 | 0.582 | 0.155 | 0.516 | > 0.999 | 0.542 | > 0.999 | 0.275 | > 0.999 | 0.551 | 0.516 | 0.454 | > 0.999 |
| kaller2017_sch23390_MNI152_3mm | Molecular | D1 | 0.366 | > 0.999 | 0.331 | > 0.999 | 0.350 | > 0.999 | 0.056 | > 0.999 | 0.100 | > 0.999 | 0.084 | > 0.999 |

| | | | | | | | | | | | | | | |
|---|---|---|---|---|---|---|---|---|---|---|---|---|---|---|
| kantonen2020_carfentanil_MNI152_3mm | Molecular | MOR | 0.461 | > 0.999 | 0.394 | > 0.999 | 0.421 | > 0.999 | -0.059 | > 0.999 | 0.154 | > 0.999 | 0.210 | > 0.999 |
| kim2020_ps13_MNI152_2mm | Molecular | COX-1 | -0.312 | > 0.999 | -0.245 | > 0.999 | -0.256 | > 0.999 | 0.156 | > 0.999 | -0.190 | > 0.999 | -0.153 | > 0.999 |
| laurikainen2018_fmpepd2_MNI152_1mm | Molecular | CB1 | -0.030 | > 0.999 | -0.077 | > 0.999 | -0.069 | > 0.999 | -0.125 | > 0.999 | 0.195 | > 0.999 | 0.211 | > 0.999 |
| lois2018_pbr28_MNI152_2mm | Molecular | TSPO | 0.543 | 0.052 | 0.519 | 0.026 | 0.536 | 0.052 | 0.198 | > 0.999 | 0.032 | > 0.999 | 0.066 | > 0.999 |
| lukow2022_ro154513_MNI152_2mm | Molecular | GABAa | 0.636 | 0.026 | 0.587 | 0.026 | 0.612 | 0.026 | -0.032 | > 0.999 | 0.195 | > 0.999 | 0.182 | > 0.999 |
| malen2022_raclopride_MNI152_2mm | Molecular | D2 | 0.244 | > 0.999 | 0.214 | > 0.999 | 0.230 | > 0.999 | 0.401 | 0.026 | 0.364 | 0.181 | 0.285 | 0.103 |
| margulies2016_fcgradient01_fsLR_32k | Brain function | FC Grad01 | 0.219 | > 0.999 | 0.149 | > 0.999 | 0.181 | > 0.999 | -0.047 | > 0.999 | 0.271 | > 0.999 | 0.185 | > 0.999 |
| margulies2016_fcgradient02_fsLR_32k | Brain function | FC Grad02 | -0.390 | > 0.999 | -0.390 | > 0.999 | -0.382 | > 0.999 | 0.023 | > 0.999 | -0.023 | > 0.999 | -0.192 | > 0.999 |
| margulies2016_fcgradient03_fsLR_32k | Brain function | FC Grad03 | -0.163 | > 0.999 | -0.197 | > 0.999 | -0.188 | > 0.999 | -0.125 | > 0.999 | -0.116 | > 0.999 | -0.153 | > 0.999 |
| margulies2016_fcgradient04_fsLR_32k | Brain function | FC Grad04 | 0.162 | > 0.999 | 0.219 | > 0.999 | 0.213 | > 0.999 | -0.277 | > 0.999 | -0.548 | 0.026 | -0.401 | > 0.999 |
| margulies2016_fcgradient05_fsLR_32k | Brain function | FC Grad05 | 0.571 | 0.026 | 0.542 | 0.052 | 0.566 | 0.026 | 0.192 | > 0.999 | 0.379 | > 0.999 | 0.225 | > 0.999 |
| margulies2016_fcgradient06_fsLR_32k | Brain function | FC Grad06 | -0.315 | > 0.999 | -0.303 | > 0.999 | -0.306 | > 0.999 | -0.234 | > 0.999 | -0.198 | > 0.999 | -0.251 | > 0.999 |

| | | | | | | | | | | | | | | |
|---|---|---|---|---|---|---|---|---|---|---|---|---|---|---|
| margulies2016_fcgradient07_fsLR_32k | Brain function | FC Grad07 | 0.070 | > 0.999 | 0.051 | > 0.999 | 0.056 | > 0.999 | -0.062 | > 0.999 | -0.376 | > 0.999 | -0.320 | > 0.999 |
| margulies2016_fcgradient08_fsLR_32k | Brain function | FC Grad08 | 0.096 | > 0.999 | 0.080 | > 0.999 | 0.086 | > 0.999 | 0.379 | 0.026 | 0.252 | > 0.999 | 0.093 | > 0.999 |
| margulies2016_fcgradient09_fsLR_32k | Brain function | FC Grad09 | -0.033 | > 0.999 | -0.042 | > 0.999 | -0.041 | > 0.999 | -0.330 | > 0.999 | -0.266 | > 0.999 | -0.130 | > 0.999 |
| margulies2016_fcgradient10_fsLR_32k | Brain function | FC Grad10 | -0.228 | > 0.999 | -0.204 | > 0.999 | -0.210 | > 0.999 | -0.326 | 0.026 | -0.238 | > 0.999 | -0.182 | > 0.999 |
| mueller2013_intersubjvar_fsLR_164k | Brain function | RSFC intvar | 0.158 | > 0.999 | 0.088 | > 0.999 | 0.119 | > 0.999 | 0.144 | > 0.999 | 0.450 | > 0.999 | 0.347 | > 0.999 |
| naganawa2020_lsn3172176_MNI152_1mm | Molecular | M1 | -0.097 | > 0.999 | -0.115 | > 0.999 | -0.118 | > 0.999 | -0.238 | > 0.999 | 0.035 | > 0.999 | 0.027 | > 0.999 |
| neurosynth_cogpc1_MNI152_2mm | Cognition | PC1 Cognition | -0.250 | > 0.999 | -0.209 | > 0.999 | -0.236 | > 0.999 | 0.096 | > 0.999 | 0.071 | > 0.999 | 0.078 | > 0.999 |
| norgaard2021_flumazenil_fsaverage_164k | Molecular | GABAa | -0.113 | > 0.999 | -0.101 | > 0.999 | -0.098 | > 0.999 | -0.322 | 0.929 | -0.294 | > 0.999 | -0.134 | > 0.999 |
| normandin2015_omar_MNI152_1mm | Molecular | CB1 | 0.367 | > 0.999 | 0.326 | > 0.999 | 0.341 | > 0.999 | -0.106 | > 0.999 | 0.172 | > 0.999 | 0.217 | > 0.999 |
| radnakrishnan2018_gsk215083_MNI152_1mm | Molecular | 5-HT6 | 0.057 | > 0.999 | 0.012 | > 0.999 | 0.028 | > 0.999 | -0.045 | > 0.999 | 0.221 | > 0.999 | 0.157 | > 0.999 |
| raichle_cbf_fsLR_164k | Physiology | Blood flow | -0.235 | > 0.999 | -0.236 | > 0.999 | -0.250 | > 0.999 | -0.257 | > 0.999 | -0.514 | > 0.999 | -0.327 | > 0.999 |
| raichle_cbv_fsLR_164k | Physiology | Blood vol | 0.176 | > 0.999 | 0.210 | > 0.999 | 0.202 | > 0.999 | -0.171 | > 0.999 | -0.333 | > 0.999 | -0.329 | > 0.999 |

| | | | | | | | | | | | | | | |
|---|---|---|---|---|---|---|---|---|---|---|---|---|---|---|
| raichle_cmr02_fsLR_164k | Metabolism | Oxygen | -0.599 | 0.129 | -0.581 | 0.361 | -0.608 | 0.284 | -0.045 | > 0.999 | -0.260 | > 0.999 | -0.213 | > 0.999 |
| raichle_cmrglc_fsLR_164k | Metabolism | Glucose | -0.628 | 0.026 | -0.629 | 0.026 | -0.643 | 0.026 | -0.175 | > 0.999 | -0.279 | > 0.999 | -0.236 | > 0.999 |
| reardon2018_scalinghcp_civet_41k | Brain structure | Expansion | -0.167 | > 0.999 | -0.202 | > 0.999 | -0.188 | > 0.999 | -0.040 | > 0.999 | 0.065 | > 0.999 | -0.001 | > 0.999 |
| reardon2018_scalingnih_civet_41k | Brain structure | Expansion | 0.065 | > 0.999 | 0.029 | > 0.999 | 0.031 | > 0.999 | 0.237 | > 0.999 | 0.585 | 0.026 | 0.399 | 0.181 |
| reardon2018_scalingpnc_civet_41k | Brain structure | Expansion | -0.076 | > 0.999 | -0.127 | > 0.999 | -0.115 | > 0.999 | 0.111 | > 0.999 | 0.350 | > 0.999 | 0.195 | > 0.999 |
| rosaneto_abp688_MNI152_1mm | Molecular | mGluR5 | 0.230 | > 0.999 | 0.202 | > 0.999 | 0.216 | > 0.999 | -0.110 | > 0.999 | 0.060 | > 0.999 | 0.124 | > 0.999 |
| sandiego2015_flb457_MNI152_1mm | Molecular | D2 | 0.604 | 0.155 | 0.543 | 0.206 | 0.564 | 0.335 | 0.222 | > 0.999 | 0.491 | > 0.999 | 0.402 | > 0.999 |
| sasaki2012_fepe2i_MNI152_1mm | Molecular | DAT | 0.090 | > 0.999 | 0.082 | > 0.999 | 0.089 | > 0.999 | 0.297 | 0.026 | 0.158 | > 0.999 | 0.145 | > 0.999 |
| satterthwaite2014_meancbf_MNI152_1mm | Physiology | Blood flow | 0.048 | > 0.999 | 0.018 | > 0.999 | 0.030 | > 0.999 | -0.033 | > 0.999 | 0.080 | > 0.999 | 0.102 | > 0.999 |
| savli2012_altanserin_MNI152_3mm | Molecular | 5-HT2a | 0.047 | > 0.999 | 0.009 | > 0.999 | 0.026 | > 0.999 | 0.064 | > 0.999 | 0.401 | > 0.999 | 0.349 | > 0.999 |
| savli2012_dasb_MNI152_3mm | Molecular | 5-HTT | 0.533 | 0.980 | 0.533 | 0.077 | 0.543 | 0.851 | -0.112 | > 0.999 | -0.300 | > 0.999 | -0.160 | > 0.999 |
| savli2012_p943_MNI152_3mm | Molecular | 5-HT1b | -0.286 | > 0.999 | -0.303 | > 0.999 | -0.299 | > 0.999 | -0.208 | > 0.999 | -0.116 | > 0.999 | 0.015 | > 0.999 |

| | | | | | | | | | | | | | | |
|---|---|---|---|---|---|---|---|---|---|---|---|---|---|---|
| savli2012_way100635_MNI152_3mm | Molecular | 5-HT1a | 0.630 | 0.026 | 0.583 | 0.026 | 0.599 | 0.026 | 0.260 | > 0.999 | 0.494 | > 0.999 | 0.416 | > 0.999 |
| smart2019_abp688_MNI152_1mm | Molecular | mGluR5 | 0.292 | > 0.999 | 0.253 | > 0.999 | 0.264 | > 0.999 | -0.051 | > 0.999 | 0.163 | > 0.999 | 0.202 | > 0.999 |
| smith2017_flb457_MNI152_1mm | Molecular | D2 | 0.530 | 0.645 | 0.476 | > 0.999 | 0.501 | > 0.999 | 0.265 | > 0.999 | 0.505 | > 0.999 | 0.446 | > 0.999 |
| sydnor2021_SAaxis_fsLR_32k | Brain function | Sensory-association | 0.373 | > 0.999 | 0.277 | > 0.999 | 0.324 | > 0.999 | -0.021 | > 0.999 | 0.305 | > 0.999 | 0.233 | > 0.999 |
| tuominen_feobv_MNI152_2mm | Molecular | VAChT | 0.548 | 0.026 | 0.562 | 0.026 | 0.552 | 0.026 | 0.114 | > 0.999 | -0.015 | > 0.999 | 0.148 | > 0.999 |
| turtonen2020_carfentanil_MNI152_1mm | Molecular | MOR | 0.429 | > 0.999 | 0.355 | > 0.999 | 0.385 | > 0.999 | -0.055 | > 0.999 | 0.170 | > 0.999 | 0.238 | > 0.999 |
| vijay2018_ly2795050_MNI152_2mm | Molecular | KOR | 0.280 | > 0.999 | 0.243 | > 0.999 | 0.243 | > 0.999 | -0.179 | > 0.999 | 0.087 | > 0.999 | 0.146 | > 0.999 |
| wey2016_martinostat_MNI152_2mm | Molecular | HDAC | -0.385 | 0.026 | -0.396 | 0.026 | -0.402 | 0.026 | -0.408 | 0.026 | -0.293 | 0.387 | -0.209 | > 0.999 |
| xu2020_FChomology_fsLR_32k | Brain function | Functional homology | -0.238 | > 0.999 | -0.188 | > 0.999 | -0.212 | > 0.999 | -0.099 | > 0.999 | -0.326 | > 0.999 | -0.232 | > 0.999 |
| xu2020_evoexp_fsLR_32k | Brain structure | Expansion | 0.105 | > 0.999 | 0.058 | > 0.999 | 0.076 | > 0.999 | 0.043 | > 0.999 | 0.348 | > 0.999 | 0.235 | > 0.999 |

263

**Table S11. Association between the brain loadings of the latent dimension of physical robustness and neuromaps.**

| | | | CT Men | | | | | | GMV Main/Mixed | | | | | |
|---|---|---|---|---|---|---|---|---|---|---|---|---|---|---|
| | | | Raw | | Proportional | | Corrected | | Raw | | Proportional | | Corrected | |
| id | Category | Subcategory | r | p-value | r | p-value | r | p-value | r | p-value | r | p-value | r | p-value |
| abagen_genepc1_fsaverage_10k | Genetics | PC1 Allen | -0.339 | > 0.999 | -0.308 | > 0.999 | -0.478 | > 0.999 | -0.370 | > 0.999 | -0.272 | > 0.999 | -0.316 | > 0.999 |
| aghourian2017_feobv_MNI152_1mm | Molecular | VAChT | 0.267 | > 0.999 | 0.507 | > 0.999 | 0.368 | > 0.999 | 0.292 | > 0.999 | 0.074 | > 0.999 | 0.493 | 0.026 |
| alarkurtti2015_raclopride_MNI152_3mm | Molecular | D2 | 0.249 | > 0.999 | 0.074 | > 0.999 | -0.018 | > 0.999 | 0.070 | > 0.999 | 0.194 | > 0.999 | 0.086 | > 0.999 |
| bedard2019_feobv_MNI152_1mm | Molecular | VAChT | 0.181 | > 0.999 | 0.368 | > 0.999 | 0.422 | > 0.999 | 0.285 | > 0.999 | 0.124 | > 0.999 | 0.299 | > 0.999 |
| beliveau2017_az10419369_fsaverage_164k | Molecular | 5-HT1b | 0.312 | > 0.999 | 0.301 | > 0.999 | 0.278 | > 0.999 | -0.050 | > 0.999 | -0.081 | > 0.999 | 0.185 | > 0.999 |
| beliveau2017_cimbi36_fsaverage_164k | Molecular | 5-HT2a | 0.356 | 0.490 | 0.080 | > 0.999 | 0.155 | > 0.999 | 0.399 | 0.052 | 0.403 | 0.258 | 0.316 | > 0.999 |
| beliveau2017_cumi101_fsaverage_164k | Molecular | 5-HT1a | 0.257 | > 0.999 | 0.193 | > 0.999 | 0.375 | > 0.999 | 0.434 | 0.026 | 0.314 | > 0.999 | 0.297 | 0.851 |
| beliveau2017_dasb_fsaverage_164k | Molecular | 5-HTT | -0.138 | > 0.999 | 0.118 | > 0.999 | 0.268 | > 0.999 | -0.151 | > 0.999 | -0.162 | > 0.999 | -0.024 | > 0.999 |

| | | | | | | | | | | | | | | |
|---|---|---|---|---|---|---|---|---|---|---|---|---|---|---|
| beliveau2017_sb207145_fsaverage_164k | Molecular | 5-HT4 | 0.167 | > 0.999 | -0.128 | > 0.999 | -0.001 | > 0.999 | 0.521 | 0.077 | 0.507 | 0.181 | 0.348 | 0.464 |
| castrillon2023_cmrglc_MNI152_3mm | Metabolism | Glucose | 0.276 | 0.697 | 0.119 | > 0.999 | 0.067 | > 0.999 | 0.242 | > 0.999 | 0.324 | 0.077 | 0.354 | 0.026 |
| ding2010_mrb_MNI152_1mm | Molecular | NET | 0.068 | > 0.999 | 0.159 | > 0.999 | -0.131 | > 0.999 | 0.116 | > 0.999 | -0.010 | > 0.999 | 0.236 | > 0.999 |
| dubois2015_abp688_MNI152_1mm | Molecular | mGluR5 | 0.360 | 0.284 | 0.244 | > 0.999 | 0.276 | > 0.999 | 0.547 | 0.026 | 0.476 | 0.026 | 0.536 | 0.026 |
| dukart2018_flumazenil_MNI152_3mm | Molecular | GABAa | -0.027 | > 0.999 | -0.124 | > 0.999 | 0.015 | > 0.999 | 0.145 | > 0.999 | 0.204 | > 0.999 | 0.059 | > 0.999 |
| dukart2018_fpcit_MNI152_3mm | Molecular | DAT | 0.077 | > 0.999 | 0.153 | > 0.999 | 0.398 | > 0.999 | -0.047 | > 0.999 | 0.157 | > 0.999 | -0.061 | > 0.999 |
| fazio2016_madam_MNI152_3mm | Molecular | 5-HTT | 0.165 | > 0.999 | 0.251 | > 0.999 | 0.409 | > 0.999 | -0.042 | > 0.999 | -0.050 | > 0.999 | 0.120 | > 0.999 |
| finnema2016_ucbj_MNI152_1mm | Molecular | SV2a | -0.054 | > 0.999 | -0.149 | > 0.999 | -0.174 | > 0.999 | 0.251 | 0.439 | 0.146 | > 0.999 | 0.250 | 0.052 |
| gallezot2010_p943_MNI152_1mm | Molecular | 5-HT1b | 0.263 | > 0.999 | 0.267 | > 0.999 | 0.208 | > 0.999 | 0.040 | > 0.999 | 0.001 | > 0.999 | 0.191 | > 0.999 |
| gallezot2017_gsk189254_MNI152_1mm | Molecular | H3 | 0.297 | > 0.999 | 0.437 | > 0.999 | 0.340 | > 0.999 | 0.357 | 0.026 | 0.152 | > 0.999 | 0.502 | 0.026 |
| galovic2021_ge179_MNI152_1mm | Molecular | NMDA | 0.256 | > 0.999 | 0.193 | > 0.999 | 0.282 | > 0.999 | 0.403 | 0.026 | 0.297 | 0.697 | 0.432 | 0.026 |
| hcps1200_megalpha_fsLR_4k | Brain function | MEG 1alpha | -0.469 | > 0.999 | -0.641 | > 0.999 | -0.614 | > 0.999 | -0.245 | > 0.999 | -0.005 | > 0.999 | -0.372 | 0.026 |

| | | | | | | | | | | | | | | |
|---|---|---|---|---|---|---|---|---|---|---|---|---|---|---|
| hcps1200_megbeta_fsLR_4k | Brain function | MEG 2beta | 0.200 | > 0.999 | 0.186 | > 0.999 | -0.119 | > 0.999 | 0.136 | > 0.999 | 0.083 | > 0.999 | 0.261 | > 0.999 |
| hcps1200_megdelta_fsLR_4k | Brain function | MEG 3delta | 0.391 | > 0.999 | 0.549 | > 0.999 | 0.704 | 0.026 | 0.194 | > 0.999 | 0.040 | > 0.999 | 0.250 | > 0.999 |
| hcps1200_meggamma1_fsLR_4k | Brain function | MEG 4low gamma | 0.343 | > 0.999 | 0.562 | > 0.999 | 0.476 | > 0.999 | 0.168 | > 0.999 | -0.056 | > 0.999 | 0.318 | 0.129 |
| hcps1200_meggamma2_fsLR_4k | Brain function | MEG 5high gamma | 0.250 | > 0.999 | 0.405 | > 0.999 | 0.596 | > 0.999 | 0.167 | > 0.999 | 0.033 | > 0.999 | 0.181 | > 0.999 |
| hcps1200_megtheta_fsLR_4k | Brain function | MEG 6theta | 0.460 | > 0.999 | 0.622 | > 0.999 | 0.493 | > 0.999 | 0.239 | > 0.999 | 0.105 | > 0.999 | 0.401 | 0.026 |
| hcps1200_megtimescale_fsLR_4k | Brain function | MEG intts | 0.315 | > 0.999 | 0.448 | > 0.999 | 0.639 | 0.258 | 0.189 | > 0.999 | 0.015 | > 0.999 | 0.189 | > 0.999 |
| hcps1200_myelinmap_fsLR_32k | Brain structure | T1w/T2w | -0.379 | > 0.999 | -0.280 | > 0.999 | -0.369 | > 0.999 | -0.485 | 0.026 | -0.296 | > 0.999 | -0.450 | 0.026 |
| hcps1200_thickness_fsLR_32k | Brain structure | CT | 0.232 | > 0.999 | 0.258 | > 0.999 | 0.381 | > 0.999 | 0.480 | 0.026 | 0.253 | > 0.999 | 0.363 | > 0.999 |
| hesse2017_methylreboxetine_MNI152_3mm | Molecular | NET | -0.084 | > 0.999 | 0.078 | > 0.999 | -0.024 | > 0.999 | 0.188 | > 0.999 | -0.006 | > 0.999 | 0.209 | > 0.999 |
| hillmer2016_flubatine_MNI152_1mm | Molecular | a4b2 | 0.480 | 0.748 | 0.401 | > 0.999 | 0.249 | > 0.999 | 0.279 | > 0.999 | 0.154 | > 0.999 | 0.411 | 0.026 |
| jaworska2020_fallypride_MNI152_1mm | Molecular | D2 | 0.061 | > 0.999 | -0.070 | > 0.999 | 0.082 | > 0.999 | 0.424 | 0.542 | 0.334 | > 0.999 | 0.300 | > 0.999 |
| kaller2017_sch23390_MNI152_3mm | Molecular | D1 | 0.231 | > 0.999 | 0.230 | > 0.999 | 0.407 | > 0.999 | 0.247 | > 0.999 | 0.292 | > 0.999 | 0.248 | > 0.999 |

| | | | | | | | | | | | | | | |
|---|---|---|---|---|---|---|---|---|---|---|---|---|---|---|
| kantonen2020_carfentanil_MNI152_3mm | Molecular | MOR | 0.502 | 0.413 | 0.440 | > 0.999 | 0.474 | > 0.999 | 0.547 | 0.026 | 0.350 | > 0.999 | 0.582 | 0.026 |
| kim2020_ps13_MNI152_2mm | Molecular | COX-1 | -0.144 | > 0.999 | -0.191 | > 0.999 | -0.126 | > 0.999 | -0.384 | 0.258 | -0.156 | > 0.999 | -0.340 | 0.052 |
| laurikainen2018_fmpepd2_MNI152_1mm | Molecular | CB1 | 0.372 | 0.980 | 0.175 | > 0.999 | 0.043 | > 0.999 | 0.522 | 0.026 | 0.471 | 0.052 | 0.549 | 0.026 |
| lois2018_pbr28_MNI152_2mm | Molecular | TSPO | 0.001 | > 0.999 | 0.069 | > 0.999 | 0.273 | > 0.999 | 0.018 | > 0.999 | 0.144 | > 0.999 | 0.041 | > 0.999 |
| lukow2022_ro154513_MNI152_2mm | Molecular | GABAa | 0.370 | > 0.999 | 0.366 | > 0.999 | 0.515 | > 0.999 | 0.459 | 0.026 | 0.252 | > 0.999 | 0.440 | 0.026 |
| malen2022_raclopride_MNI152_2mm | Molecular | D2 | 0.201 | > 0.999 | 0.080 | > 0.999 | 0.168 | > 0.999 | 0.102 | > 0.999 | 0.330 | 0.026 | 0.100 | > 0.999 |
| margulies2016_fcgradient01_fsLR_32k | Brain function | FC Grad01 | 0.276 | > 0.999 | 0.029 | > 0.999 | 0.147 | > 0.999 | 0.399 | 0.026 | 0.357 | > 0.999 | 0.309 | > 0.999 |
| margulies2016_fcgradient02_fsLR_32k | Brain function | FC Grad02 | -0.208 | > 0.999 | -0.502 | > 0.999 | -0.266 | > 0.999 | -0.286 | > 0.999 | 0.009 | > 0.999 | -0.444 | 0.026 |
| margulies2016_fcgradient03_fsLR_32k | Brain function | FC Grad03 | 0.296 | 0.568 | 0.228 | > 0.999 | 0.191 | > 0.999 | 0.060 | > 0.999 | 0.189 | > 0.999 | 0.076 | > 0.999 |
| margulies2016_fcgradient04_fsLR_32k | Brain function | FC Grad04 | -0.078 | > 0.999 | 0.098 | > 0.999 | 0.219 | > 0.999 | -0.176 | > 0.999 | -0.167 | > 0.999 | -0.022 | > 0.999 |
| margulies2016_fcgradient05_fsLR_32k | Brain function | FC Grad05 | 0.280 | > 0.999 | 0.149 | > 0.999 | 0.258 | > 0.999 | 0.362 | > 0.999 | 0.345 | > 0.999 | 0.348 | 0.103 |
| margulies2016_fcgradient06_fsLR_32k | Brain function | FC Grad06 | -0.082 | > 0.999 | -0.146 | > 0.999 | -0.220 | > 0.999 | -0.010 | > 0.999 | 0.176 | > 0.999 | 0.013 | > 0.999 |

| | | | | | | | | | | | | | | |
|---|---|---|---|---|---|---|---|---|---|---|---|---|---|---|
| margulies2016_fcgradient07_fsLR_32k | Brain function | FC Grad07 | 0.071 | > 0.999 | 0.136 | > 0.999 | 0.165 | > 0.999 | -0.130 | > 0.999 | 0.014 | > 0.999 | -0.113 | > 0.999 |
| margulies2016_fcgradient08_fsLR_32k | Brain function | FC Grad08 | -0.296 | > 0.999 | -0.403 | 0.490 | -0.329 | > 0.999 | -0.143 | > 0.999 | 0.033 | > 0.999 | -0.425 | 0.026 |
| margulies2016_fcgradient09_fsLR_32k | Brain function | FC Grad09 | 0.361 | 0.103 | 0.456 | 0.026 | 0.401 | 0.929 | 0.143 | > 0.999 | -0.010 | > 0.999 | 0.379 | 0.026 |
| margulies2016_fcgradient10_fsLR_32k | Brain function | FC Grad10 | 0.032 | > 0.999 | 0.131 | > 0.999 | 0.101 | > 0.999 | -0.006 | > 0.999 | -0.032 | > 0.999 | 0.059 | > 0.999 |
| mueller2013_intersubjvar_fsLR_164k | Brain function | RSFC intvar | 0.536 | 0.026 | 0.284 | > 0.999 | 0.262 | > 0.999 | 0.496 | 0.026 | 0.553 | 0.026 | 0.433 | 0.026 |
| naganawa2020_lsn3172176_MNI152_1mm | Molecular | M1 | -0.065 | > 0.999 | -0.165 | > 0.999 | -0.203 | > 0.999 | 0.228 | > 0.999 | 0.090 | > 0.999 | 0.234 | 0.464 |
| neurosynth_cogpc1_MNI152_2mm | Cognition | PC1 Cognition | 0.010 | > 0.999 | 0.096 | > 0.999 | -0.144 | > 0.999 | -0.044 | > 0.999 | -0.144 | > 0.999 | 0.031 | > 0.999 |
| norgaard2021_flumazenil_fsaverage_164k | Molecular | GABAa | 0.099 | > 0.999 | 0.024 | > 0.999 | 0.125 | > 0.999 | 0.103 | > 0.999 | 0.114 | > 0.999 | 0.157 | > 0.999 |
| normandin2015_omar_MNI152_1mm | Molecular | CB1 | 0.207 | > 0.999 | 0.177 | > 0.999 | 0.197 | > 0.999 | 0.473 | 0.026 | 0.274 | > 0.999 | 0.447 | 0.026 |
| radnakrishnan2018_gsk215083_MNI152_1mm | Molecular | 5-HT6 | 0.136 | > 0.999 | -0.031 | > 0.999 | 0.009 | > 0.999 | 0.331 | 0.568 | 0.325 | 0.361 | 0.289 | 0.645 |
| raichle_cbf_fsLR_164k | Physiology | Blood flow | 0.175 | > 0.999 | 0.361 | > 0.999 | 0.217 | > 0.999 | -0.167 | > 0.999 | -0.010 | > 0.999 | 0.136 | > 0.999 |
| raichle_cbv_fsLR_164k | Physiology | Blood vol | -0.249 | > 0.999 | -0.079 | > 0.999 | 0.010 | > 0.999 | -0.167 | > 0.999 | -0.061 | > 0.999 | -0.112 | > 0.999 |

| | | | | | | | | | | | | | | |
|---|---|---|---|---|---|---|---|---|---|---|---|---|---|---|
| raichle_cmr02_fsLR_164k | Metabolism | Oxygen | -0.034 | > 0.999 | -0.030 | > 0.999 | -0.189 | > 0.999 | -0.331 | > 0.999 | -0.091 | > 0.999 | -0.115 | > 0.999 |
| raichle_cmrglc_fsLR_164k | Metabolism | Glucose | 0.104 | > 0.999 | 0.084 | > 0.999 | -0.089 | > 0.999 | -0.214 | > 0.999 | -0.138 | > 0.999 | 0.000 | > 0.999 |
| reardon2018_scalinghcp_civet_41k | Brain structure | Expansion | 0.304 | > 0.999 | 0.070 | > 0.999 | 0.053 | > 0.999 | 0.211 | > 0.999 | 0.606 | 0.026 | 0.015 | > 0.999 |
| reardon2018_scalingnih_civet_41k | Brain structure | Expansion | 0.313 | 0.052 | -0.027 | > 0.999 | -0.056 | > 0.999 | 0.464 | 0.026 | 0.500 | 0.026 | 0.238 | > 0.999 |
| reardon2018_scalingpnc_civet_41k | Brain structure | Expansion | 0.376 | 0.026 | 0.114 | > 0.999 | 0.042 | > 0.999 | 0.280 | > 0.999 | 0.470 | 0.026 | 0.128 | > 0.999 |
| rosaneto_abp688_MNI152_1mm | Molecular | mGluR5 | 0.358 | 0.026 | 0.210 | > 0.999 | 0.222 | > 0.999 | 0.519 | 0.026 | 0.480 | 0.026 | 0.576 | 0.026 |
| sandiego2015_flb457_MNI152_1mm | Molecular | D2 | 0.088 | > 0.999 | -0.027 | > 0.999 | 0.135 | > 0.999 | 0.410 | 0.568 | 0.277 | > 0.999 | 0.252 | > 0.999 |
| sasaki2012_fepe2i_MNI152_1mm | Molecular | DAT | 0.198 | > 0.999 | 0.203 | > 0.999 | 0.186 | > 0.999 | -0.032 | > 0.999 | 0.033 | > 0.999 | 0.068 | > 0.999 |
| satterthwaite2014_meancbf_MNI152_1mm | Physiology | Blood flow | 0.337 | > 0.999 | 0.293 | > 0.999 | 0.271 | > 0.999 | 0.286 | > 0.999 | 0.235 | > 0.999 | 0.379 | 0.464 |
| savli2012_altanserin_MNI152_3mm | Molecular | 5-HT2a | 0.149 | > 0.999 | -0.044 | > 0.999 | 0.057 | > 0.999 | 0.351 | > 0.999 | 0.476 | 0.026 | 0.207 | > 0.999 |
| savli2012_dasb_MNI152_3mm | Molecular | 5-HTT | -0.040 | > 0.999 | 0.177 | > 0.999 | 0.373 | > 0.999 | -0.021 | > 0.999 | 0.107 | > 0.999 | 0.028 | > 0.999 |
| savli2012_p943_MNI152_3mm | Molecular | 5-HT1b | 0.239 | > 0.999 | 0.175 | > 0.999 | 0.101 | > 0.999 | 0.128 | > 0.999 | 0.262 | > 0.999 | 0.231 | > 0.999 |

| | | | | | | | | | | | | | | |
|---|---|---|---|---|---|---|---|---|---|---|---|---|---|---|
| savli2012_way100635_MNI152_3mm | Molecular | 5-HT1a | 0.193 | > 0.999 | 0.128 | > 0.999 | 0.289 | > 0.999 | 0.398 | 0.284 | 0.354 | 0.748 | 0.240 | > 0.999 |
| smart2019_abp688_MNI152_1mm | Molecular | mGluR5 | 0.346 | 0.026 | 0.223 | > 0.999 | 0.201 | > 0.999 | 0.566 | 0.026 | 0.536 | 0.026 | 0.564 | 0.026 |
| smith2017_flb457_MNI152_1mm | Molecular | D2 | 0.075 | > 0.999 | -0.054 | > 0.999 | 0.092 | > 0.999 | 0.424 | 0.181 | 0.295 | > 0.999 | 0.275 | > 0.999 |
| sydnor2021_SAaxis_fsLR_32k | Brain function | Sensory-association | 0.439 | > 0.999 | 0.297 | > 0.999 | 0.374 | > 0.999 | 0.466 | 0.026 | 0.396 | 0.671 | 0.407 | 0.026 |
| tuominen_feobv_MNI152_2mm | Molecular | VAChT | 0.289 | > 0.999 | 0.523 | > 0.999 | 0.498 | > 0.999 | 0.269 | > 0.999 | 0.050 | > 0.999 | 0.462 | 0.026 |
| turtonen2020_carfentanil_MNI152_1mm | Molecular | MOR | 0.548 | 0.052 | 0.490 | > 0.999 | 0.504 | > 0.999 | 0.544 | 0.026 | 0.390 | > 0.999 | 0.597 | 0.026 |
| vijay2018_ly2795050_MNI152_2mm | Molecular | KOR | 0.319 | > 0.999 | 0.455 | > 0.999 | 0.339 | > 0.999 | 0.397 | 0.026 | 0.086 | > 0.999 | 0.508 | 0.026 |
| wey2016_martinostat_MNI152_2mm | Molecular | HDAC | 0.155 | > 0.999 | 0.142 | > 0.999 | 0.058 | > 0.999 | 0.075 | > 0.999 | 0.070 | > 0.999 | 0.191 | > 0.999 |
| xu2020_FChomology_fsLR_32k | Brain function | Functional homology | -0.305 | 0.439 | -0.053 | > 0.999 | -0.089 | > 0.999 | -0.496 | 0.026 | -0.503 | 0.129 | -0.414 | 0.026 |
| xu2020_evoexp_fsLR_32k | Brain structure | Expansion | 0.215 | > 0.999 | -0.023 | > 0.999 | -0.013 | > 0.999 | 0.414 | 0.026 | 0.314 | > 0.999 | 0.316 | > 0.999 |

**Table S12. Statistics of the latent dimensions for subcortical and cerebellar volumes.**

| | Main/Mixed sample | | | | | | | | | |
|---|---|---|---|---|---|---|---|---|---|---|
| | 1st LD | | 2nd LD | | 3rd LD | | 4th LD | | 5th LD | |
| Split | r | p-value | r | p-value | r | p-value | r | p-value | r | p-value |
| 1 | 0.27 | **0.005** | 0.13 | **0.005** | 0.08 | **0.005** | 0.03 | 0.459 | 0.03 | 0.739 |
| 2 | 0.25 | **0.005** | 0.14 | **0.005** | 0.10 | **0.005** | 0.08 | **0.005** | 0.02 | 0.889 |
| 3 | 0.27 | **0.005** | 0.18 | **0.005** | 0.11 | **0.005** | 0.08 | **0.005** | 0.03 | 0.424 |
| 4 | 0.27 | **0.005** | 0.17 | **0.005** | 0.08 | **0.025** | 0.06 | 0.115 | 0.003 | >0.99 |
| 5 | 0.26 | **0.005** | 0.16 | **0.005** | 0.11 | **0.005** | 0.05 | 0.105 | 0.06 | **0.03** |

p-values are corrected using the Bonferroni method over 5 comparisons; significant p-values are in bold; r: Pearson's correlation coefficient

### 4. S4: appendix: Supplementary figures

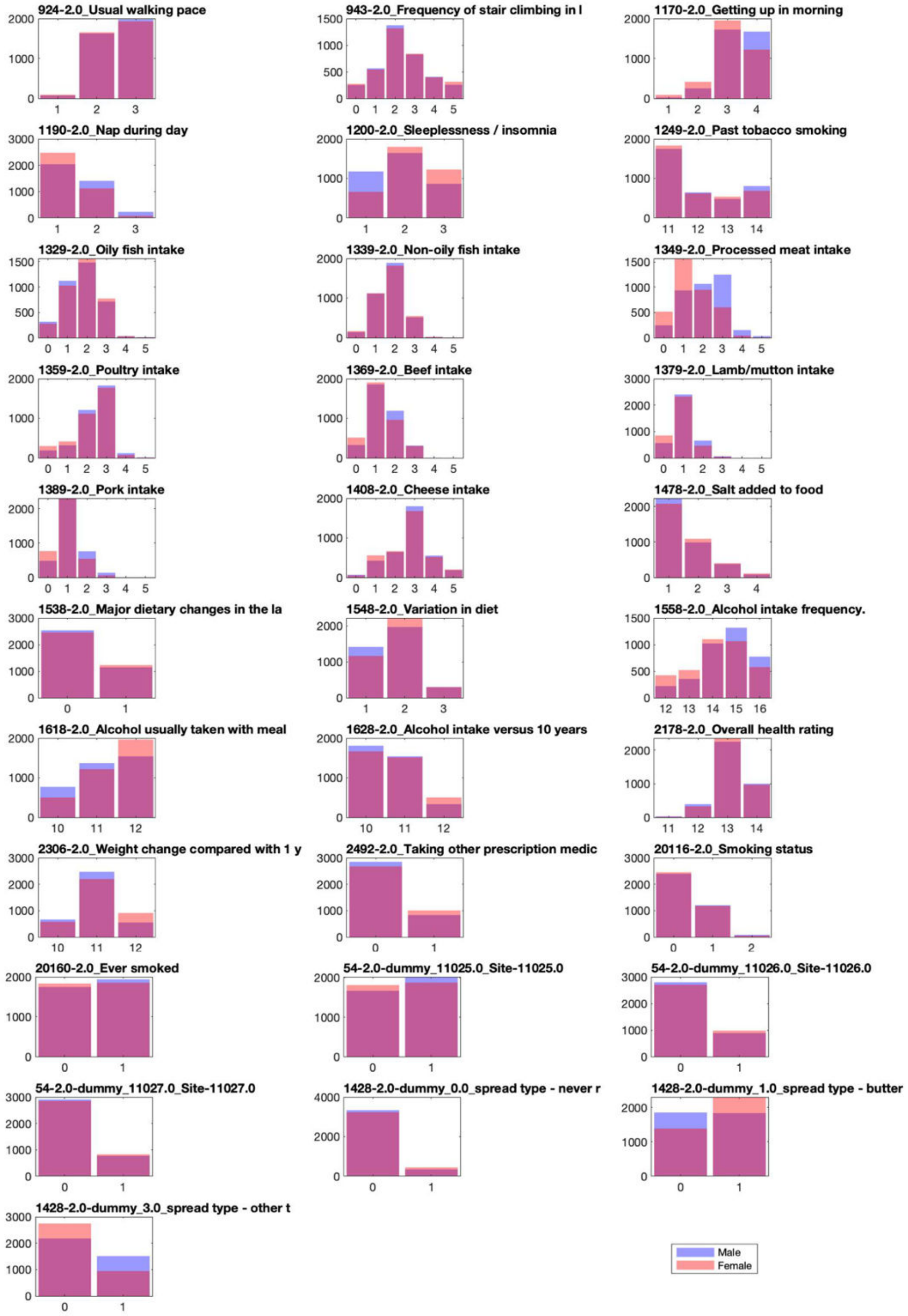

**Figure S1. Distribution of risk factors. Part 1.** Bar plots depicting the distribution of risk factors for women and men in the analyzed samples.

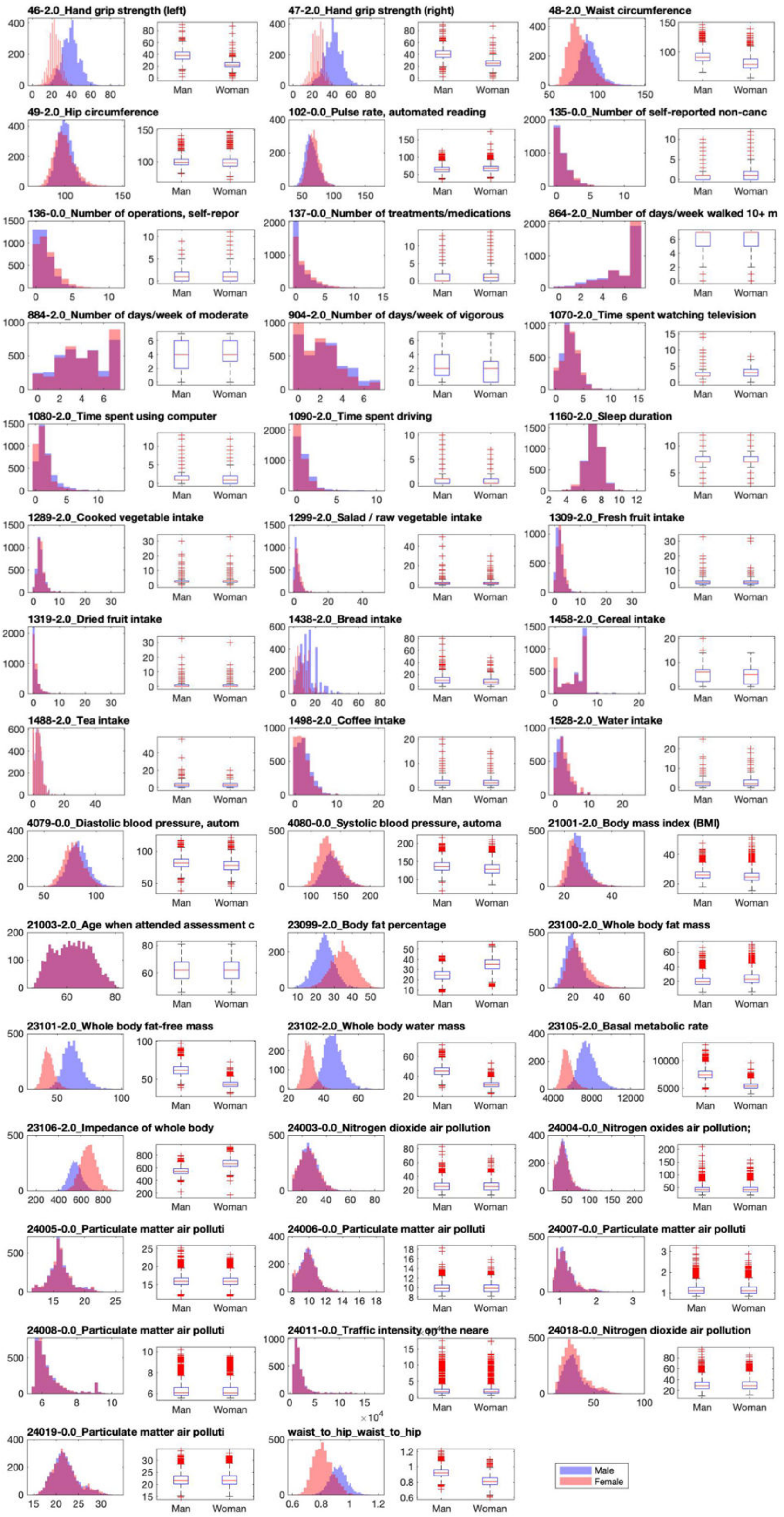

**Figure S2. Distribution of risk factors. Part 2.** Histograms and boxplots depicting the distribution of risk factors for women and men in the analyzed samples. Data on each boxplot corresponds to the histogram to the left.

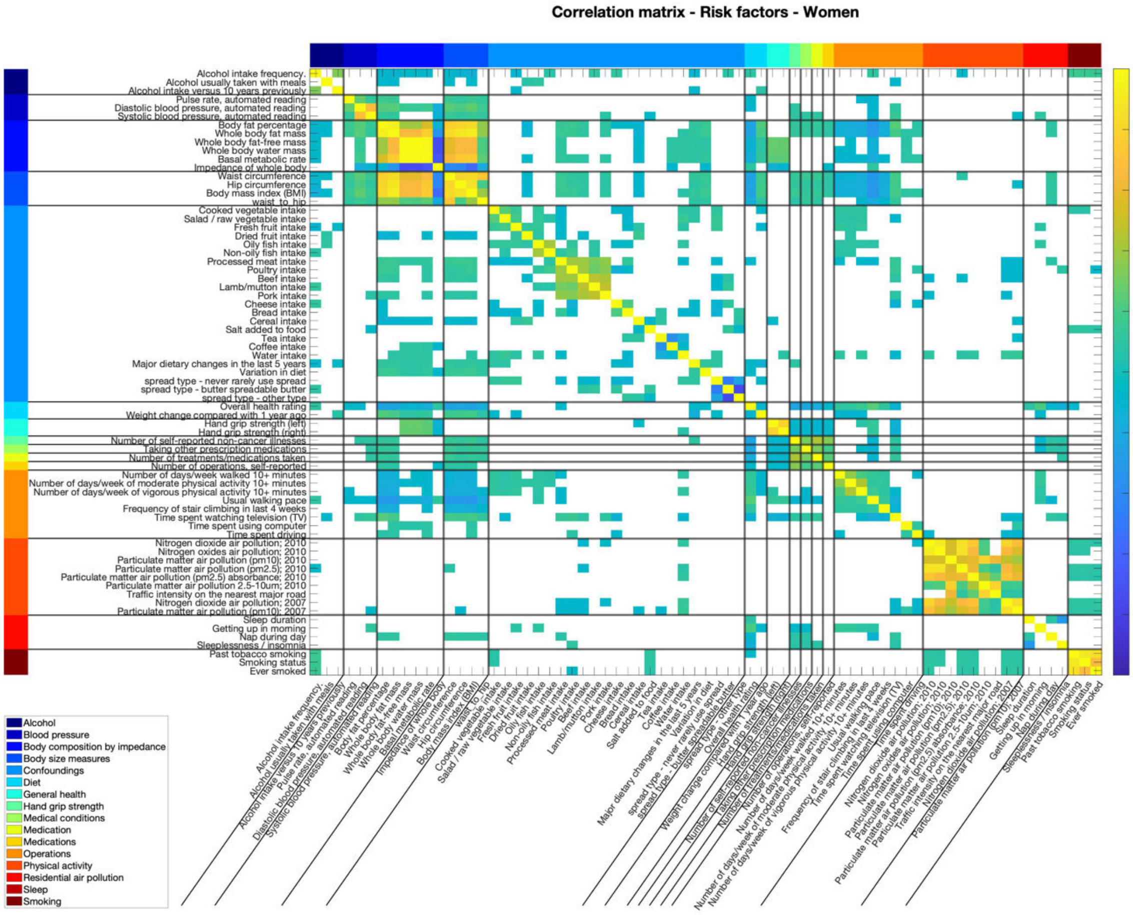

**Figure S3. Correlation matrix of risk factors in women.** Intercorrelation among risk factors in the subsample of women. Color bar depicts the Pearson's correlation coefficient. Only significant associations after correction for multiple comparisons (Bonferroni) are shown. The colored bands indicate the categories to which the risk factors belong.

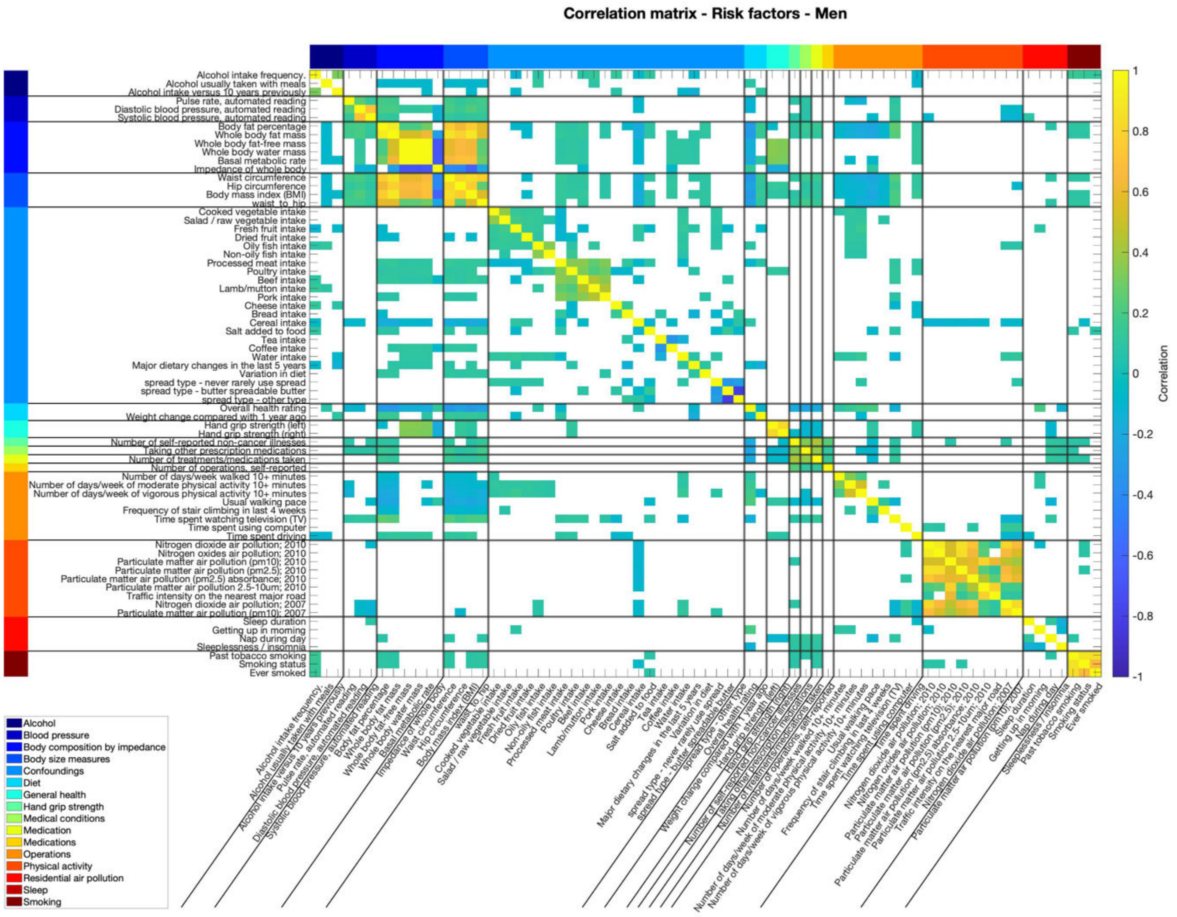

**Figure S4. Correlation matrix of risk factors in men.** Intercorrelation among risk factors in the subsample of men. Color bar depicts the Pearson's correlation coefficient. Only significant associations after correction for multiple comparisons (Bonferroni) are shown. The colored bands indicate the categories to which the risk factors belong.

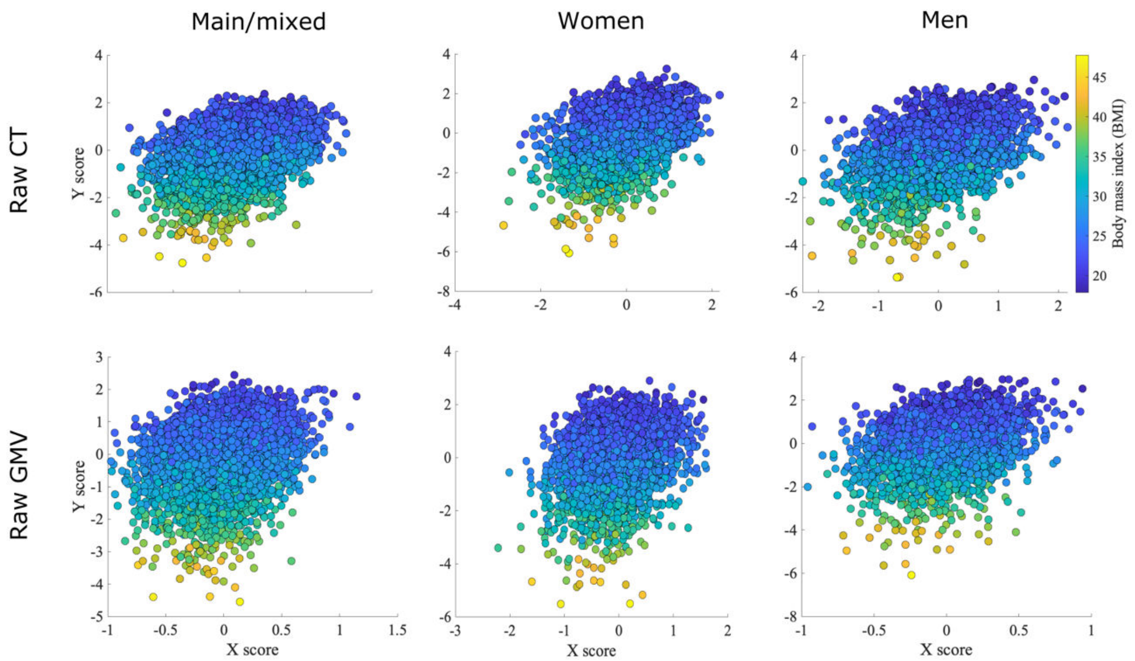

**Figure S5. Latent dimension of cardiometabolic health.** Each scatterplot shows the brain structural (X) and risk factors (Y) scores averaged over the splits for each model. Each dot represents one participant. CT: cortical thickness. GMV: grey matter volume

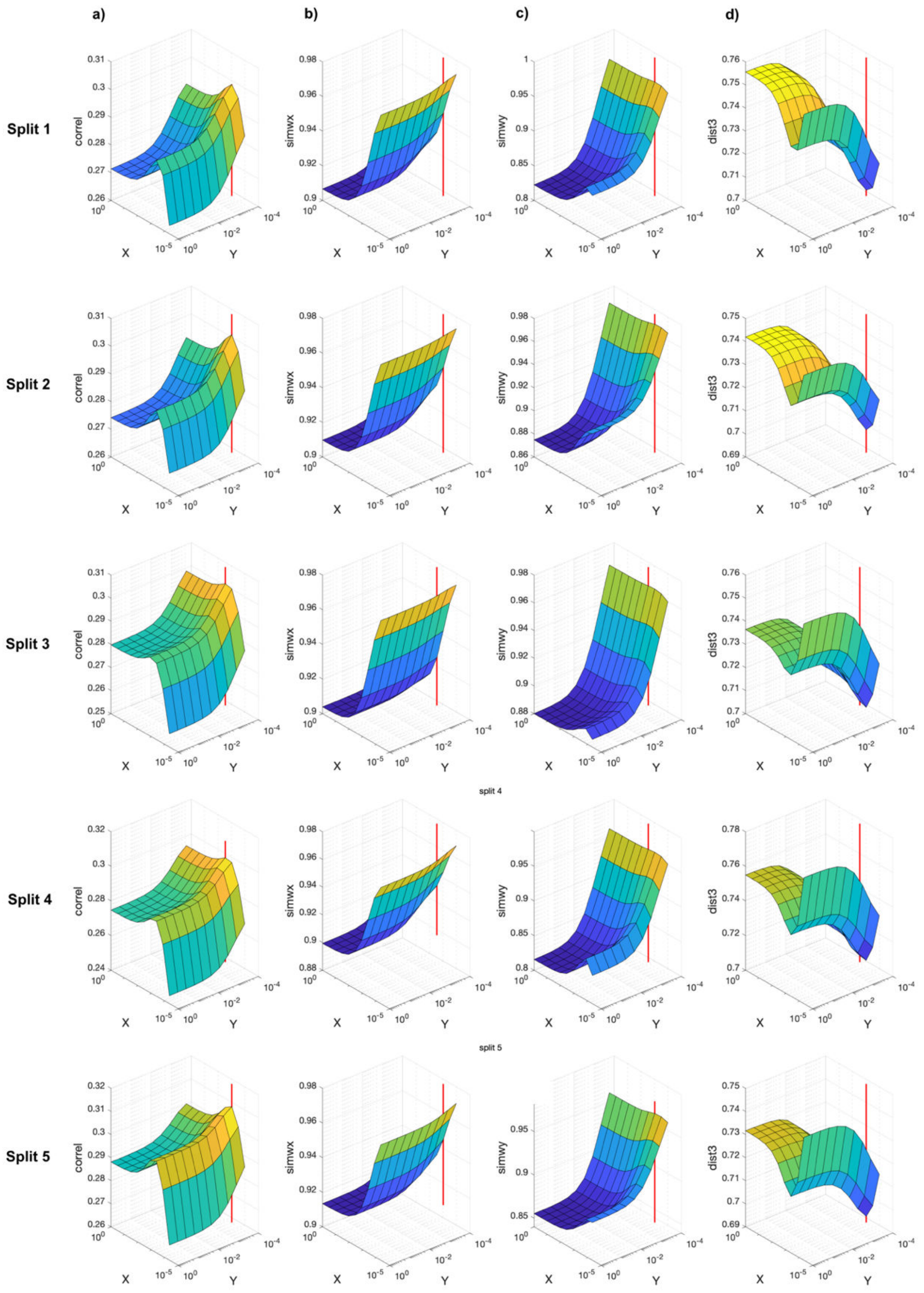

**Figure S6. Model optimization for the latent dimension of cardiometabolic health for raw cortical thickness in the main sample.** The red line indicates the selected model. The z axis represents the test canonical correlation (column a), the similarity of weights in cortical thickness (column b) and risk factors (column c), and the joint generalizability-stability criteria (column d). The x and y axes represent 1 minus the hyperparameters tested for cortical thickness and risk factors, respectively (1-hyperparameter). The x and y axes are shown in logarithmic scale.

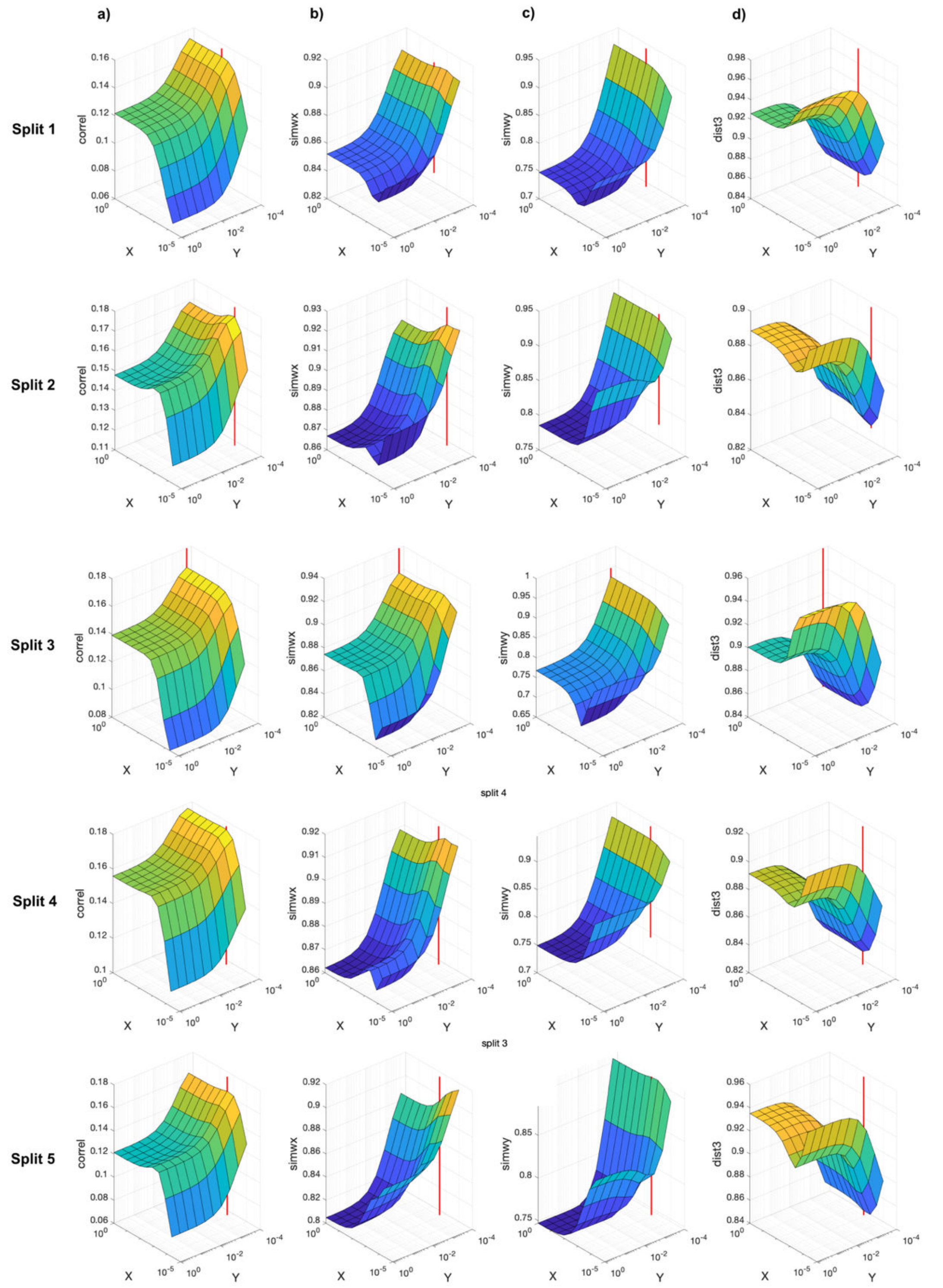

**Figure S7. Model optimization for the latent dimension of cardiometabolic health for raw grey matter volume in the main sample.** The red line indicates the selected model. The z axis represents the test canonical correlation (column a), the similarity of weights in grey matter volume (column b) and risk factors (column c), and the joint generalizability-stability criteria (column d). The x and y axes represent 1 minus the hyperparameters tested for grey matter volume and risk factors, respectively (1-hyperparameter). The x and y axes are shown in logarithmic scale.

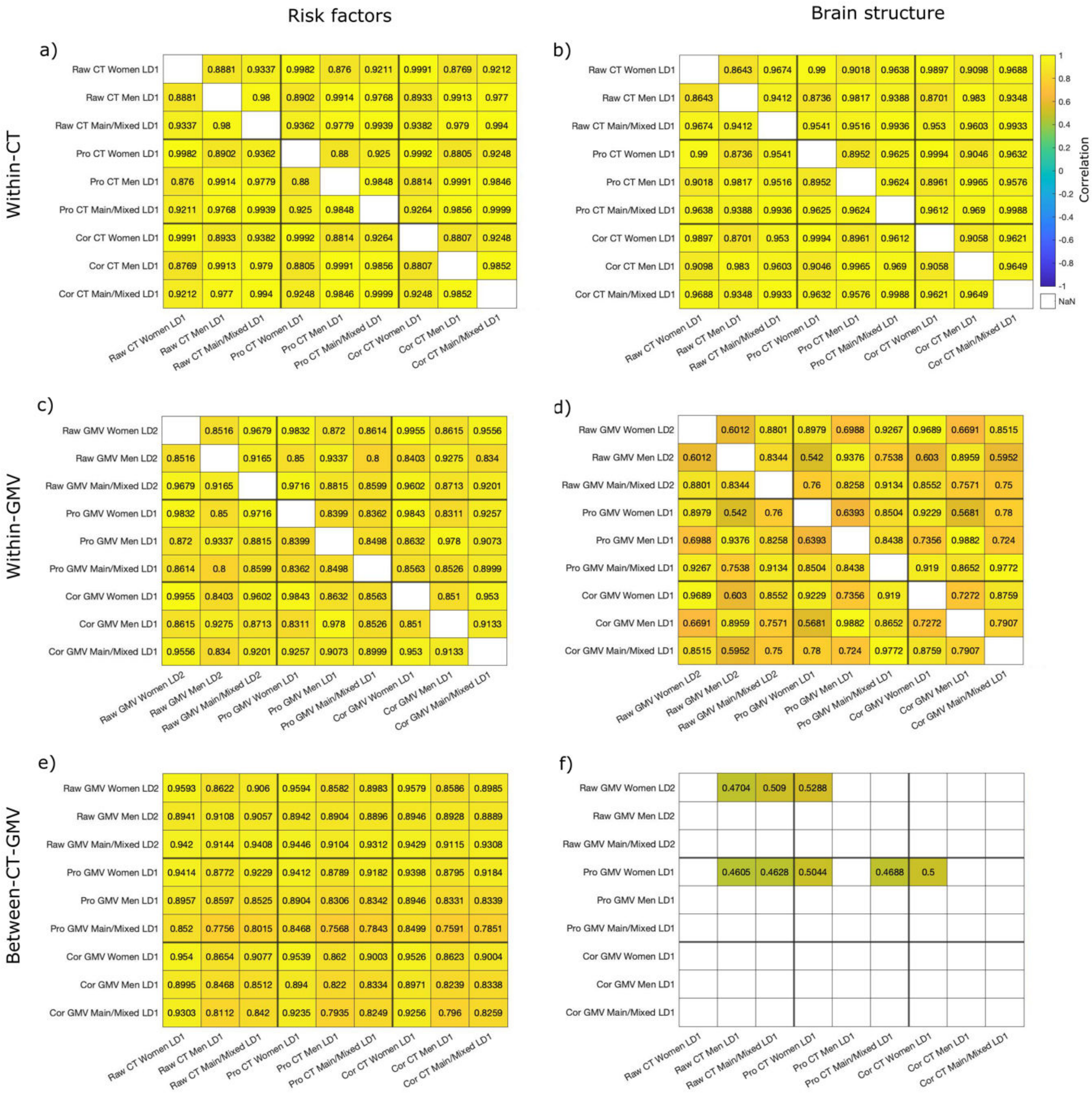

**Figure S8. Comparison of risk factor loadings and brain loadings for the cardiometabolic health latent dimension across samples and brain structural measures.** The first row shows the Spearman correlation of a) risk factors loadings and b) brain loadings for the latent dimension of cardiometabolic health yielded with cortical thickness. The second row shows the Spearman correlation of c) risk factors loadings and d) brain loadings for the latent dimension of cardiometabolic health yielded with grey matter volume. Only comparisons that were significant after Bonferroni correction are shown.

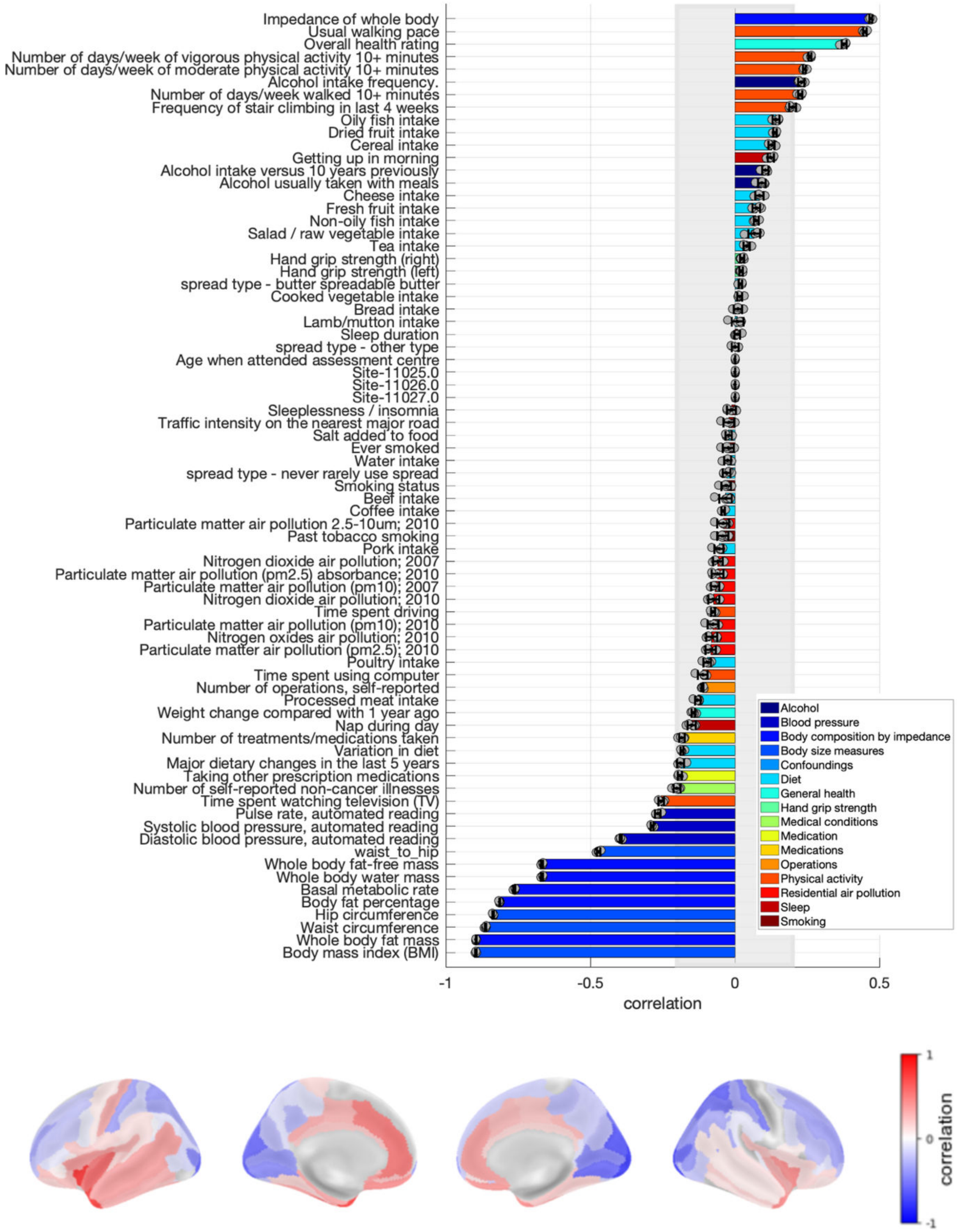

**Figure S9. Loadings of the latent dimension of cardiometabolic health for raw cortical thickness in women.** Risk factors loadings and brain loadings. Shown loadings represent the average over the five outer splits. Error bars depict one standard deviation. The shadowed zone marks loadings between −0.2 and 0.2.

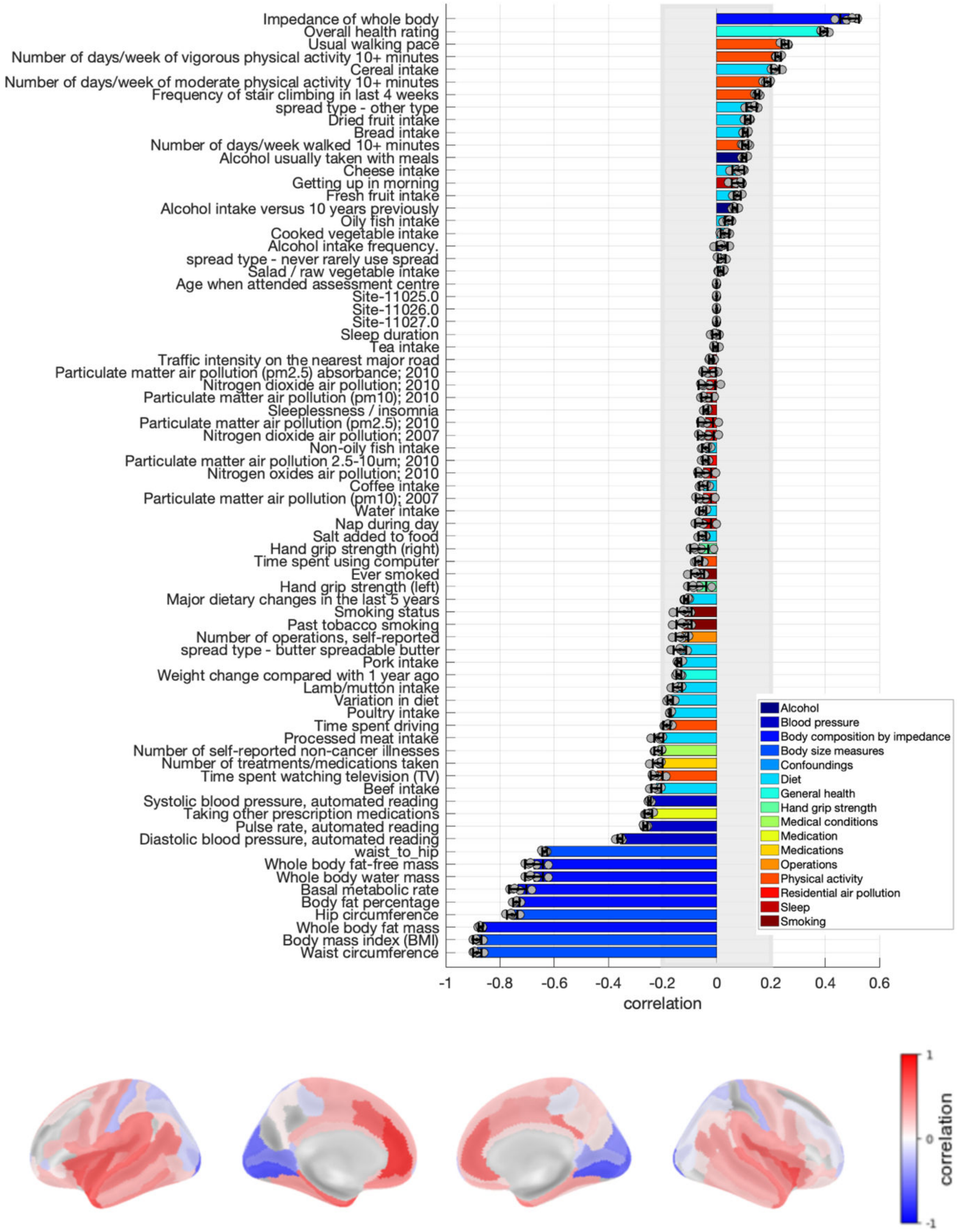

**Figure S10. Loadings of the latent dimension of cardiometabolic health for raw cortical thickness in men.** Risk factors loadings and brain loadings. Shown loadings represent the average over the five outer splits. Error bars depict one standard deviation. The shadowed zone marks loadings between −0.2 and 0.2.

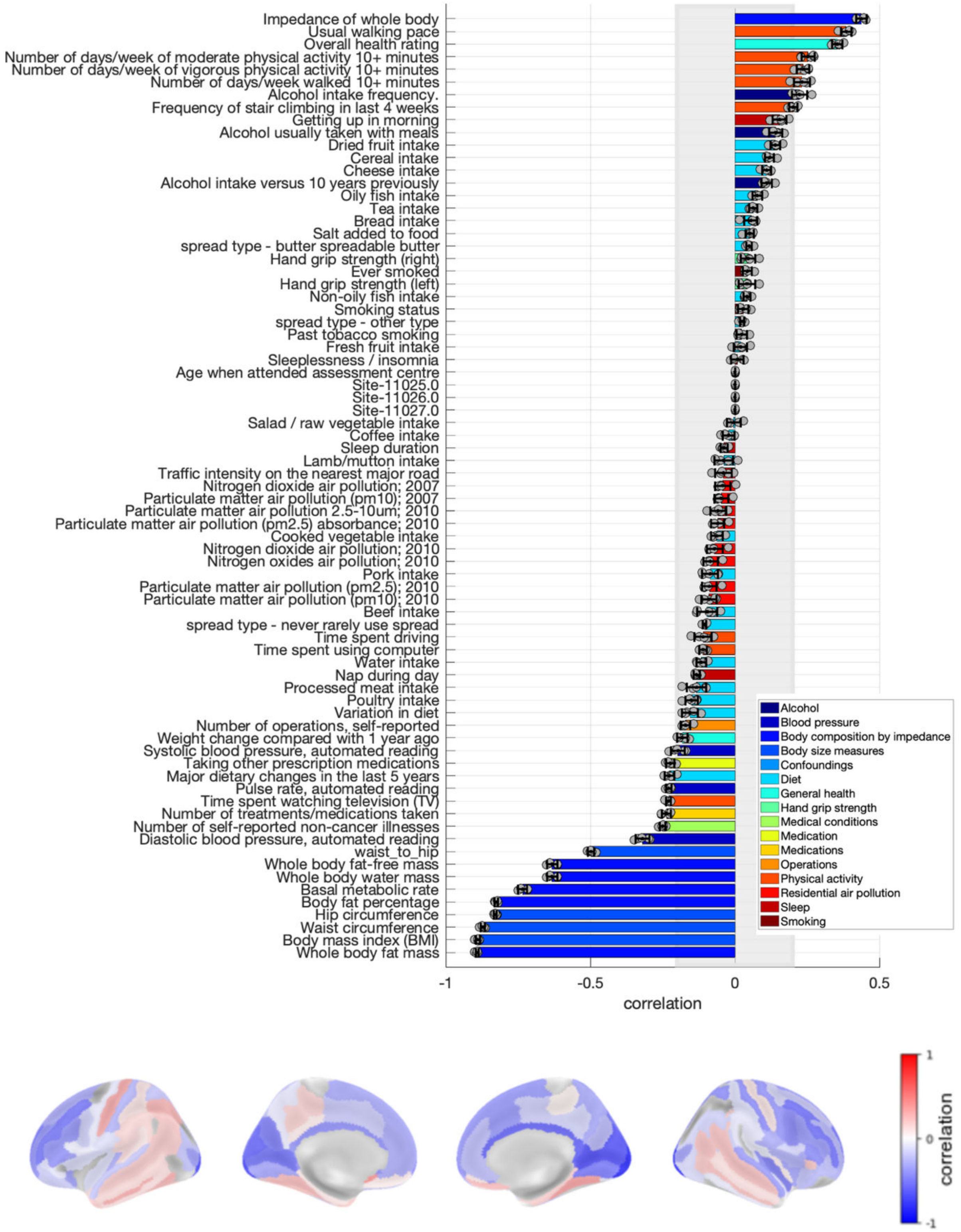

**Figure S11. Loadings of the latent dimension of cardiometabolic health for raw grey matter volume in women.** Risk factors loadings and brain loadings. Shown loadings represent the average over the five outer splits. Error bars depict one standard deviation. The shadowed zone marks loadings between −0.2 and 0.2.

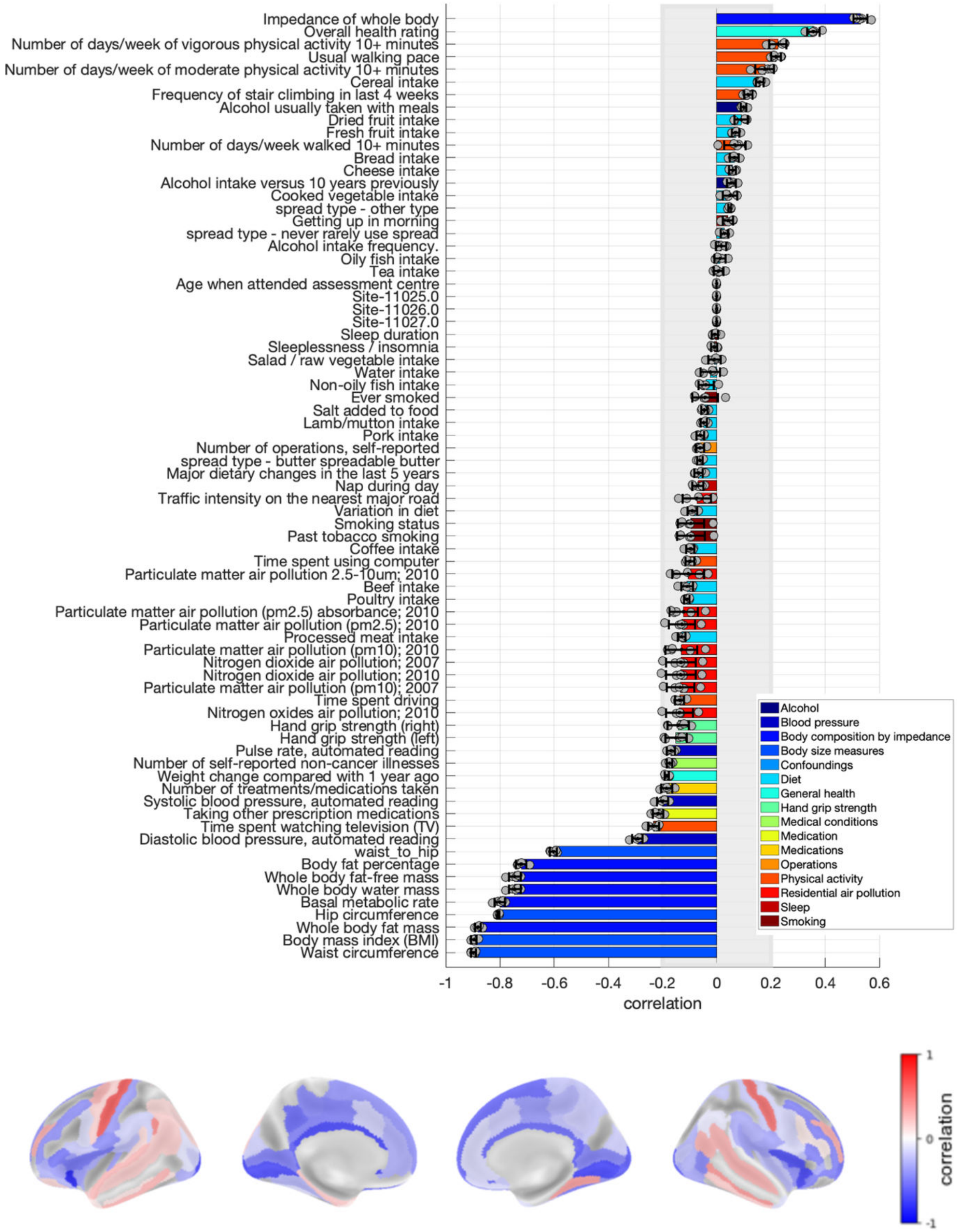

**Figure S12. Loadings of the latent dimension of cardiometabolic health for raw grey matter volume in men.** Risk factors loadings and brain loadings. Shown loadings represent the average over the five outer splits. Error bars depict one standard deviation. The shadowed zone marks loadings between −0.2 and 0.2.

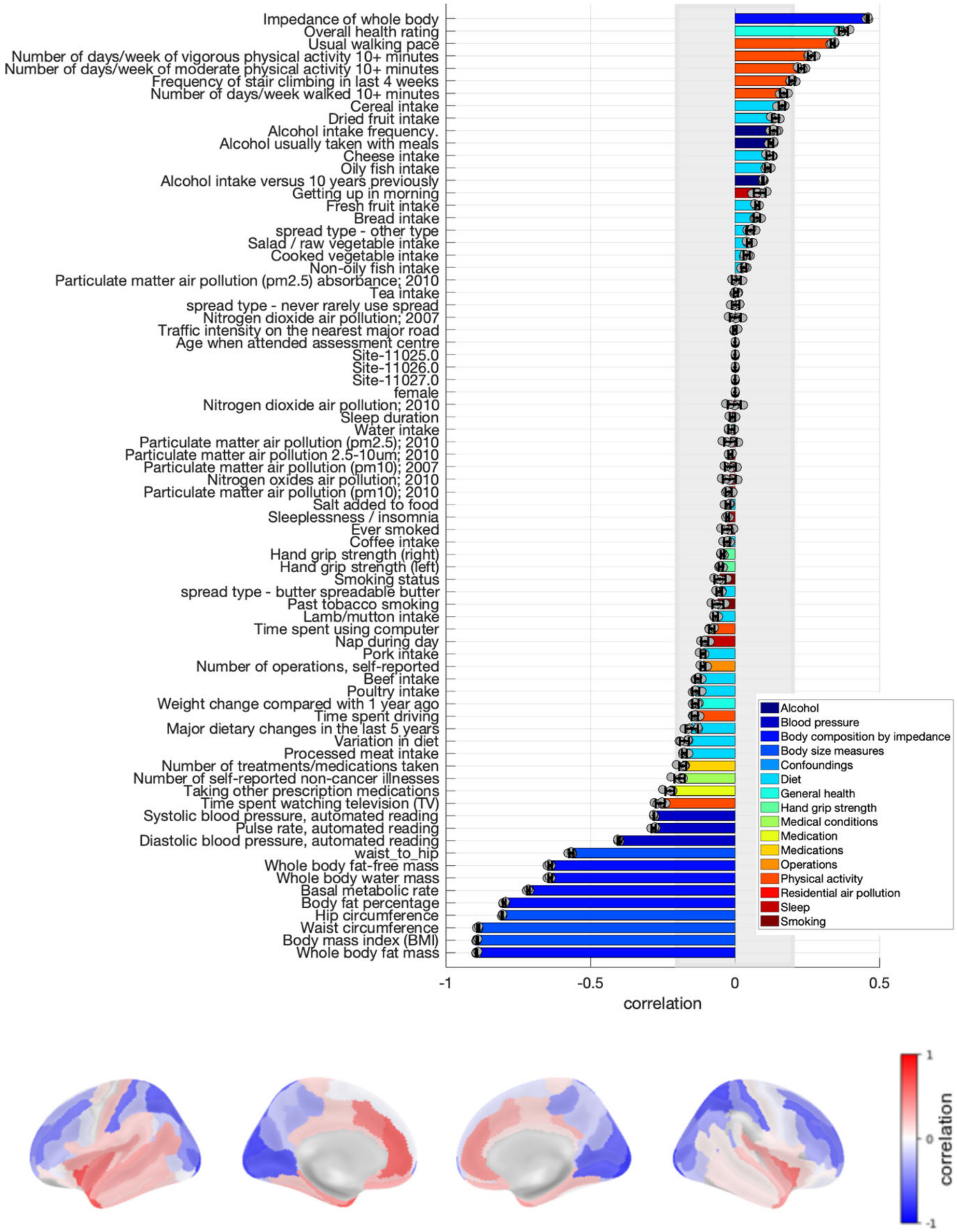

**Figure S13. Loadings of the latent dimension of cardiometabolic health for proportional cortical thickness in the main sample.** Risk factors loadings and brain loadings. Shown loadings represent the average over the five outer splits. Error bars depict one standard deviation. The shadowed zone marks loadings between −0.2 and 0.2.

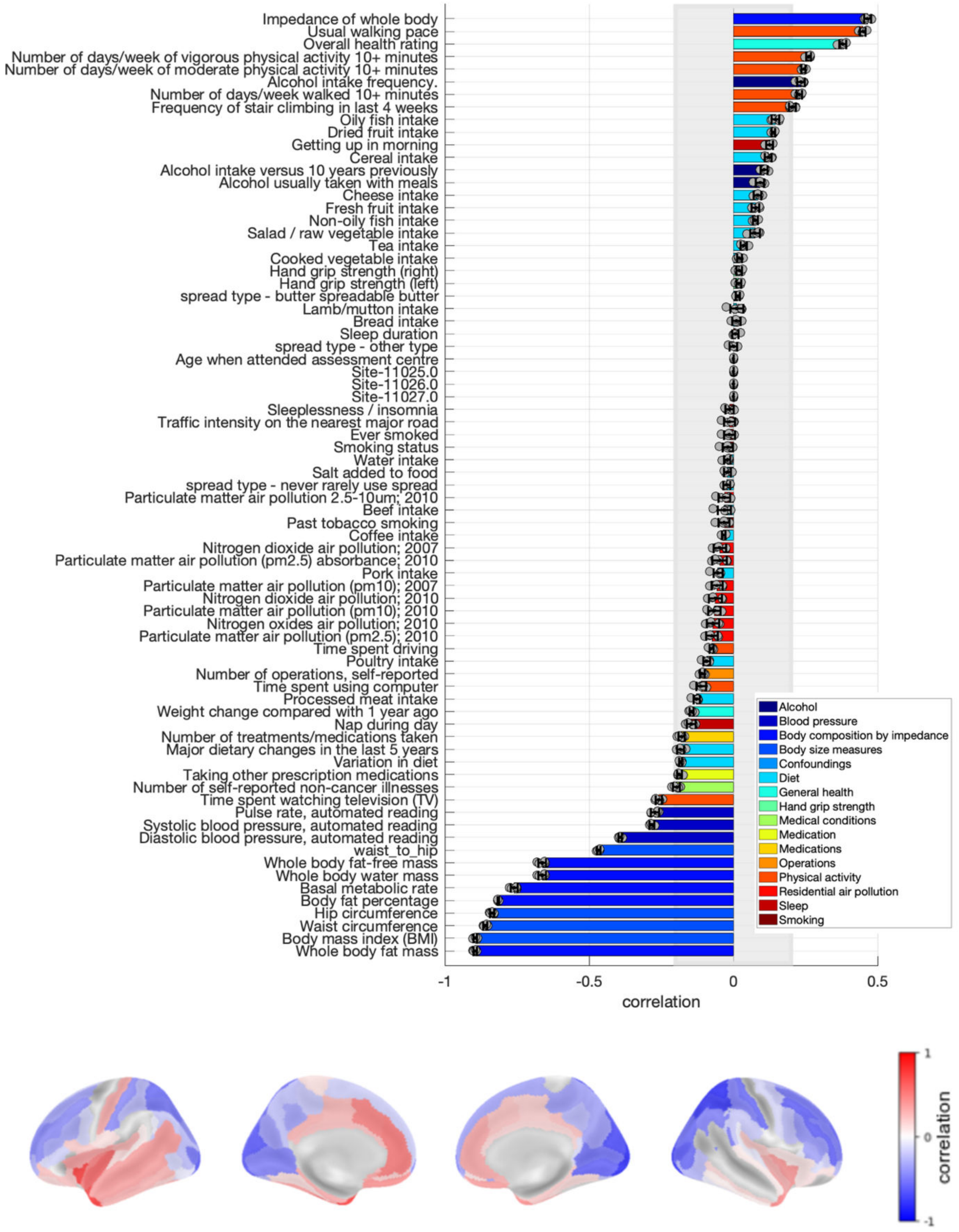

**Figure S14. Loadings of the latent dimension of cardiometabolic health for proportional cortical thickness in women.** Risk factors loadings and brain loadings. Shown loadings represent the average over the five outer splits. Error bars depict one standard deviation. The shadowed zone marks loadings between −0.2 and 0.2.

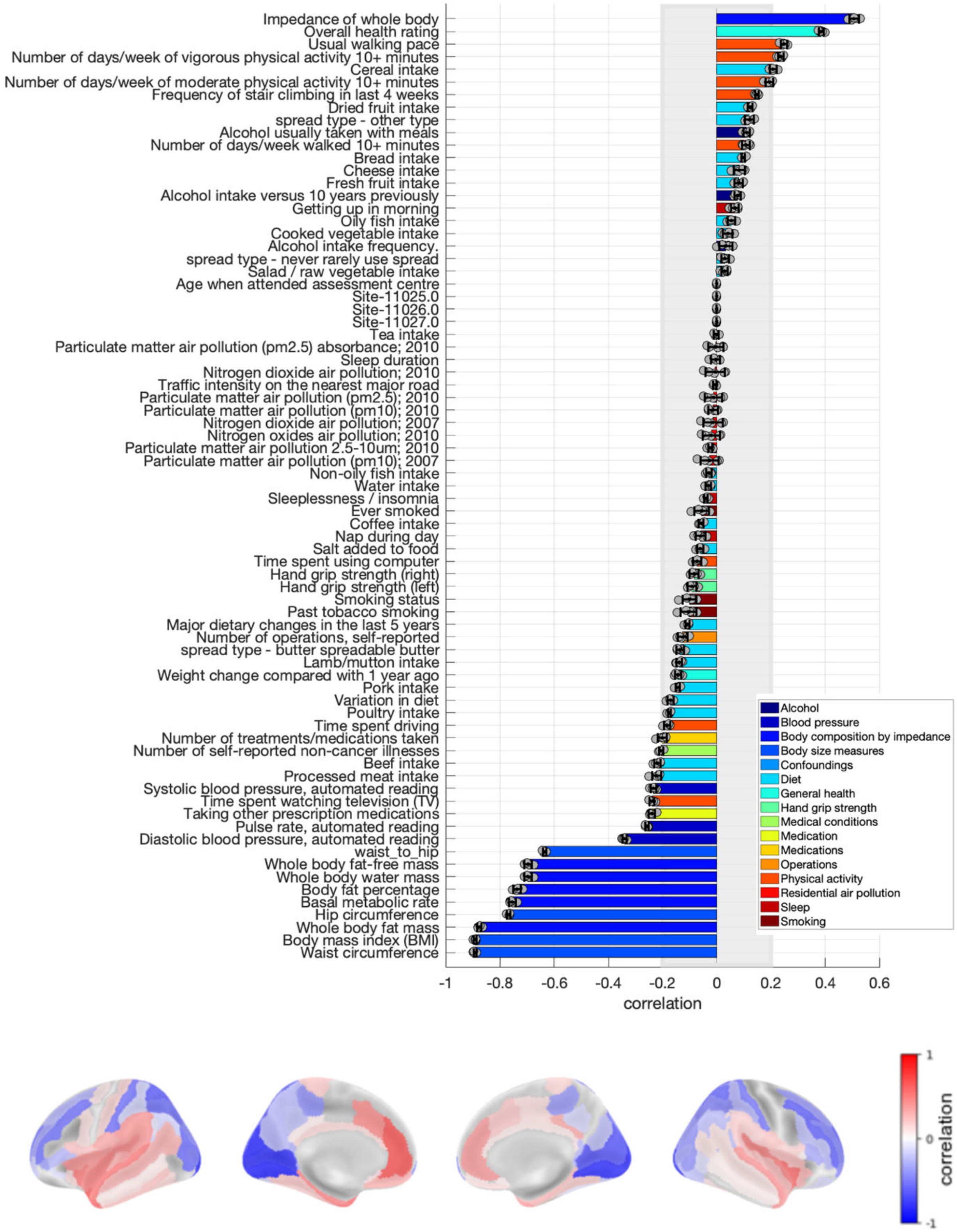

**Figure S15. Loadings of the latent dimension of cardiometabolic health for proportional cortical thickness in men.** Risk factors loadings and brain loadings. Shown loadings represent the average over the five outer splits. Error bars depict one standard deviation. The shadowed zone marks loadings between −0.2 and 0.2.

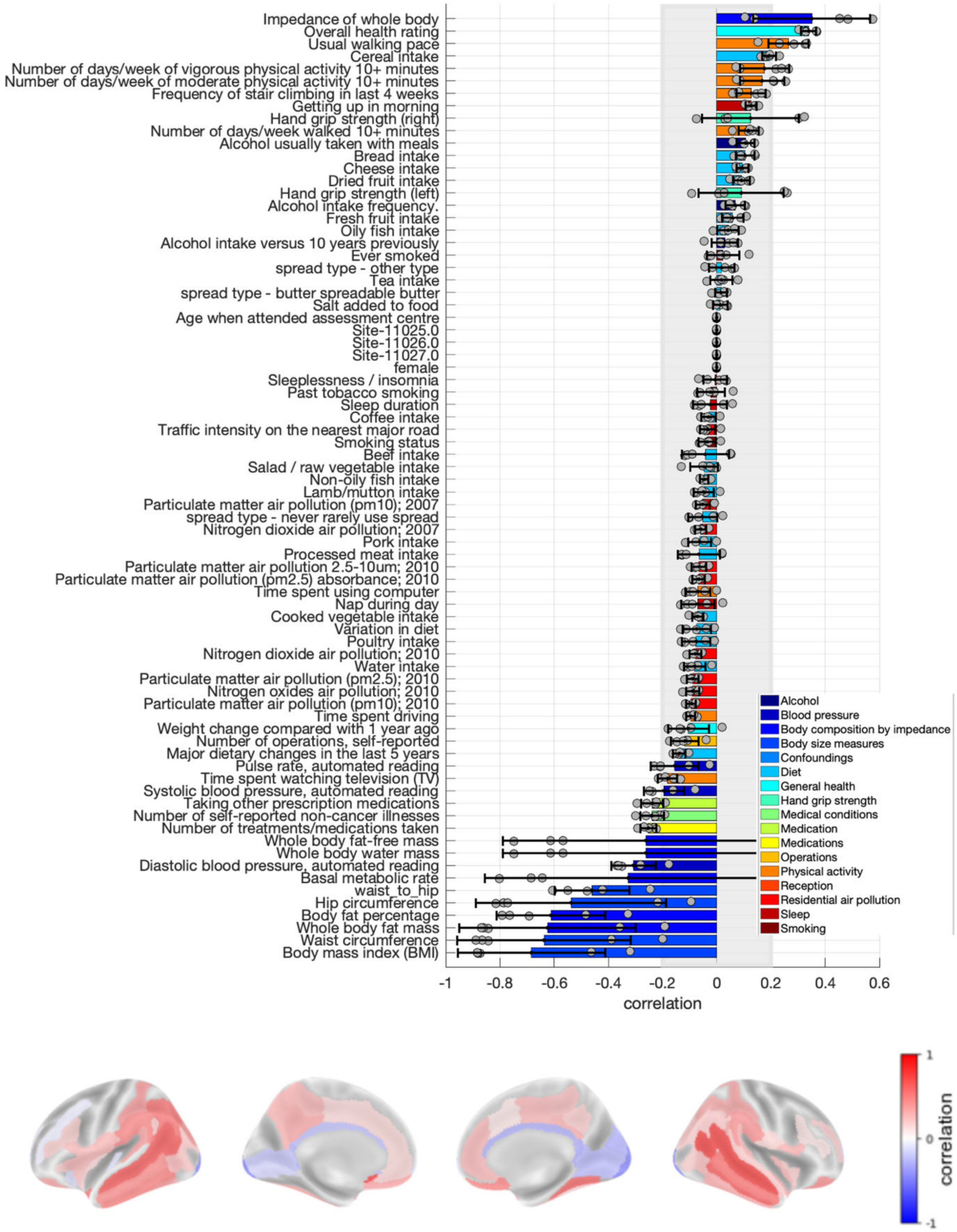

**Figure S16. Loadings of the latent dimension of cardiometabolic health for proportional grey matter volume in the main sample.** Risk factors loadings and brain loadings. Shown loadings represent the average over the five outer splits. Error bars depict one standard deviation. The shadowed zone marks loadings between −0.2 and 0.2.

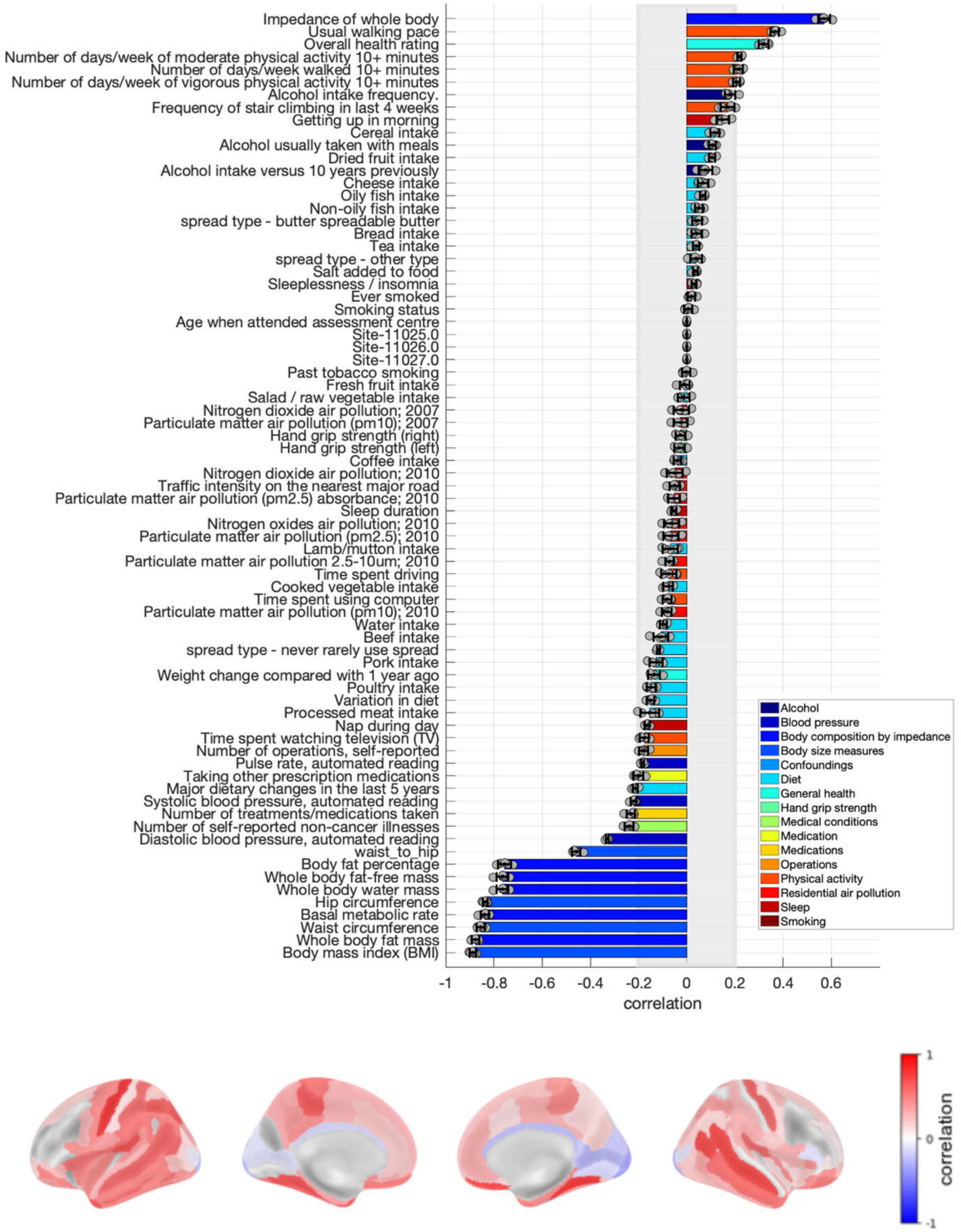

**Figure S17. Loadings of the latent dimension of cardiometabolic health for proportional grey matter volume in women.** Risk factors loadings and brain loadings. Shown loadings represent the average over the five outer splits. Error bars depict one standard deviation. The shadowed zone marks loadings between −0.2 and 0.2.

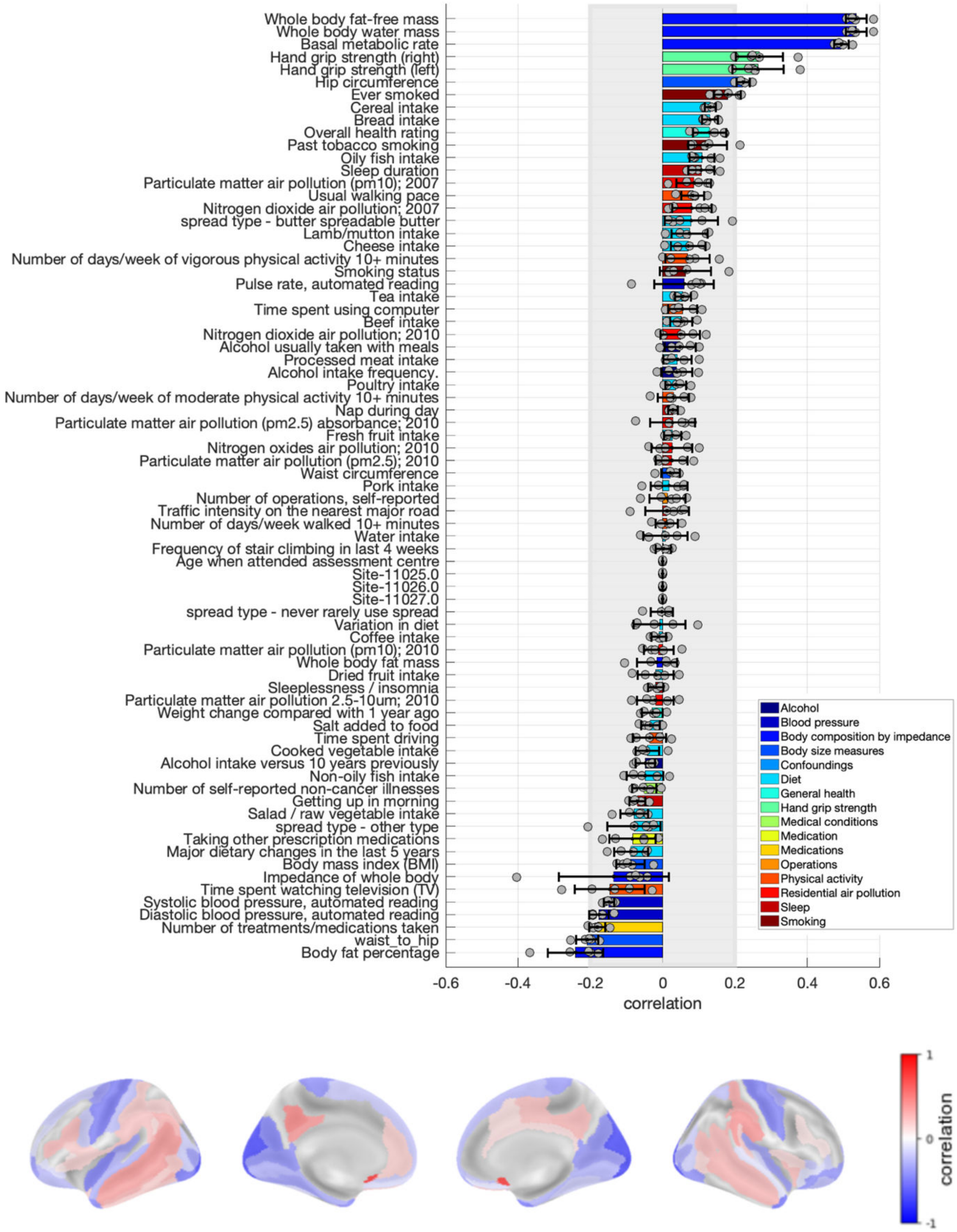

**Figure S18. Loadings of the latent dimension of cardiometabolic health for proportional grey matter volume in men.** Risk factors loadings and brain loadings. Shown loadings represent the average over the five outer splits. Error bars depict one standard deviation. The shadowed zone marks loadings between −0.2 and 0.2.

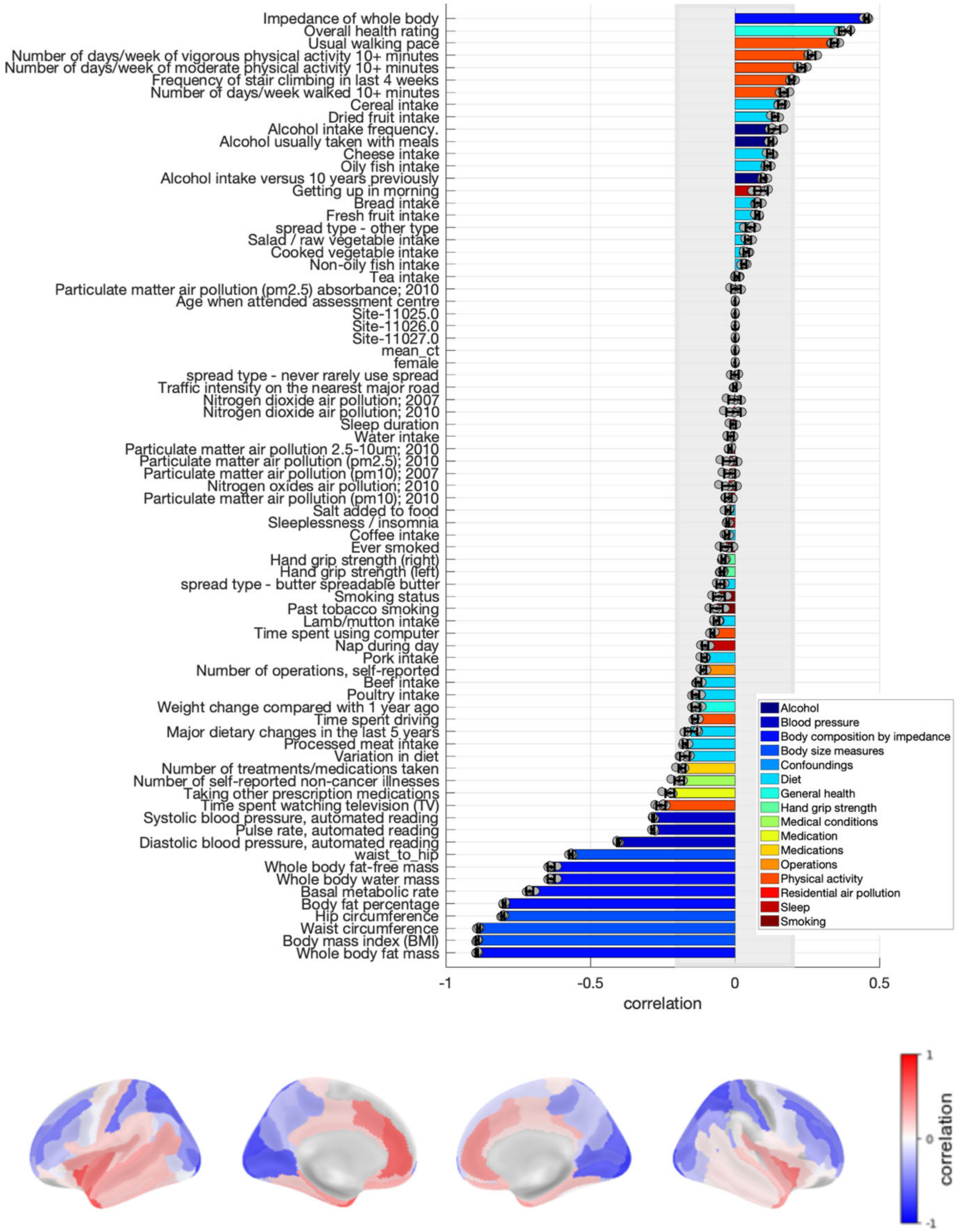

**Figure S19. Loadings of the latent dimension of cardiometabolic health for corrected cortical thickness in the main sample.** Risk factors loadings and brain loadings. Shown loadings represent the average over the five outer splits. Error bars depict one standard deviation. The shadowed zone marks loadings between −0.2 and 0.2.

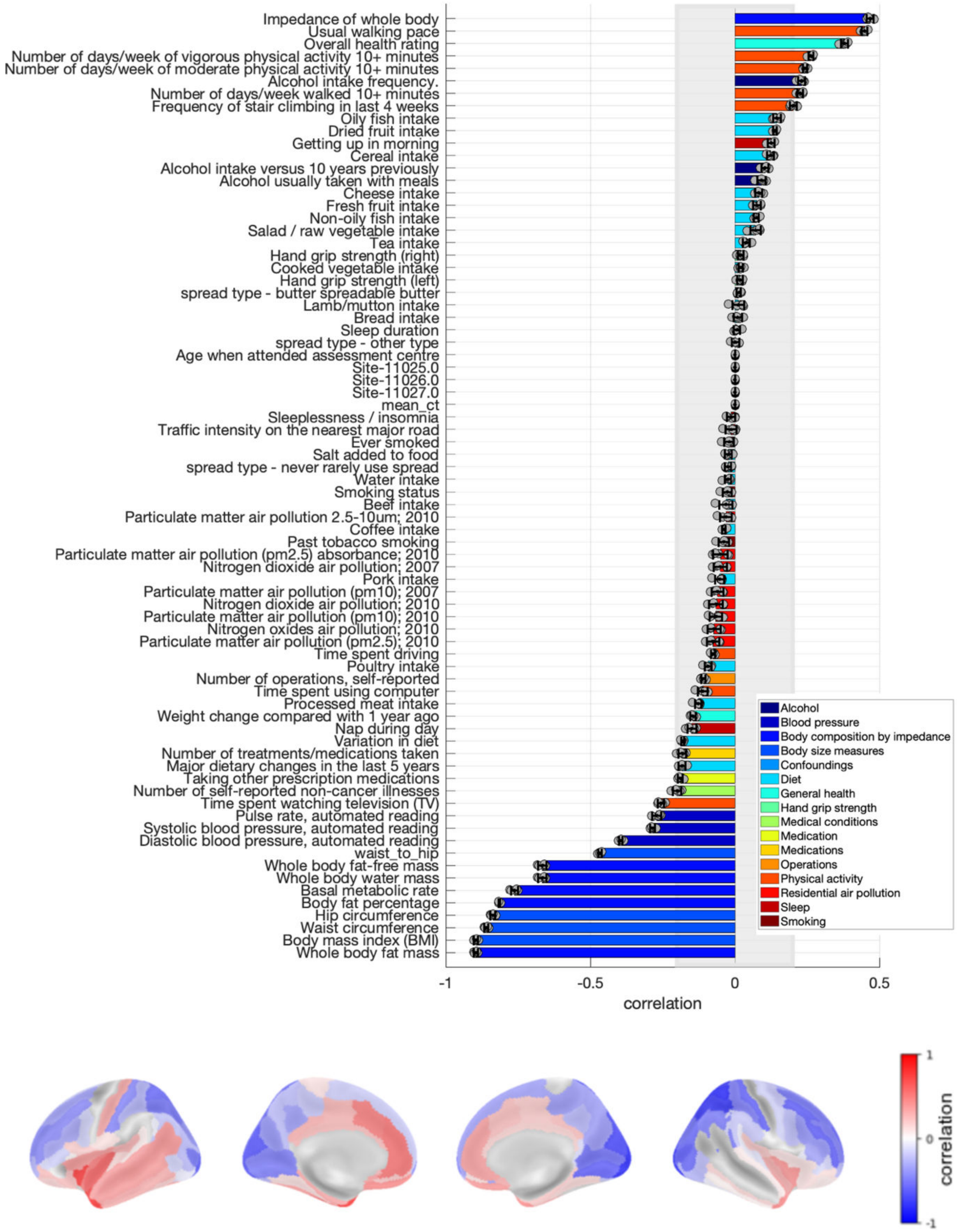

**Figure S20. Loadings of the latent dimension of cardiometabolic health for corrected cortical thickness in women.** Risk factors loadings and brain loadings. Shown loadings represent the average over the five outer splits. Error bars depict one standard deviation. The shadowed zone marks loadings between −0.2 and 0.2.

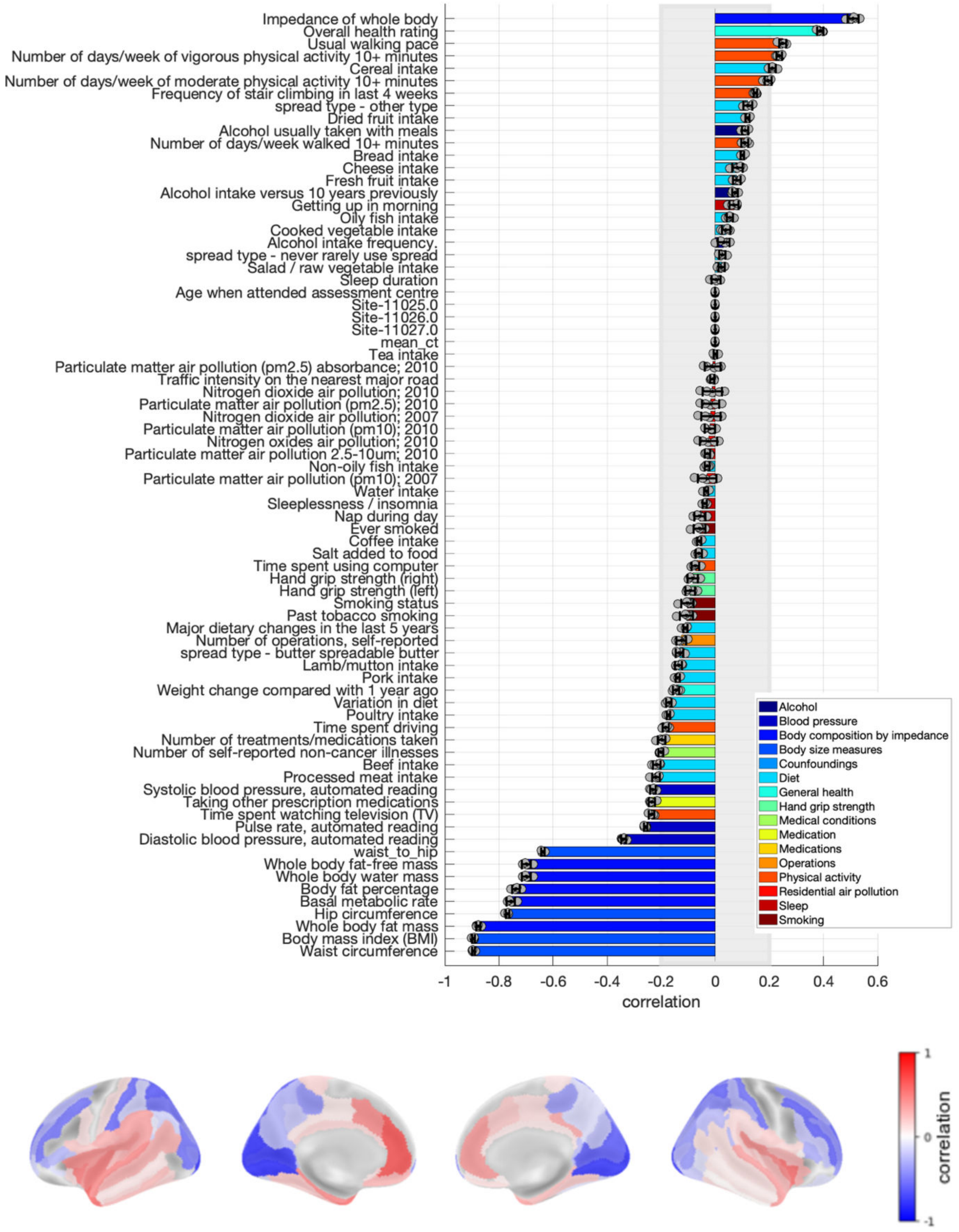

**Figure S21. Loadings of the latent dimension of cardiometabolic health for corrected cortical thickness in men.** Risk factors loadings and brain loadings. Shown loadings represent the average over the five outer splits. Error bars depict one standard deviation. The shadowed zone marks loadings between −0.2 and 0.2.

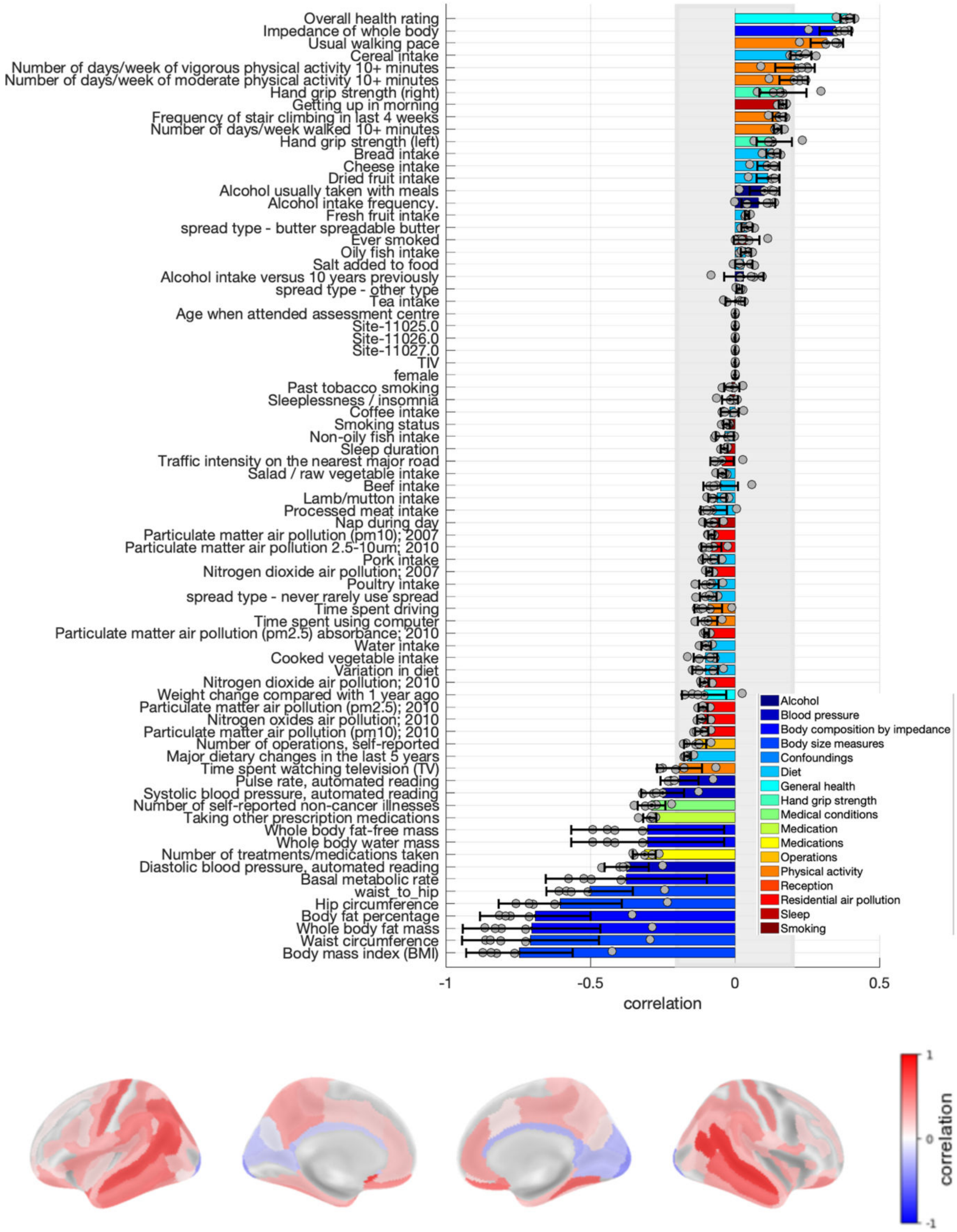

**Figure S22. Loadings of the latent dimension of cardiometabolic health for corrected grey matter volume in the main sample.** Risk factors loadings and brain loadings. Shown loadings represent the average over the five outer splits. Error bars depict one standard deviation. The shadowed zone marks loadings between −0.2 and 0.2.

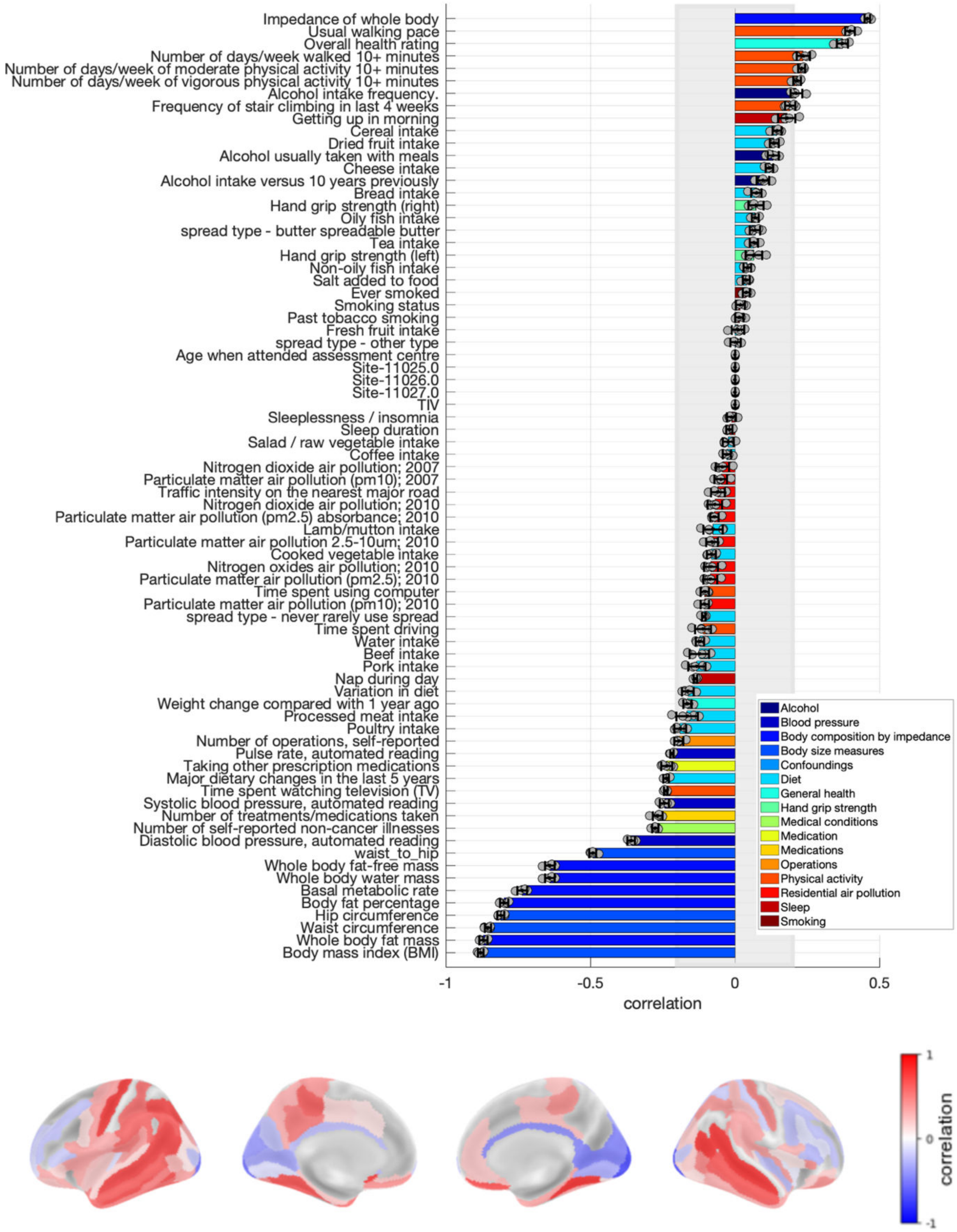

**Figure S23. Loadings of the latent dimension of cardiometabolic health for corrected grey matter volume in women.** Risk factors loadings and brain loadings. Shown loadings represent the average over the five outer splits. Error bars depict one standard deviation. The shadowed zone marks loadings between −0.2 and 0.2.

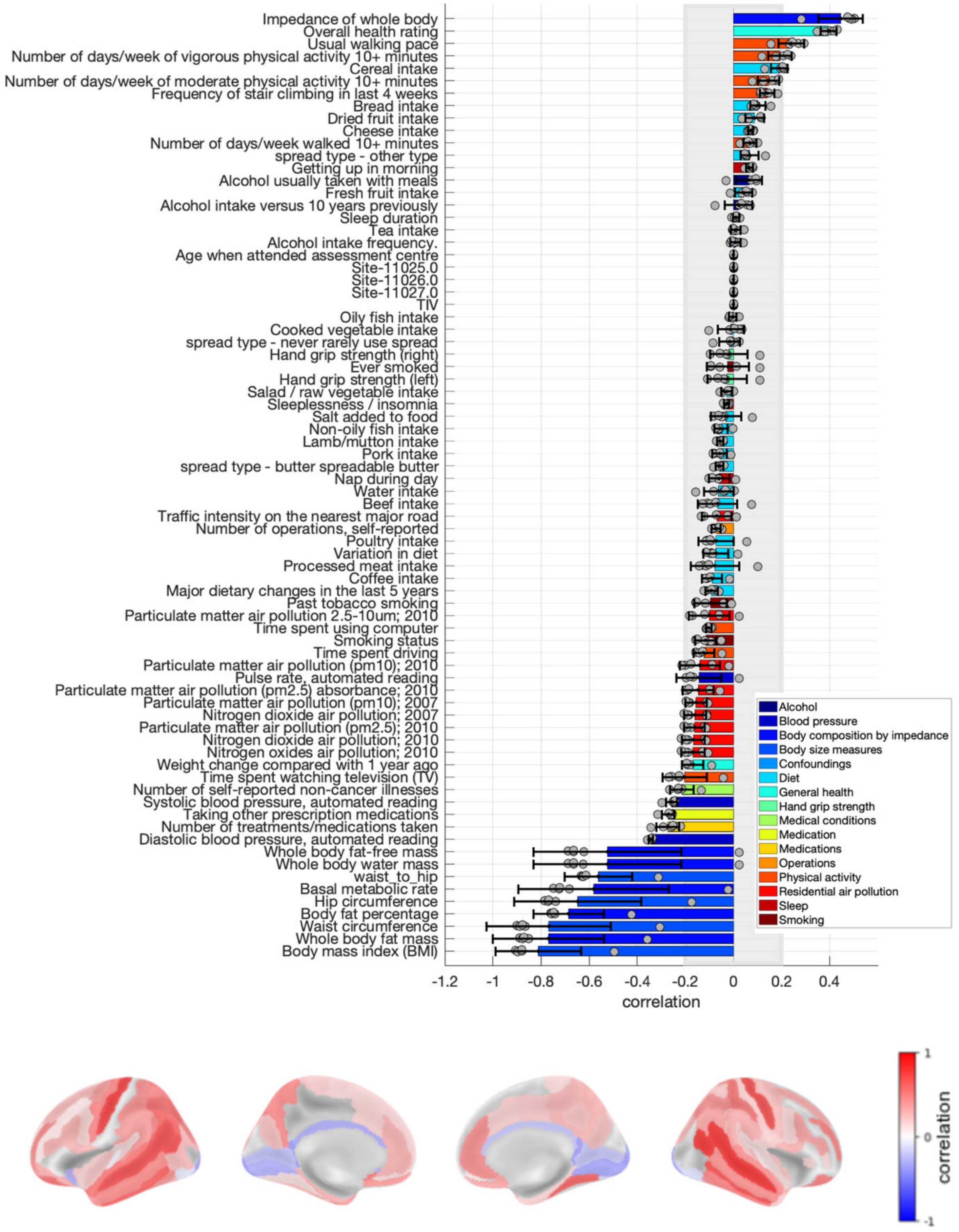

**Figure S24. Loadings of the latent dimension of cardiometabolic health for corrected grey matter volume in men.** Risk factors loadings and brain loadings. Shown loadings represent the average over the five outer splits. Error bars depict one standard deviation. The shadowed zone marks loadings between −0.2 and 0.2.

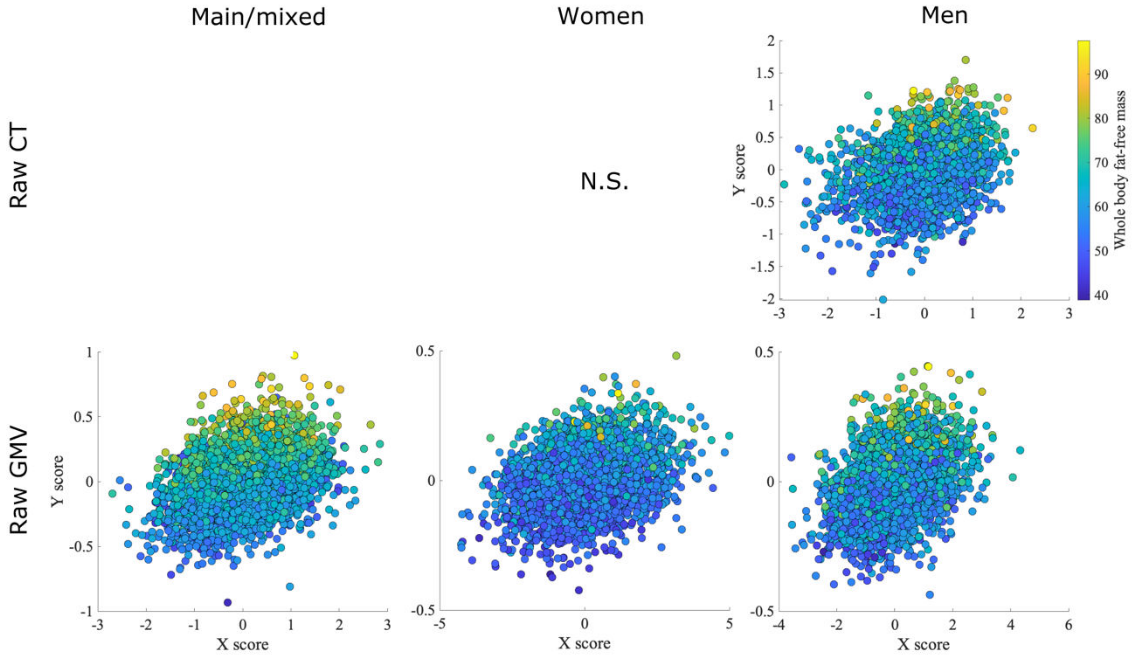

**Figure S25. Latent dimension of physical robustness.** Each scatterplot shows the brain structural (X) and risk factors (Y) scores averaged over the splits for each model. Each dot represents one participant. CT: cortical thickness. GMV: grey matter volume

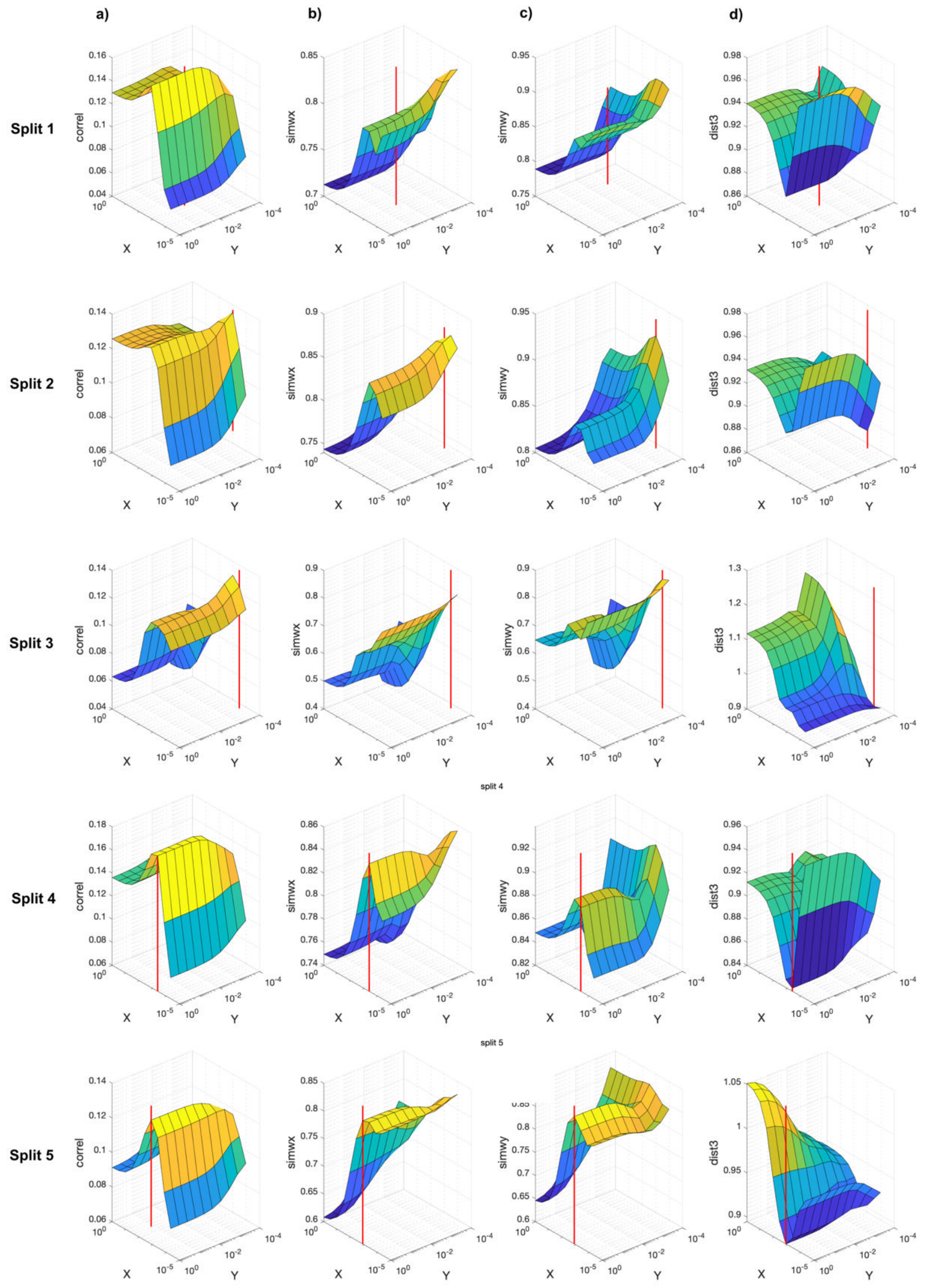

**Figure S26. Model optimization for the latent dimension of physical robustness for raw cortical thickness in the sample of men.** The red line indicates the selected model. The z axis represents the test canonical correlation (column a), the similarity of weights in cortical thickness (column b) and risk factors (column c), and the joint generalizability-stability criteria (column d). The x and y axes represent 1 minus the hyperparameters tested for cortical thickness and risk factors, respectively (1-hyperparameter). The x and y axes are shown in logarithmic scale.

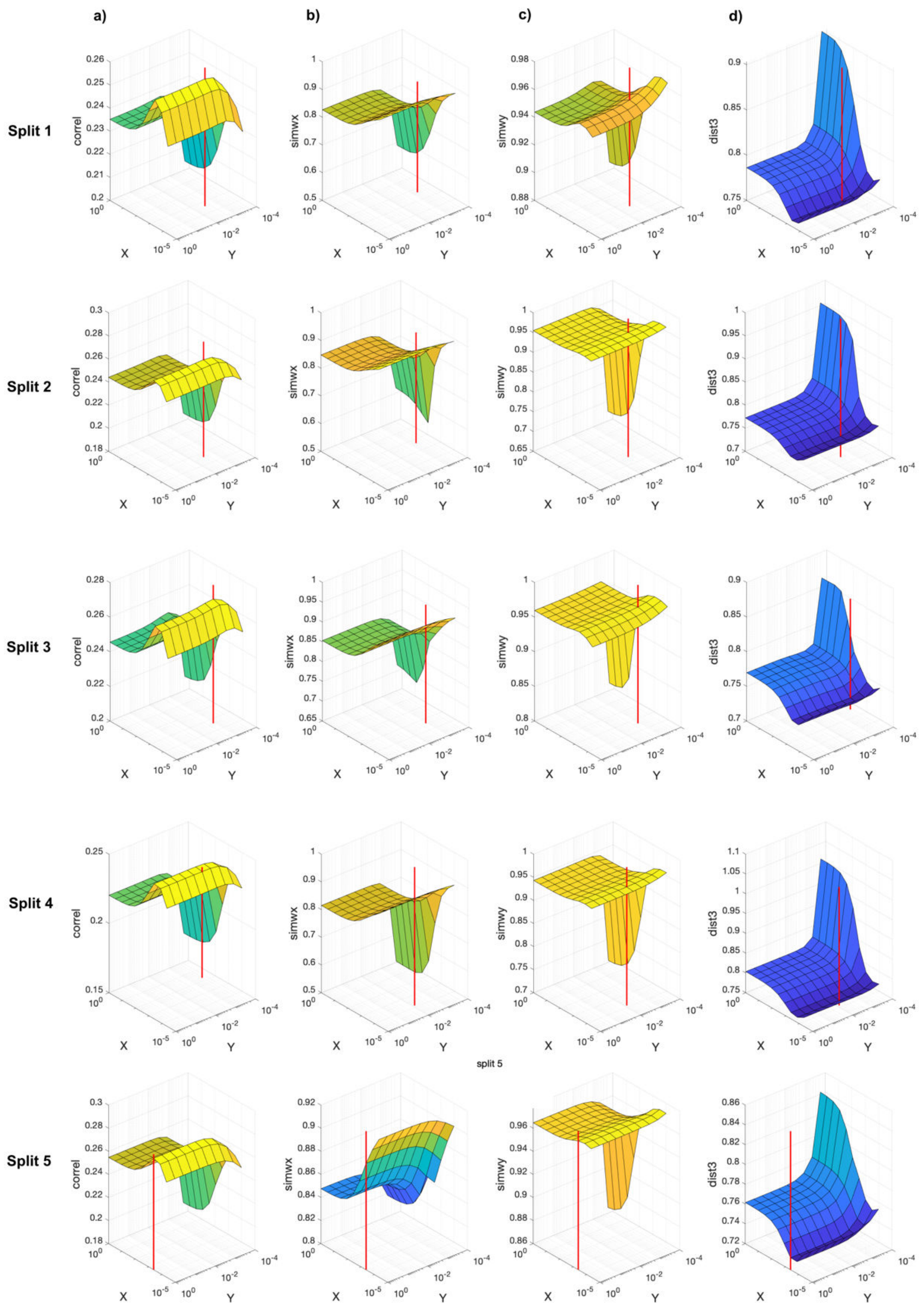

**Figure S27. Model optimization for the latent dimension of physical robustness for raw grey matter volume in the main sample.** The red line indicates the selected model. The z axis represents the test canonical correlation (column a), the similarity of weights in grey matter volume (column b) and risk factors (column c), and the joint generalizability-stability criteria (column d). The x and y axes represent 1-minus the hyperparameters tested for grey matter volume and risk factors, respectively (1-hyperparameter). The x and y axes are shown in logarithmic scale.

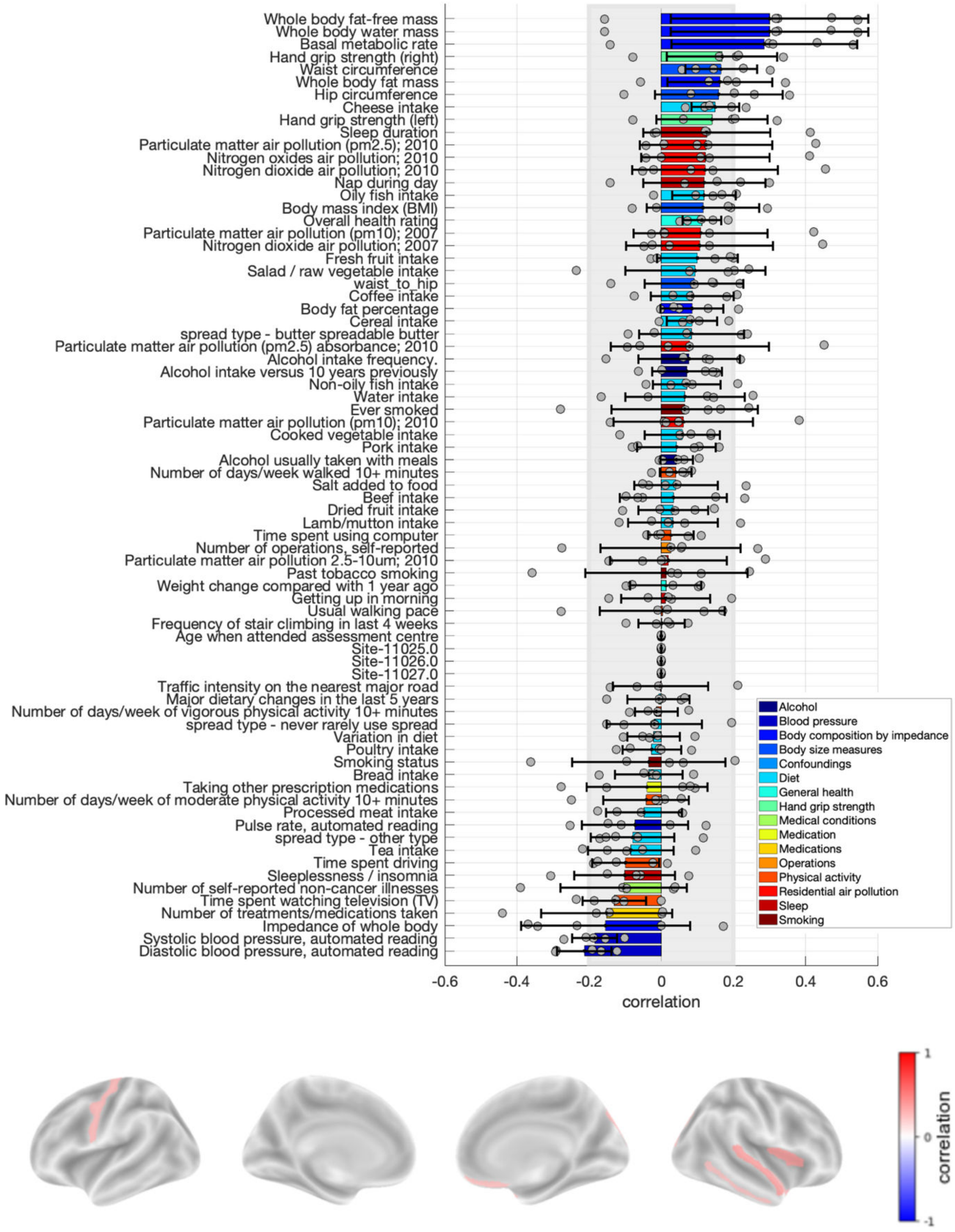

**Figure S28. Loadings of the latent dimension of physical robustness for raw cortical thickness in women.** Risk factors loadings and brain loadings. Shown loadings represent the average over the five outer splits. Error bars depict one standard deviation. The shadowed zone marks loadings between −0.2 and 0.2. Note that this latent dimension was significant only in one split out of five.

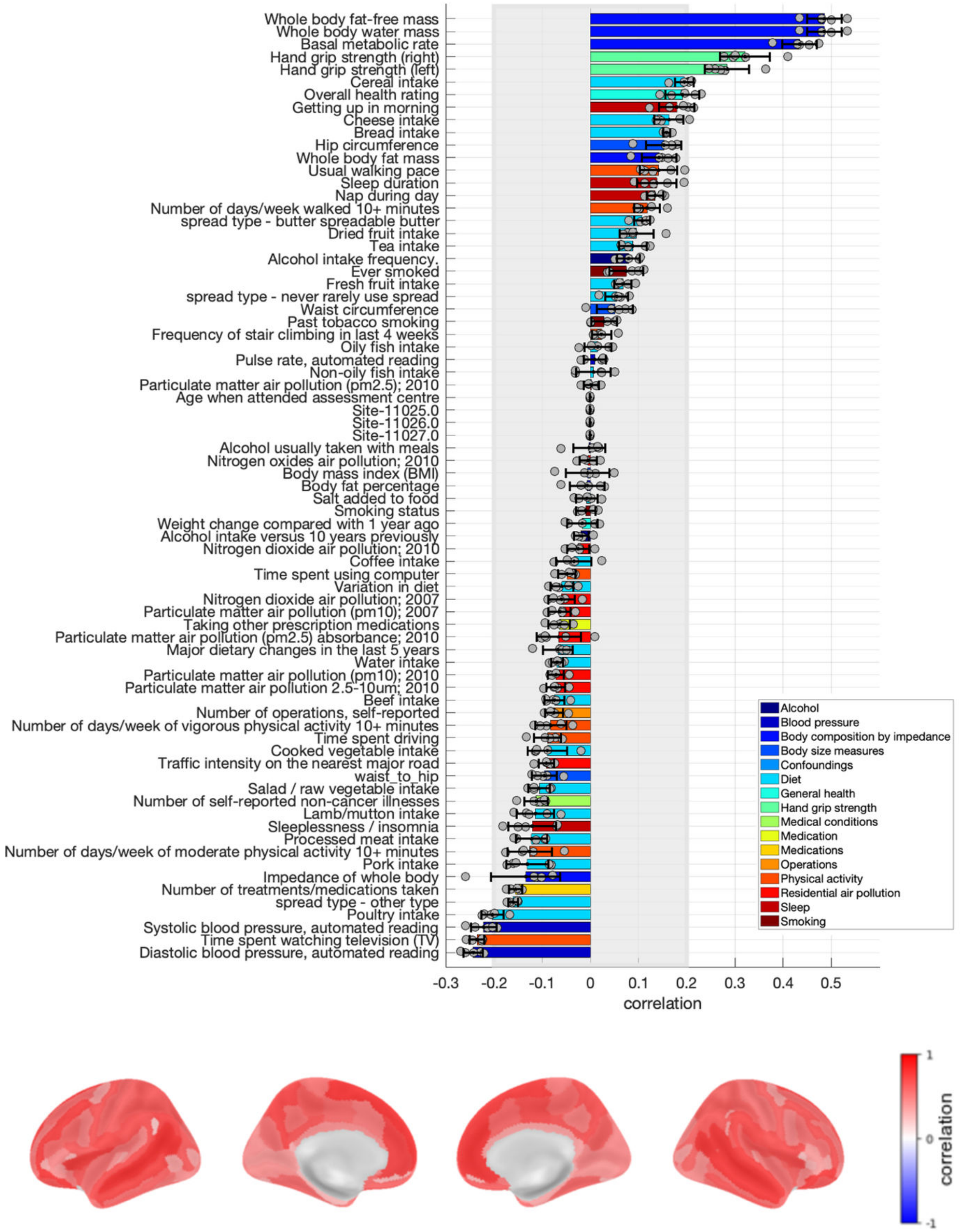

**Figure S29. Loadings of the latent dimension of physical robustness for raw grey matter volume in women.** Risk factors loadings and brain loadings. Shown loadings represent the average over the five outer splits. Error bars depict one standard deviation. The shadowed zone marks loadings between −0.2 and 0.2.

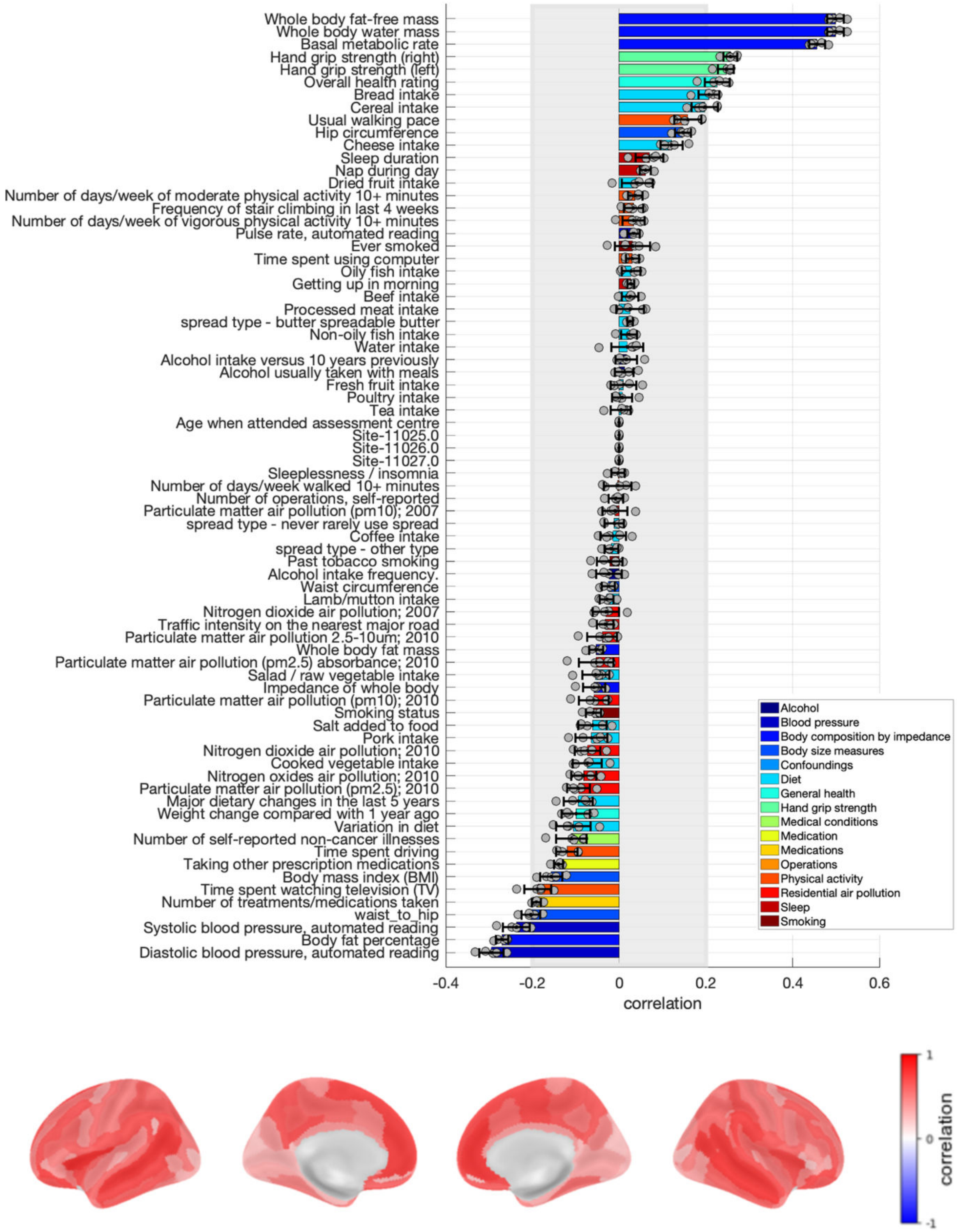

**Figure S30. Loadings of the latent dimension of physical robustness for raw grey matter volume in men.** Risk factors loadings and brain loadings. Shown loadings represent the average over the five outer splits. Error bars depict one standard deviation. The shadowed zone marks loadings between −0.2 and 0.2.

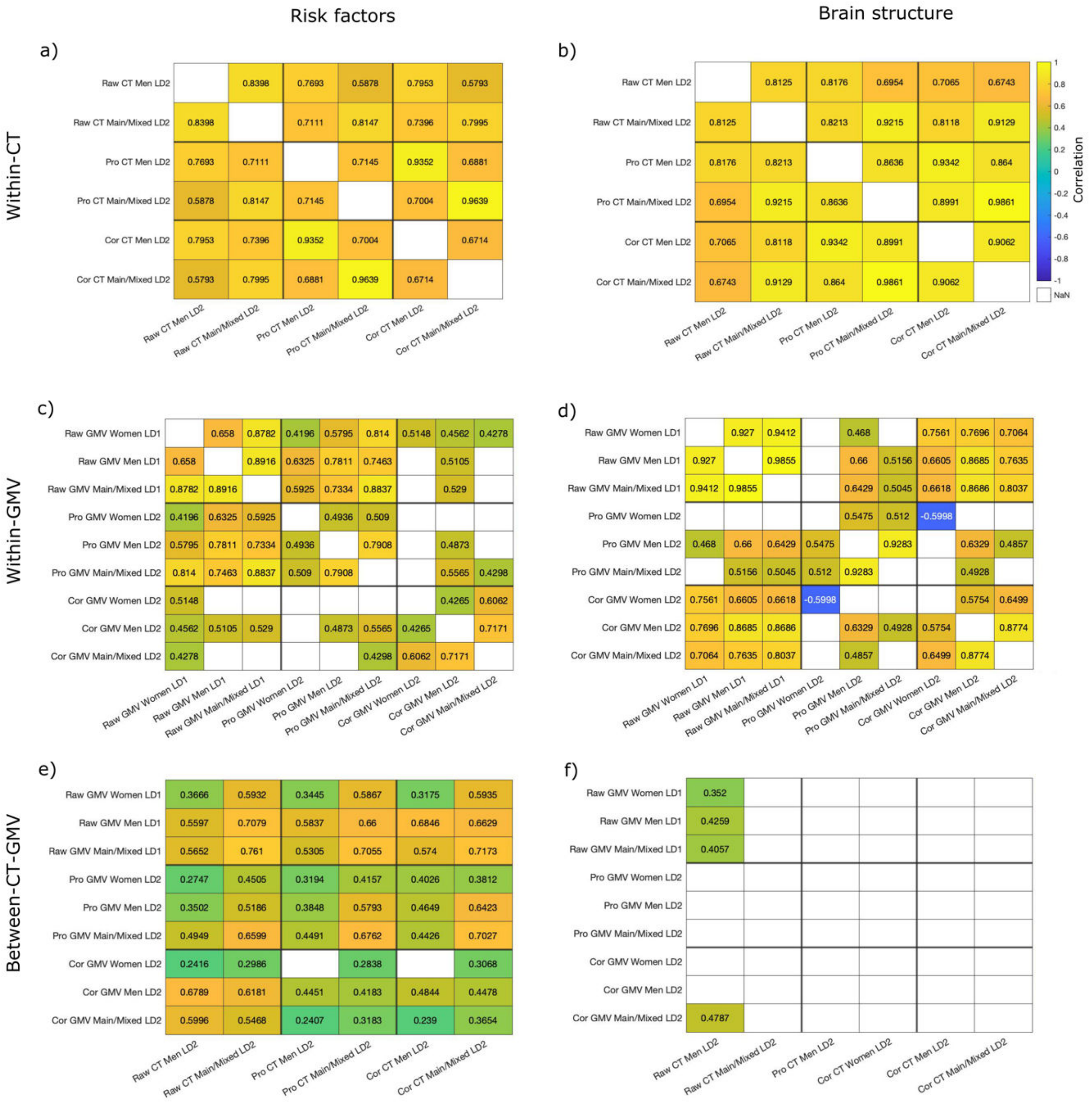

**Figure S31. Comparison of risk factor loadings and brain loadings for the physical robustness latent dimension across samples and brain structural measures.** The first row shows the Spearman correlation of a) risk factors loadings and b) brain loadings for the latent dimension of physical robustness yielded with cortical thickness. The second row shows the correlation of c) risk factors loadings and d) brain loadings for the latent dimension of physical robustness yielded with grey matter volume. Only comparisons that were significant after Bonferroni correction are shown.

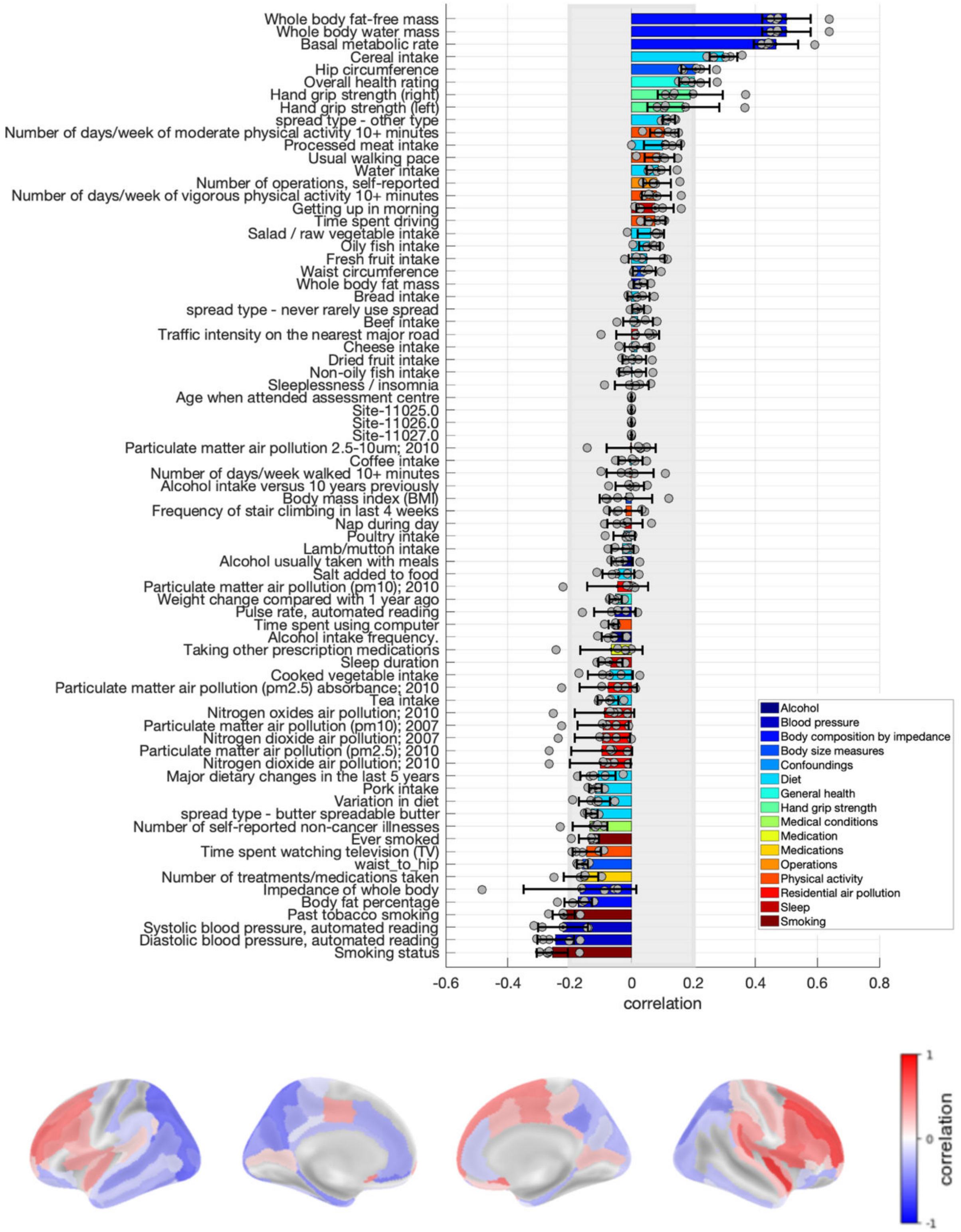

**Figure S32. Loadings of the latent dimension of physical robustness for proportional cortical thickness in men.** Risk factors loadings and brain loadings. Shown loadings represent the average over the five outer splits. Error bars depict one standard deviation. The shadowed zone marks loadings between −0.2 and 0.2.

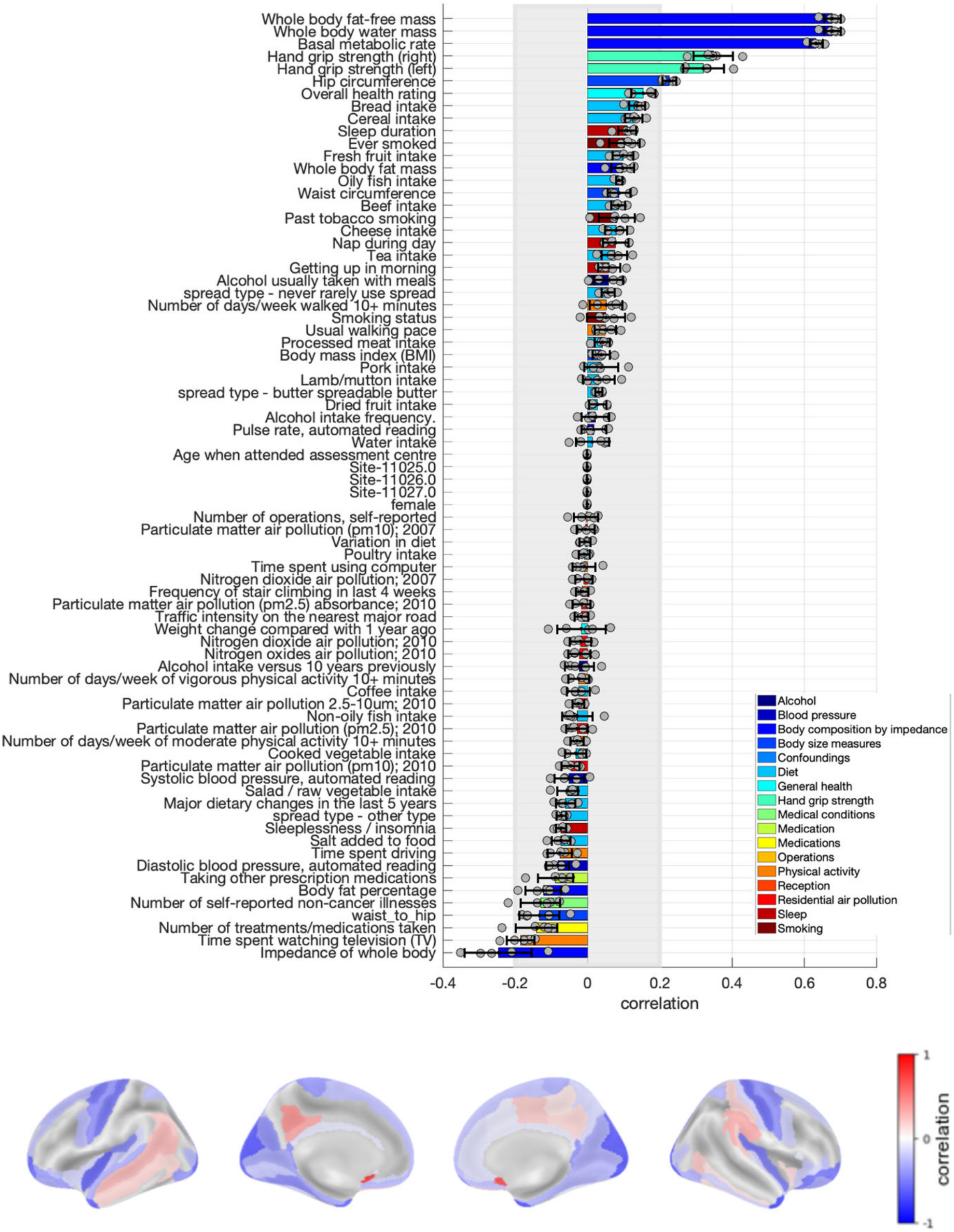

**Figure S33. Loadings of the latent dimension of physical robustness for proportional grey matter volume in the main sample.** Risk factors loadings and brain loadings. Shown loadings represent the average over the five outer splits. Error bars depict one standard deviation. The shadowed zone marks loadings between −0.2 and 0.2.

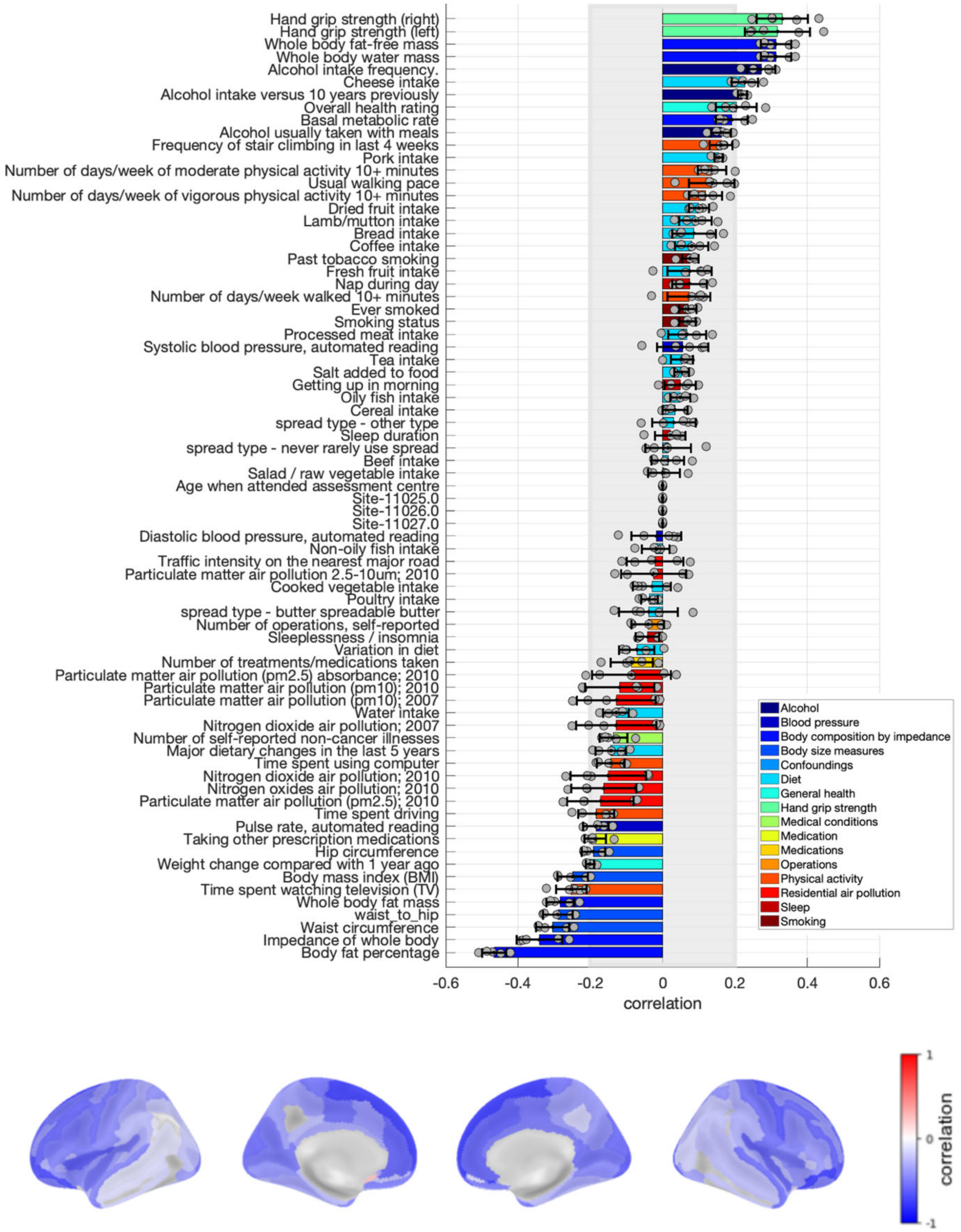

**Figure S34. Loadings of the latent dimension of physical robustness for proportional grey matter volume in women.** Risk factors loadings and brain loadings. Shown loadings represent the average over the five outer splits. Error bars depict one standard deviation. The shadowed zone marks loadings between −0.2 and 0.2.

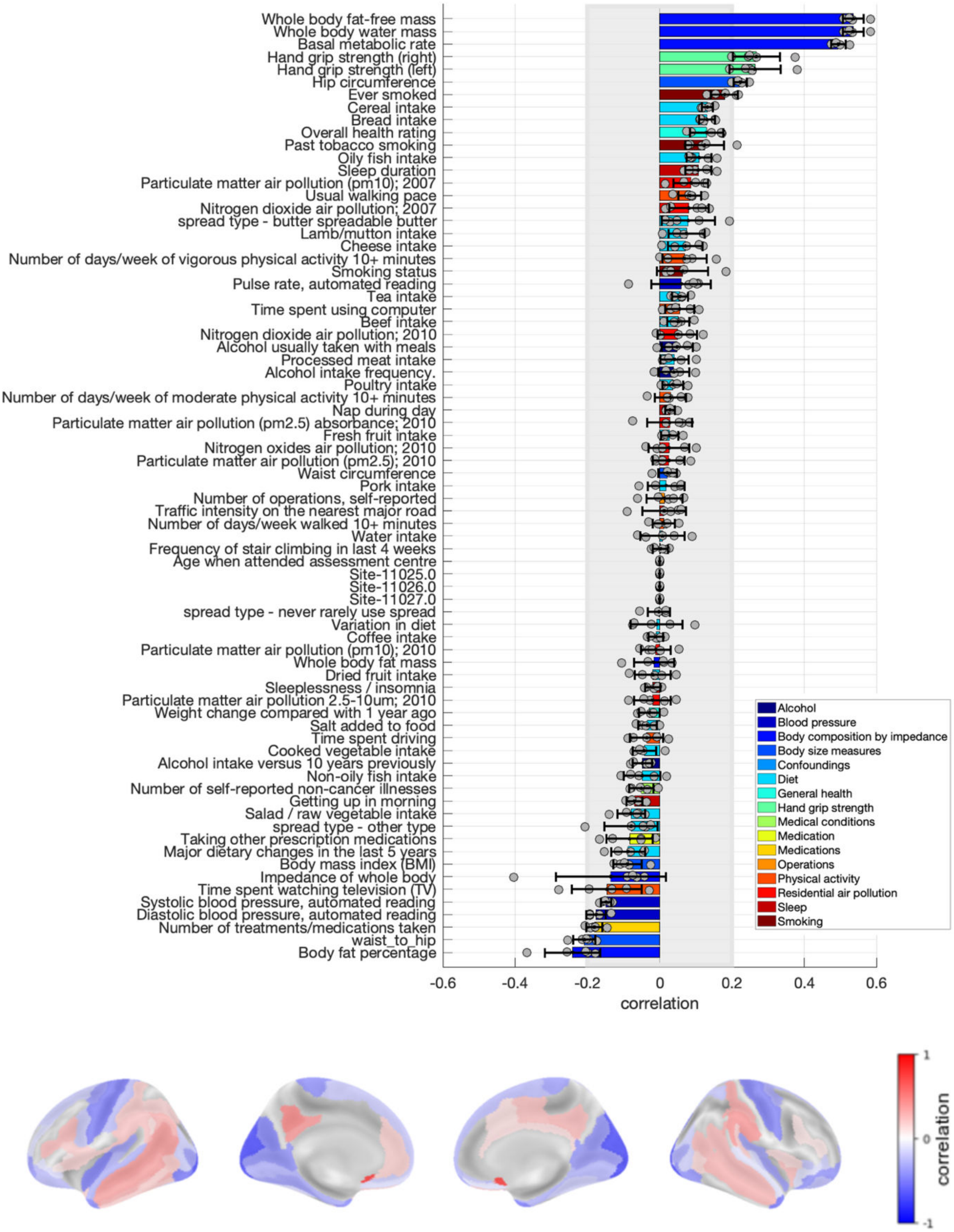

**Figure S35. Loadings of the latent dimension of physical robustness for proportional grey matter volume in men.** Risk factors loadings and brain loadings. Shown loadings represent the average over the five outer splits. Error bars depict one standard deviation. The shadowed zone marks loadings between −0.2 and 0.2.

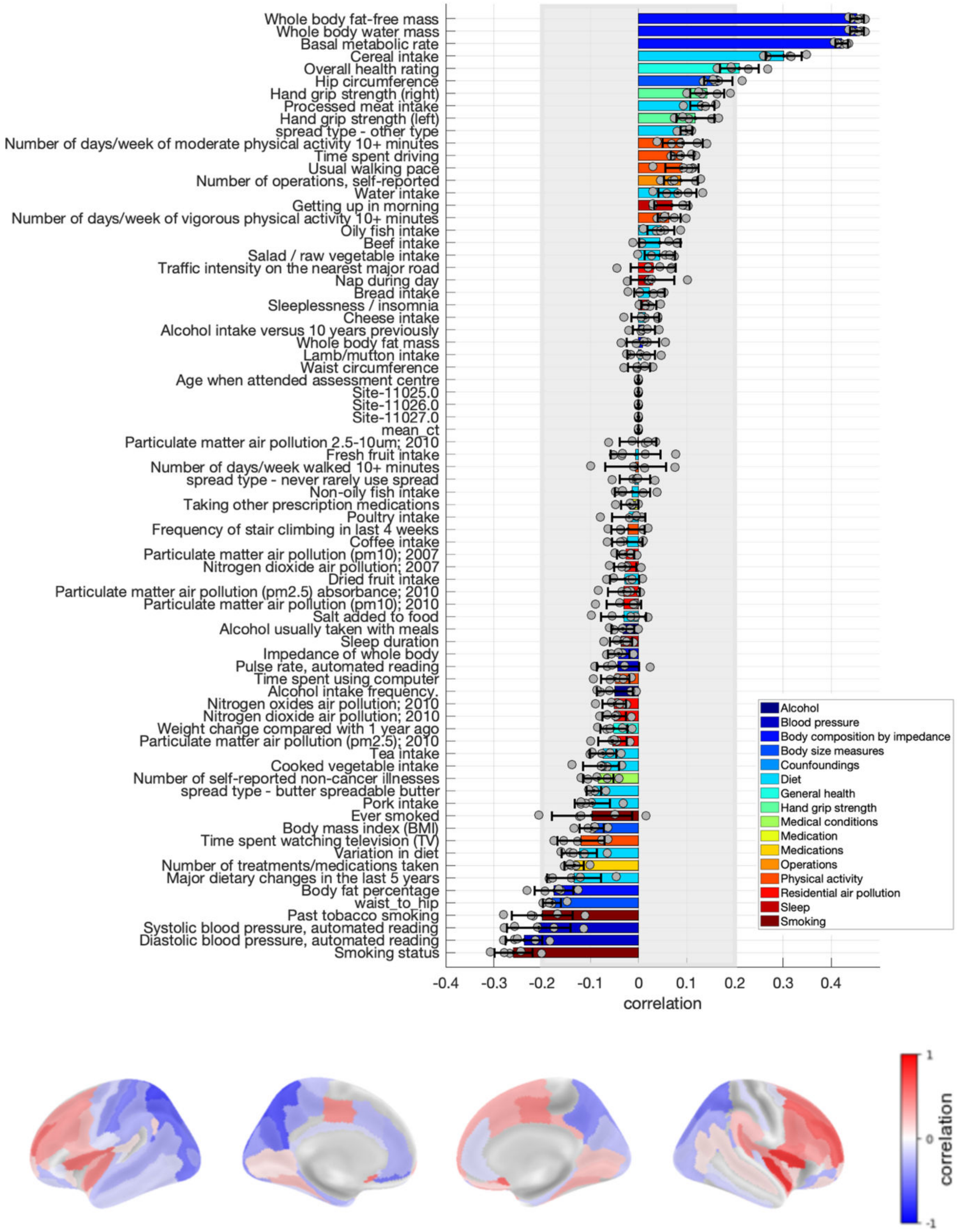

**Figure S36. Loadings of the latent dimension of physical robustness for corrected cortical thickness in men.** Risk factors loadings and brain loadings. Shown loadings represent the average over the five outer splits. Error bars depict one standard deviation. The shadowed zone marks loadings between −0.2 and 0.2.

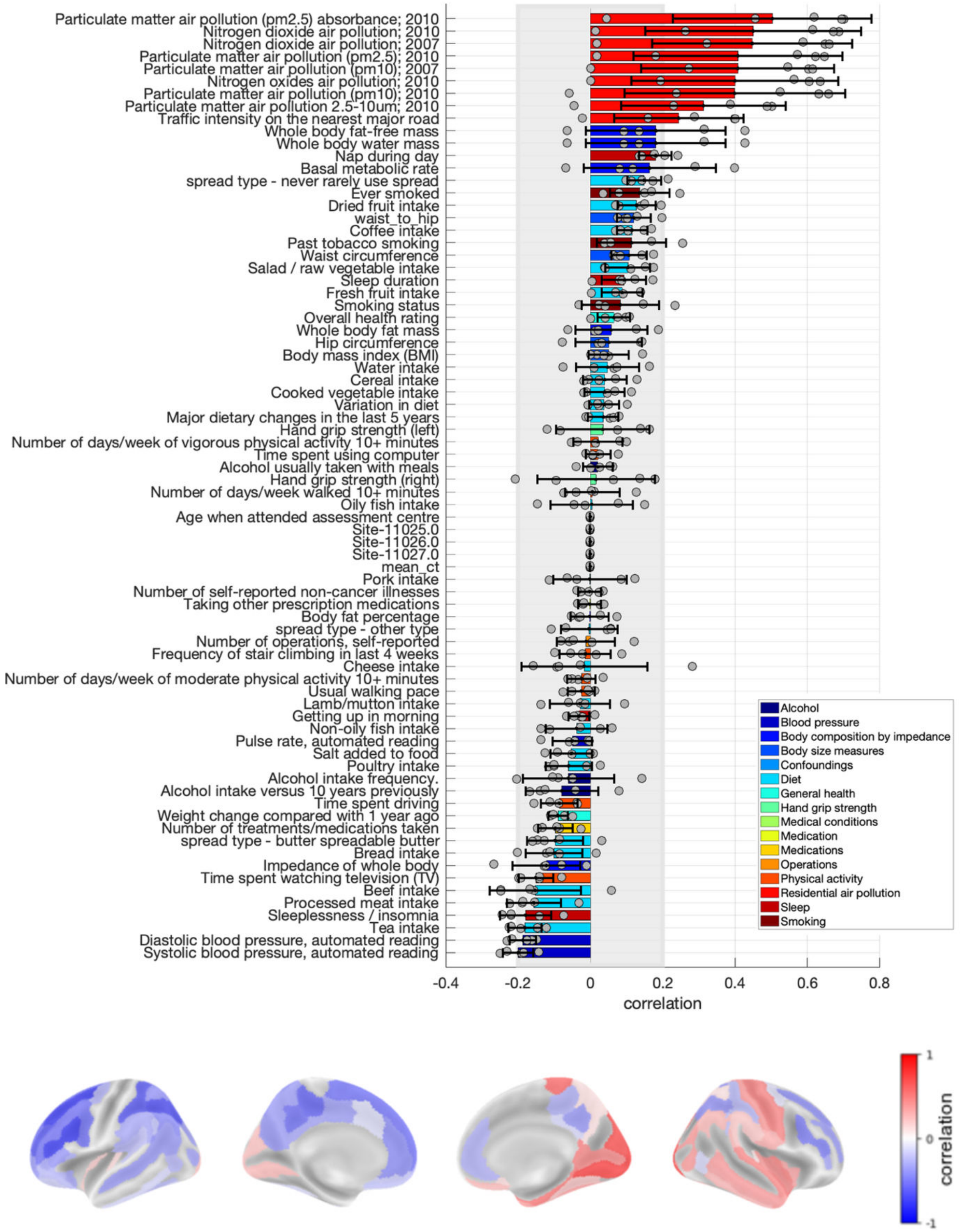

**Figure S37. Loadings of the latent dimension of physical robustness for corrected cortical thickness in women.** Risk factors loadings and brain loadings. Shown loadings represent the average over the five outer splits. Error bars depict one standard deviation. The shadowed zone marks loadings between −0.2 and 0.2. Note that this latent dimension was significant only in one split out of five.

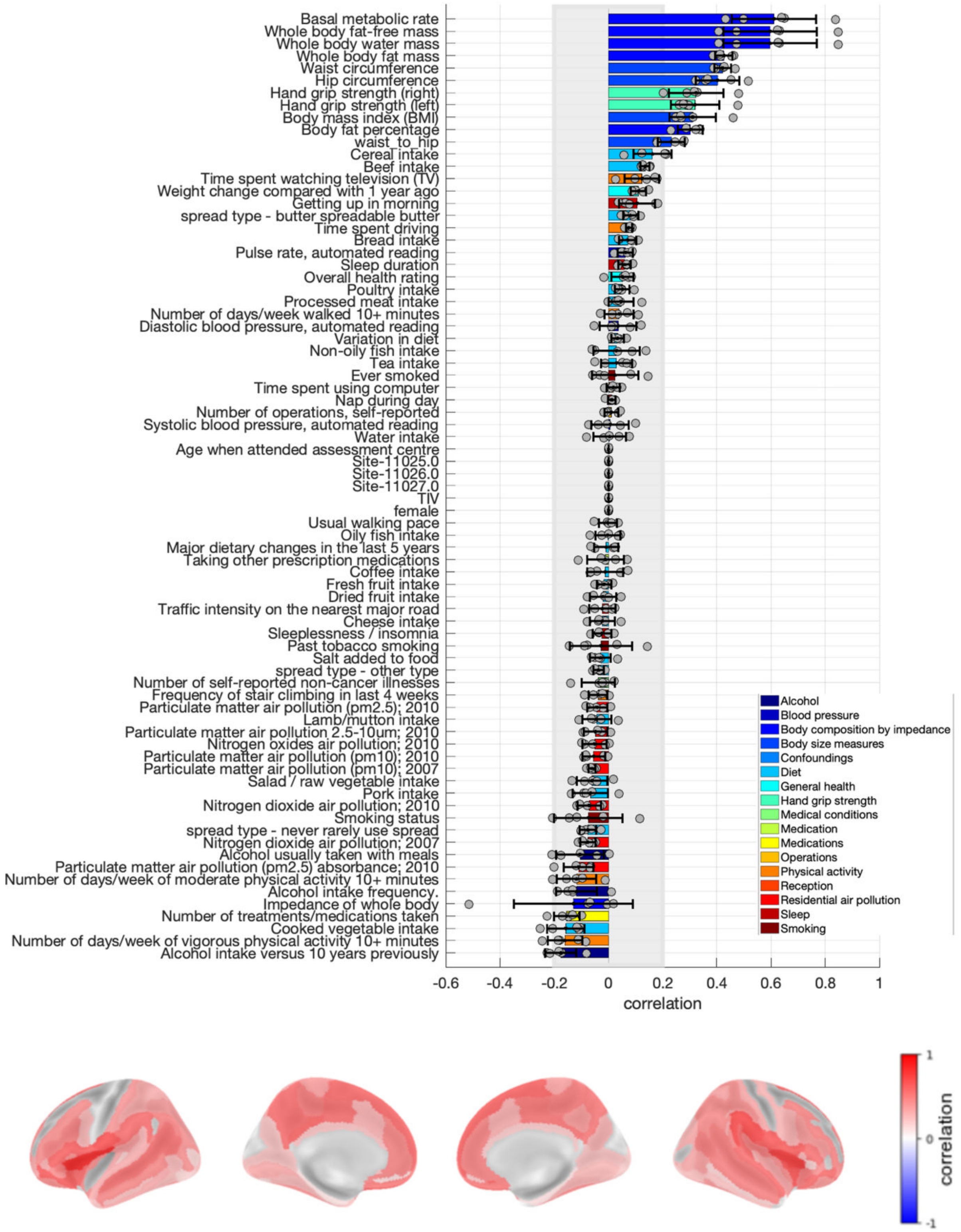

**Figure S38. Loadings of the latent dimension of physical robustness for corrected grey matter volume in the main sample.** Risk factors loadings and brain loadings. Shown loadings represent the average over the five outer splits. Error bars depict one standard deviation. The shadowed zone marks loadings between −0.2 and 0.2.

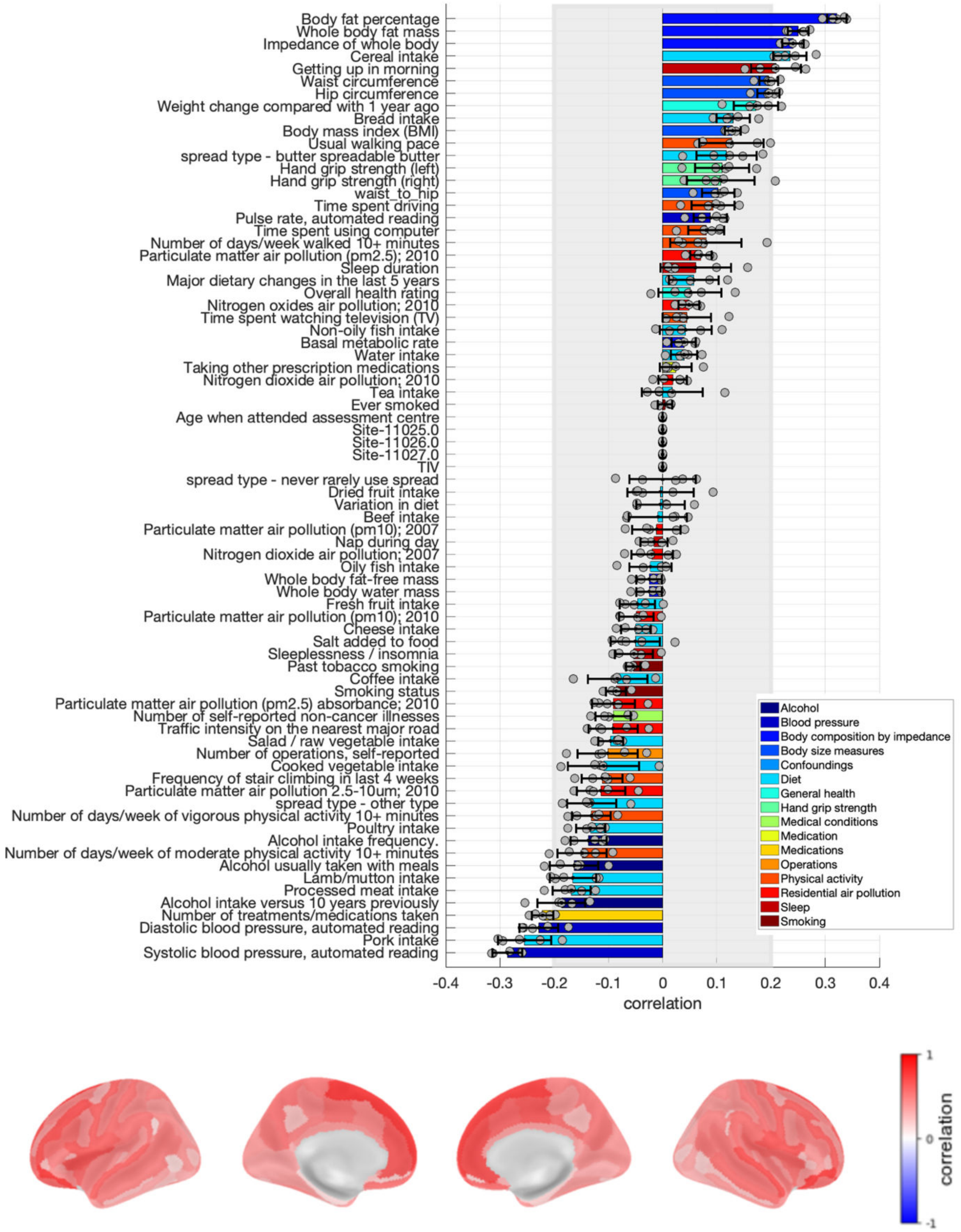

**Figure S39. Loadings of the latent dimension of physical robustness for corrected grey matter volume in women.** Risk factors loadings and brain loadings. Shown loadings represent the average over the five outer splits. Error bars depict one standard deviation. The shadowed zone marks loadings between −0.2 and 0.2.

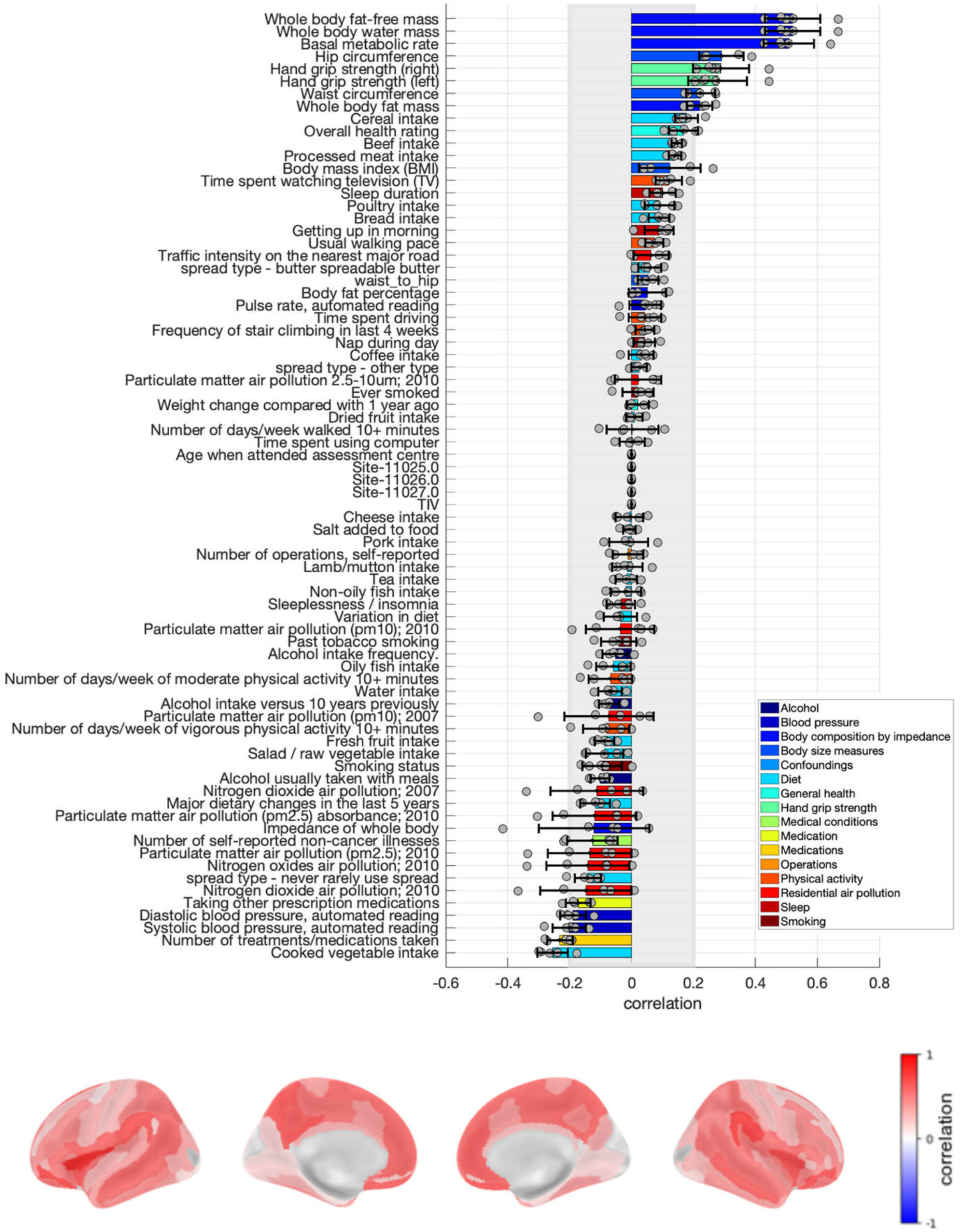

**Figure S40. Loadings of the latent dimension of physical robustness for corrected grey matter volume in men.** Risk factors loadings and brain loadings. Shown loadings represent the average over the five outer splits. Error bars depict one standard deviation. The shadowed zone marks loadings between −0.2 and 0.2.

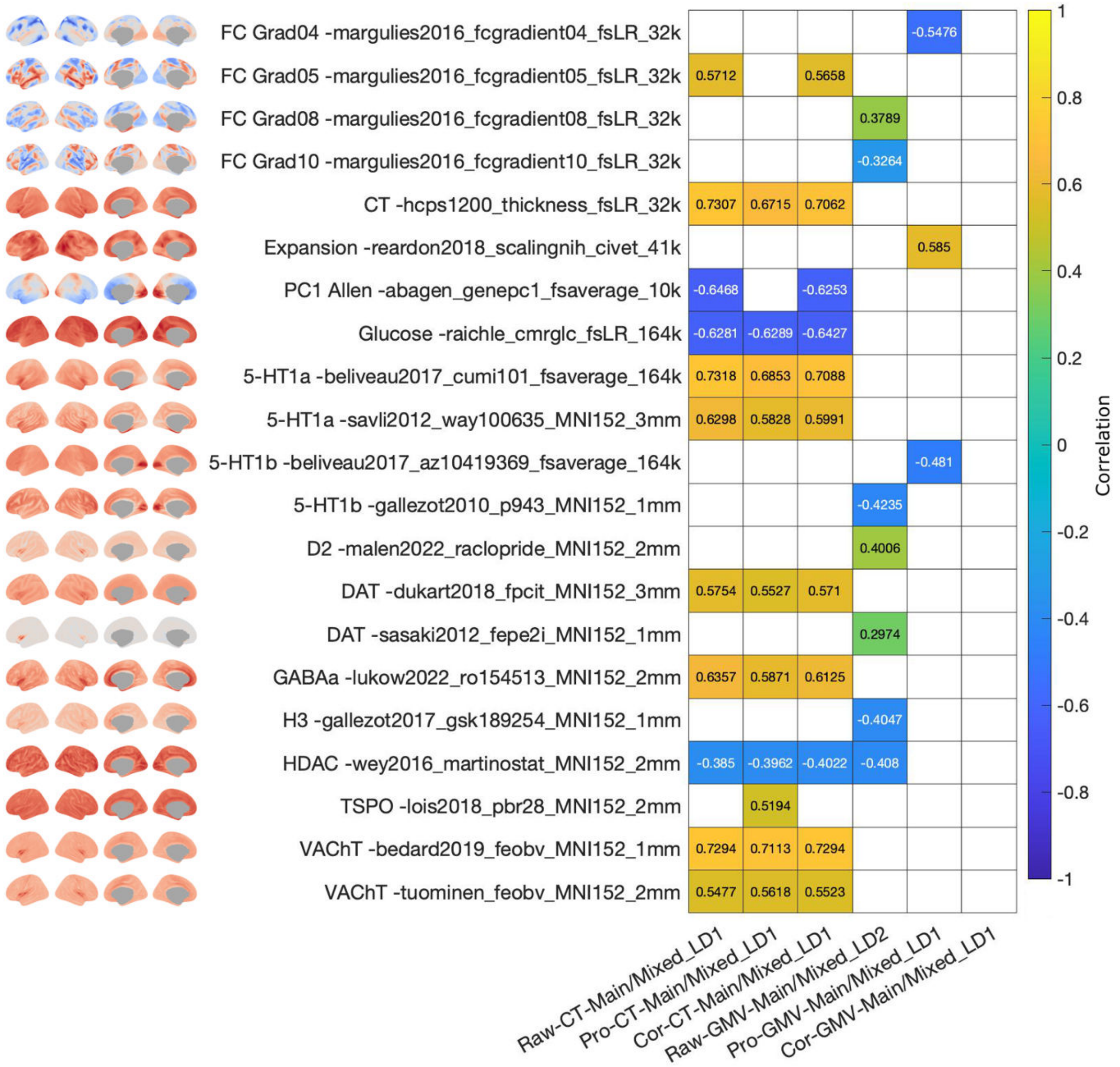

Figure S41. Association of brain structural loadings with brain maps for the latent dimension of cardiometabolic health. The association between the brain pattern of the latent dimension and brain maps was assessed with spin test. Only data for brain maps that yielded a significant association with at least one map of loadings are shown. White tiles represent non-significant associations.

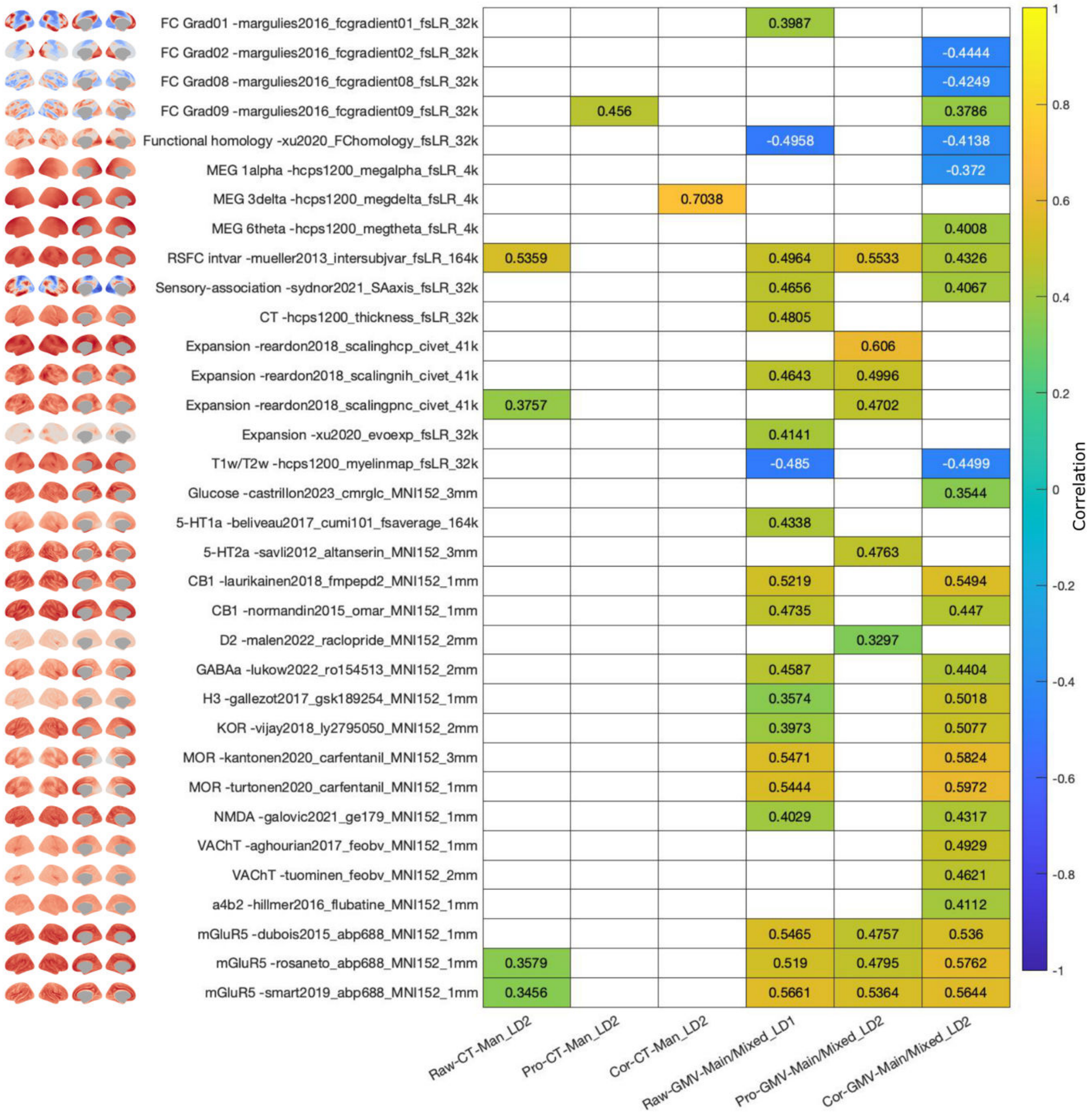

**Figure S42. Association of brain structural loadings with brain maps for the latent dimension of physical robustness.** The association between the brain pattern of the latent dimension and brain maps was assessed with spin test. Only data for brain maps that yielded a significant association with at least one map of loadings are shown. White tiles represent non-significant associations.

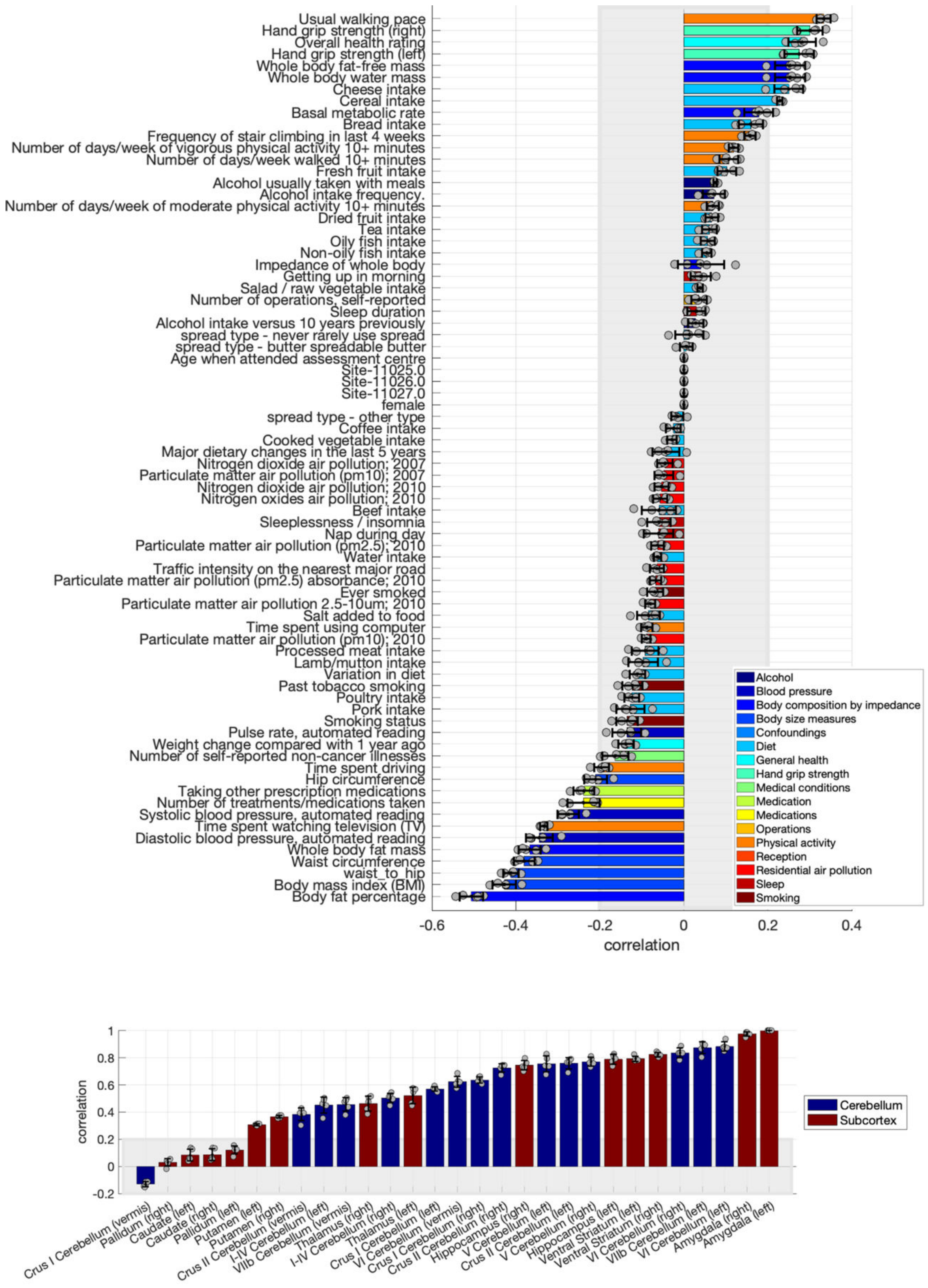

**Figure S43. Loadings of the first latent dimension linking risk factors to subcortical and cerebellar volumes.** Risk factors loadings and brain loadings. Shown loadings represent the average over the five outer splits. Error bars depict one standard deviation. The shadowed zone marks loadings between −0.2 and 0.2.

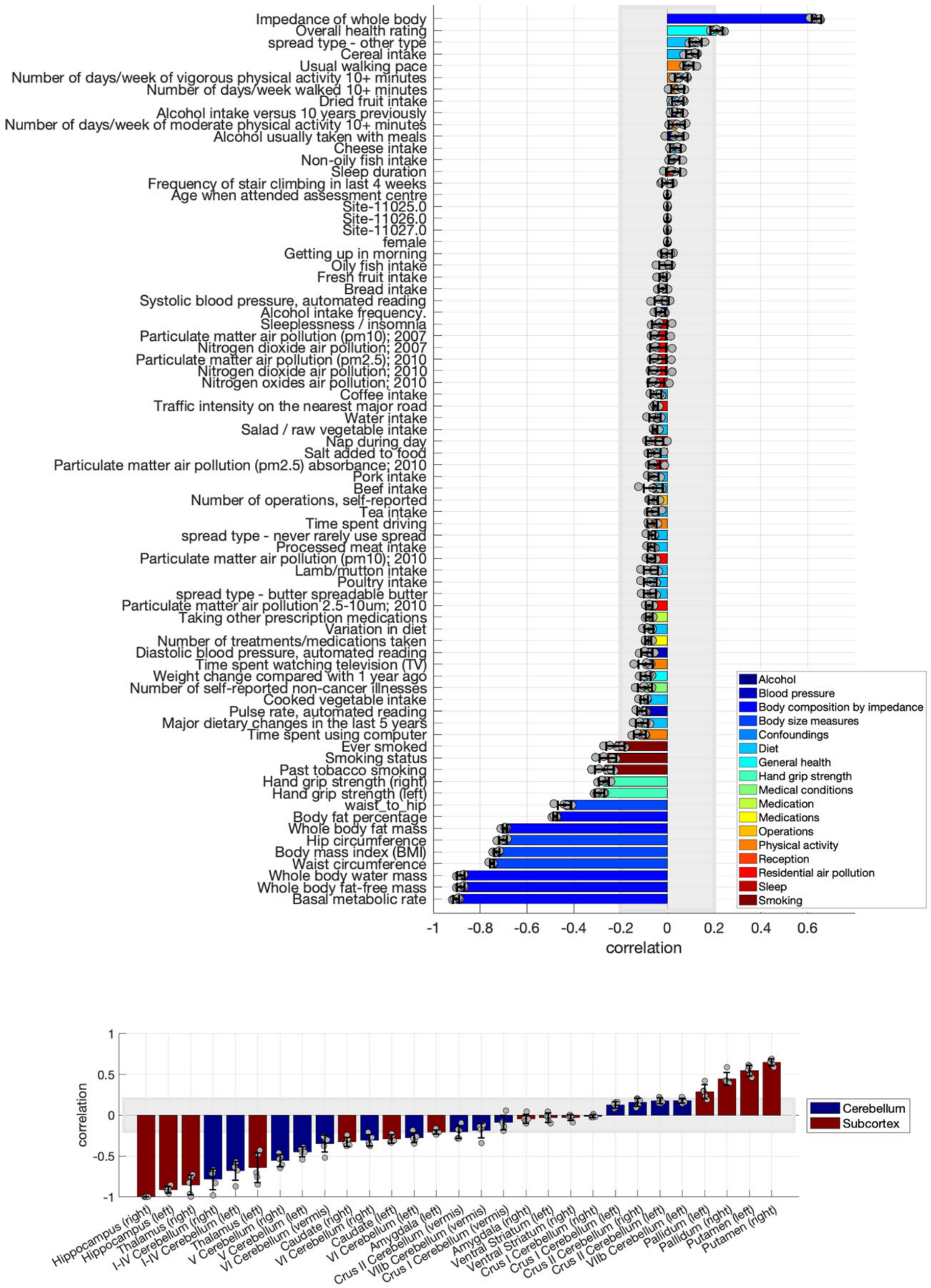

**Figure S44. Loadings of the second latent dimension linking risk factors to subcortical and cerebellar volumes.** Risk factors loadings and brain loadings. Shown loadings represent the average over the five outer splits. Error bars depict one standard deviation. The shadowed zone marks loadings between −0.2 and 0.2.

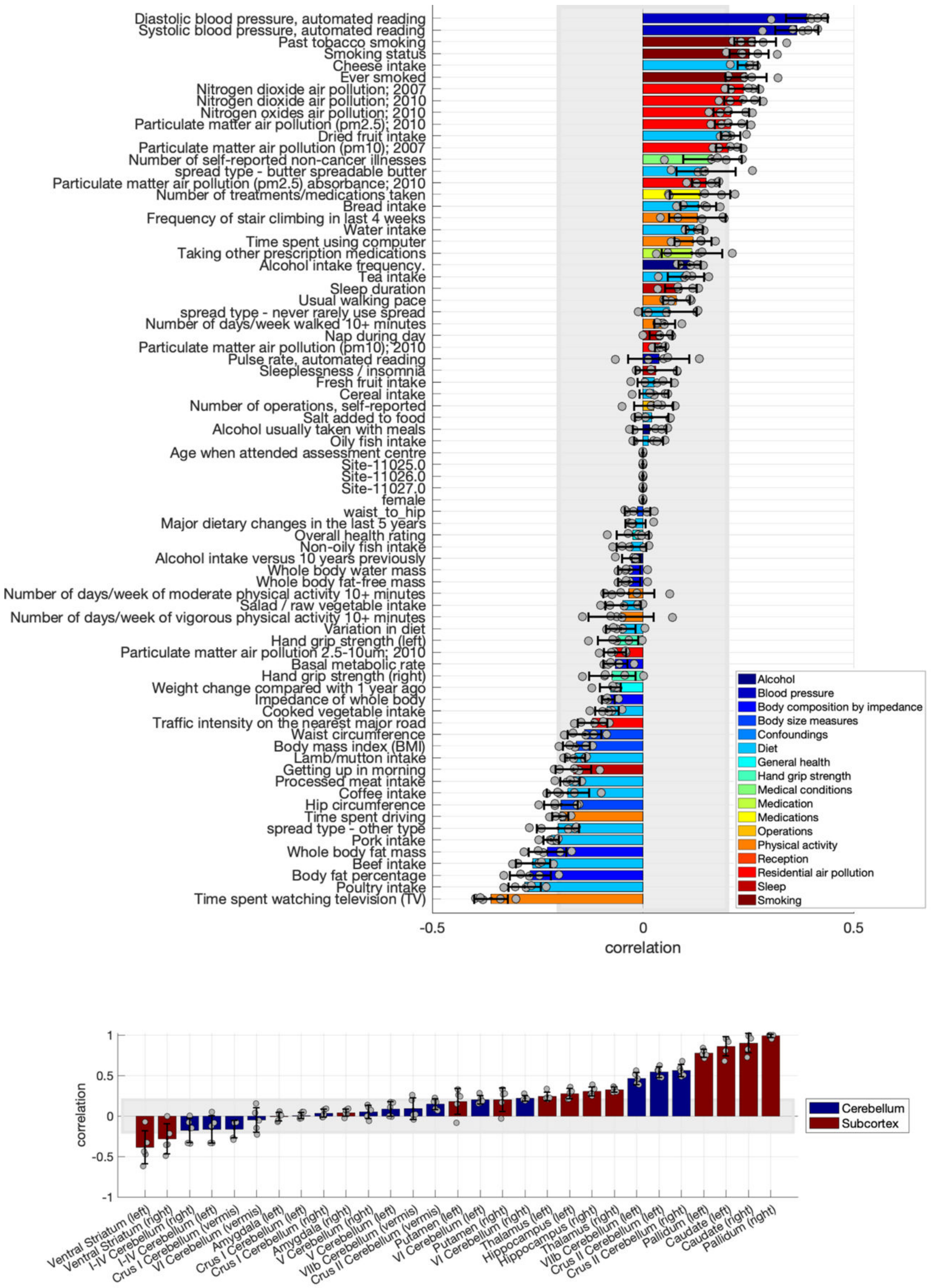

**Figure S45. Loadings of the third latent dimension linking risk factors to subcortical and cerebellar volumes.** Risk factors loadings and brain loadings. Shown loadings represent the average over the five outer splits. Error bars depict one standard deviation. The shadowed zone marks loadings between −0.2 and 0.2.

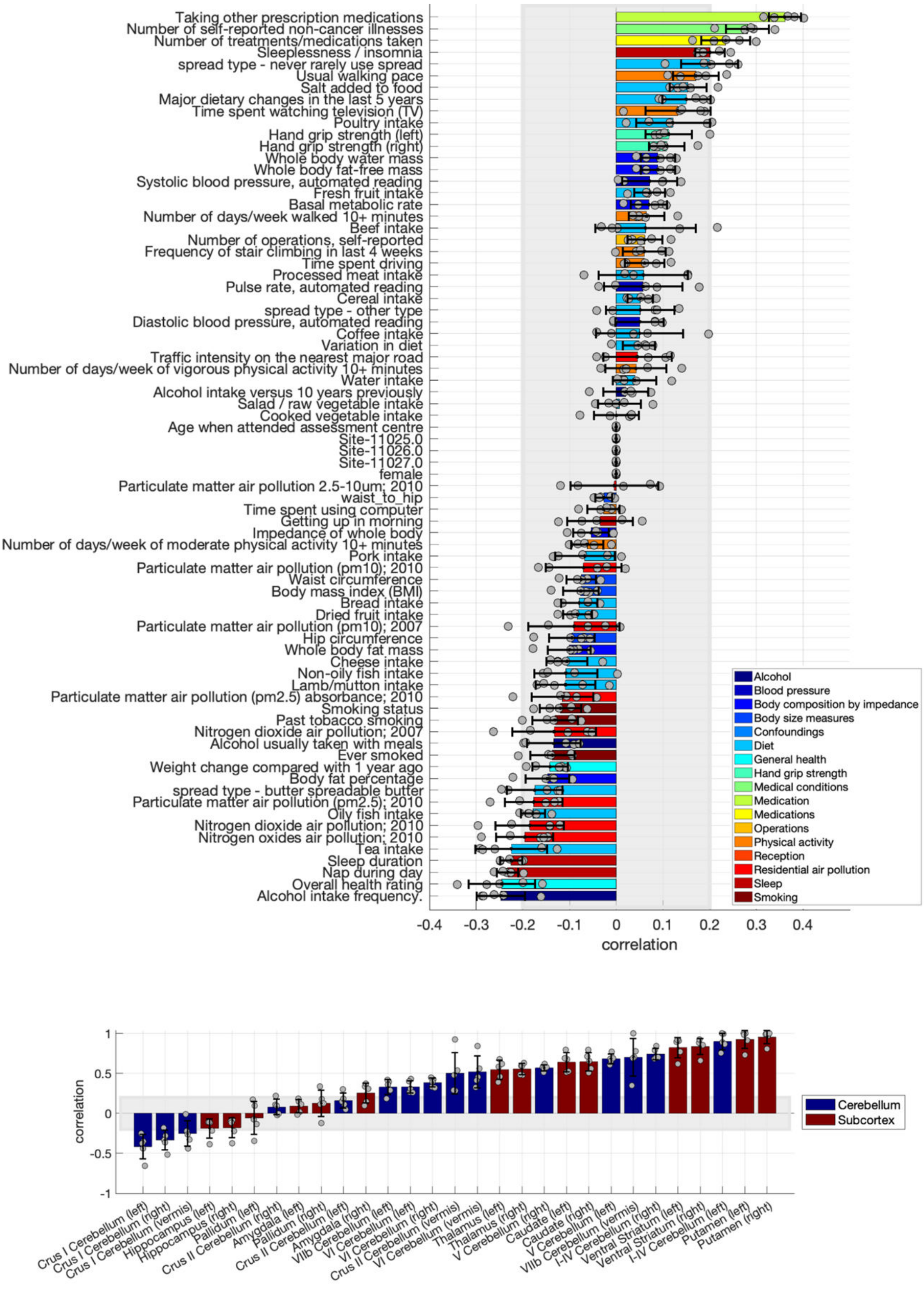

**Figure S46. Loadings of the fourth latent dimension linking risk factors to subcortical and cerebellar volumes.** Risk factors loadings and brain loadings. Shown loadings represent the average over the five outer splits. Error bars depict one standard deviation. The shadowed zone marks loadings between −0.2 and 0.2.

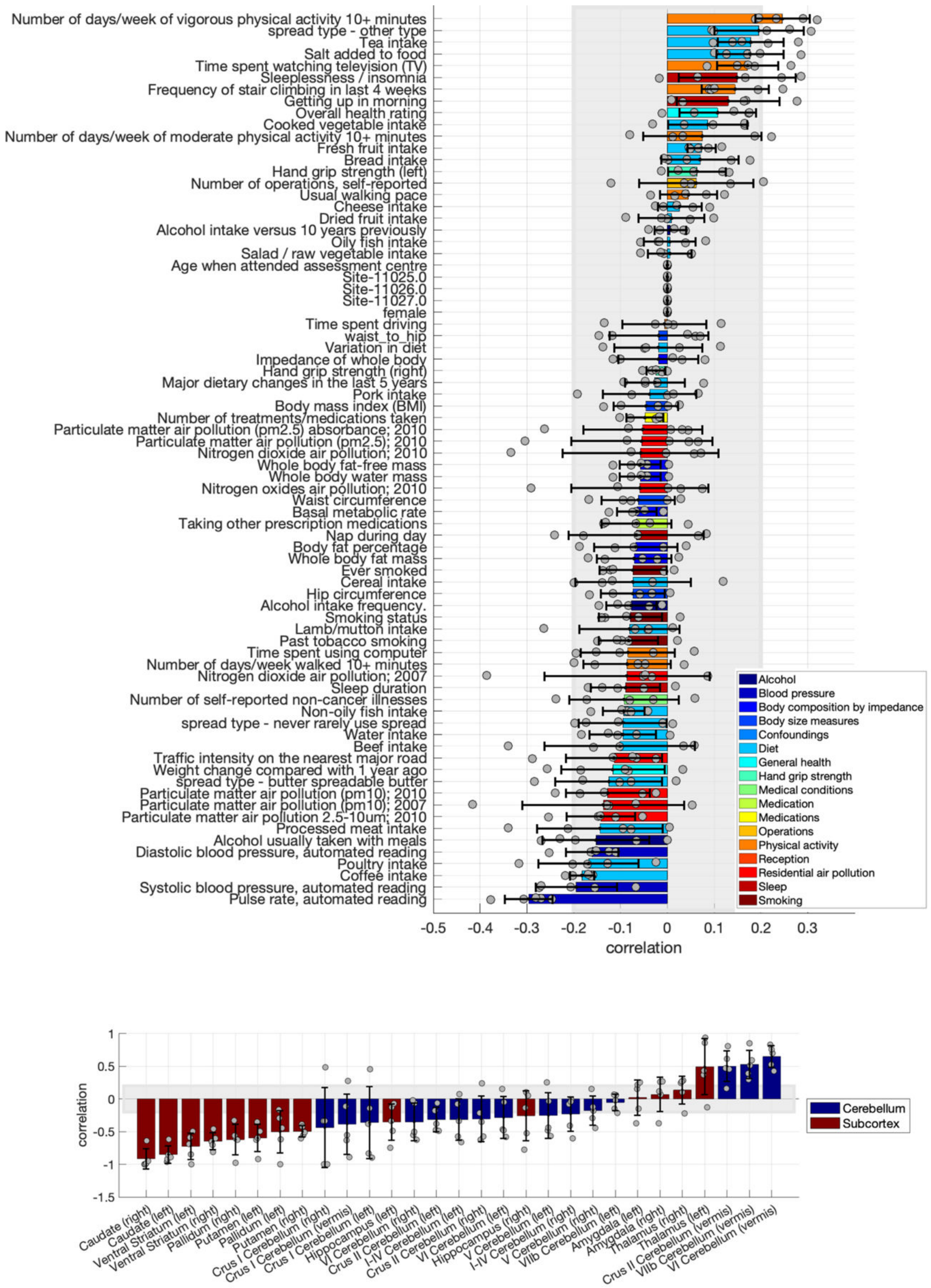

**Figure S47. Loadings of the fifth latent dimension linking risk factors to subcortical and cerebellar volumes.** Risk factors loadings and brain loadings. Shown loadings represent the average over the five outer splits. Error bars depict one standard deviation. The shadowed zone marks loadings between −0.2 and 0.2.

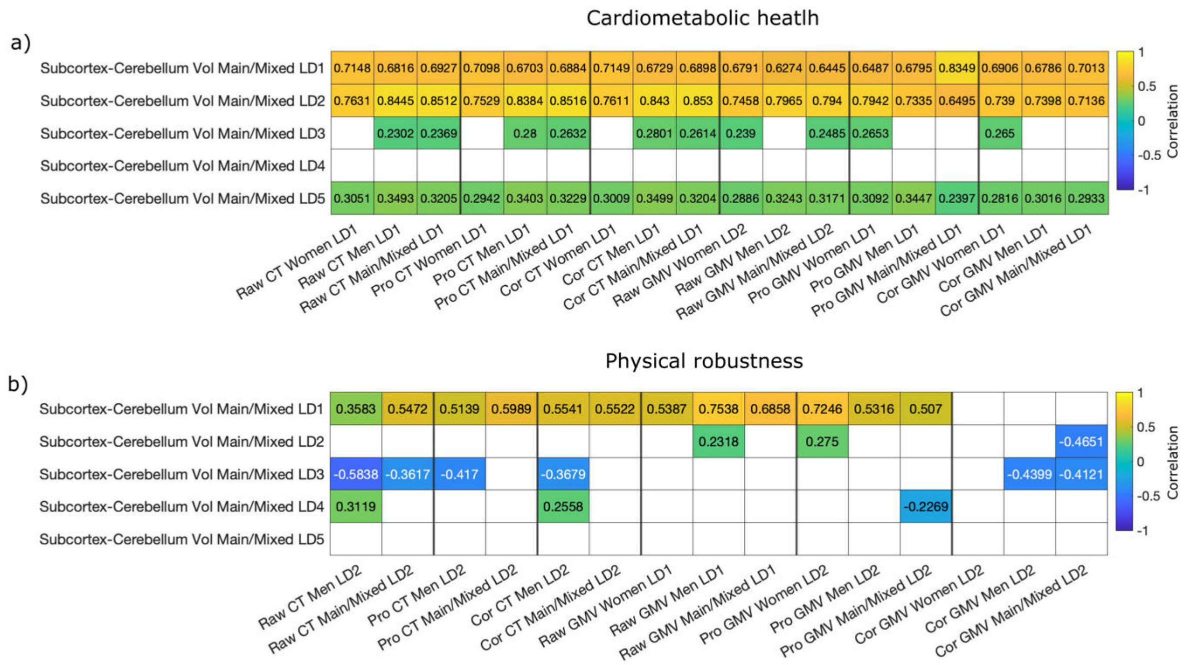

**Figure S48. Comparison of risk factor loadings between latent dimensions yielded with subcortex-cerebellum and with CT and GMV in cortex.** Panel a shows associations of risk factors for the latent dimension of cardiometabolic health. Panel b shows associations of risk factors for the latent dimension of physical robustness. Only associations that remain significant after Bonferroni correction are shown. White tiles represent non-significant associations.

### 5. Bibliography

1. Kim SE, Lee JS, Woo S, Kim S, Kim HJ, Park S, et al. Sex-specific relationship of cardiometabolic syndrome with lower cortical thickness. Neurology. 2019;93(11):e1045–57.

2. Miller AA, Spencer SJ. Obesity and neuroinflammation: A pathway to cognitive impairment. Brain, Behavior, and Immunity. 2014 Nov;42:10–21.

3. Dresser R. Wanted single, white male for medical research. The Hastings Center Report. 1992;22(1):24–9.

4. Hodes GE, Kropp DR. Sex as a biological variable in stress and mood disorder research. Nature Mental Health. 2023;1(7):453–61.

5. Beery AK, Zucker I. Sex bias in neuroscience and biomedical research. Neuroscience & Biobehavioral Reviews. 2011;35(3):565–72.

6. Alfaro-Almagro F, Jenkinson M, Bangerter NK, Andersson JLR, Sotiropoulos SN, Jbabdi S, et al. Image processing and Quality Control for the first 10,000 brain imaging datasets from UK Biobank. NeuroImage. 2018;166(October 2017):400–24.

7. Sasse L, Nicolaisen-Sobesky E, Dukart J, Eickhoff SB, Götz M, Hamdan S, et al. Overview of leakage scenarios in supervised machine learning. J Big Data. 2025 May 29;12(1):135.
